# Di-berberine conjugates as chemical probes of *Pseudomonas aeruginosa* MexXY-OprM efflux function and inhibition

**DOI:** 10.1101/2023.03.24.533986

**Authors:** Logan G. Kavanaugh, Andrew R. Mahoney, Debayan Dey, William M. Wuest, Graeme L. Conn

**Affiliations:** Department of Biochemistry, Emory University School of Medicine, Atlanta, GA; Department of Chemistry, Emory University, Atlanta, GA; Emory Antibiotic Resistance Center, Emory University, Atlanta, GA

**Keywords:** *Pseudomonas*, antibiotic, efflux, Resistance-Nodulation-Division (RND), berberine, efflux pump inhibitor, MexXY-OprM

## Abstract

The Resistance-Nodulation-Division (RND) efflux pump superfamily is pervasive among Gram-negative pathogens and contributes extensively to clinical antibiotic resistance. The opportunistic pathogen *Pseudomonas aeruginosa* contains 12 RND-type efflux systems, with four contributing to resistance including MexXY-OprM which is uniquely able to export aminoglycosides. At the site of initial substrate recognition, small molecule probes of the inner membrane transporter (e.g., MexY) have potential as important functional tools to understand substrate selectivity and a foundation for developing adjuvant efflux pump inhibitors (EPIs). Here, we optimized the scaffold of berberine, a known but weak MexY EPI, using an *in-silico* high-throughput screen to identify di-berberine conjugates with enhanced synergistic action with aminoglycosides. Further, docking and molecular dynamics simulations of di-berberine conjugates reveal unique contact residues and thus sensitivities of MexY from distinct *P. aeruginosa* strains. This work thereby reveals di-berberine conjugates to be useful probes of MexY transporter function and potential leads for EPI development.

## Introduction

Multi-drug resistant bacteria pose a global health threat with escalating infection cases resulting in increased morbidity and mortality, longer hospital stays, and increased financial burden^1,2^. In patients with cystic fibrosis, chronic lung infections caused by *Pseudomonas aeruginosa* are common and eradication is challenging^3–5^. In *P. aeruginosa* and other Gram-negative pathogens, expression of the chromosomally encoded Resistance-Nodulation-Division (RND) superfamily efflux pumps contributes extensively to multidrug resistance^6–11^. However, development of efflux pump inhibitors (EPIs) as therapeutics has proven challenging and is further impacted by the more broadly slowed progression of new antibiotic discovery and development^12,13^.

*P. aeruginosa* encodes 12 hydrophobe/ amphiphile efflux-1 (HAE-1) family efflux pumps within the RND superfamily, of which four contribute to clinical antibiotic resistance: MexAB-OprM, MexCD-OprJ, MexEF-OprN, and MexXY-OprM^7–10^. These four pumps have overlapping, yet distinct antibiotic substrate profiles with MexXY-OprM having a unique ability to recognize and efflux aminoglycoside antibiotics^14–16^. For therapeutic regimes requiring aminoglycoside use, such as inhaled tobramycin for treatment of cystic fibrosis-associated *P. aeruginosa* infections, the MexXY-OprM efflux system plays a crucial role in resistance development and treatment failure^17–19^. A deeper understanding of the mechanisms governing antibiotic recognition by RND systems and the development of small molecule EPIs are urgently needed to negate efflux-mediated antibiotic resistance.

RND efflux pumps are tripartite protein complexes comprising a homotrimeric outer membrane channel (e.g., OprM), homohexameric periplasmic adaptor protein (e.g., MexX), and a homotrimeric inner membrane transporter (e.g., MexY; **Figure 1a**)^20–22^. The inner membrane transporter is a substrate/ H^+^ antiporter and is the initial site for substrate recognition. Each protomer of the trimeric inner membrane transporter contains 12 transmembrane helices (TM1-TM12) and a periplasmic domain with six subdomains that comprise the pore (PN1, PN2, PC1, PC2) and the MexX docking interface (DN, DC) (**Figure 1b**)^23–26^. Of the four predicted entry channels, two main predicted entry channels from the periplasmic space into the transporter are the cleft, between PC1 and PC2 of a single protomer, and the vestibule, between PN2 of one protomer and PC2 of a second protomer (**Figure 1b**)^18,24,27,28^. Each protomer cycles through three main conformational states that are required for substrate entry into the transporter and subsequent translocation to the channel of the periplasmic adapter: access, binding, and extrusion^24–26,28,29^. Potential efflux substrates from the cleft first enter the proximal binding pocket (PBP) of the access state located within PC1 and PC2 and activate a conformational change at the glycine-rich “switch loop” (**Figure 1b**)^30^. Substrates in the PBP or substrates entering from the vestibule move to the distal binding pocket (DBP) as the transporter cycles to the binding and then extrusion states for release of substrate to the open efflux tunnel^27^. However, specific residues within the PBP and DBP required for substrate recognition and conformational state change have yet to be elucidated.

**Figure 1.**
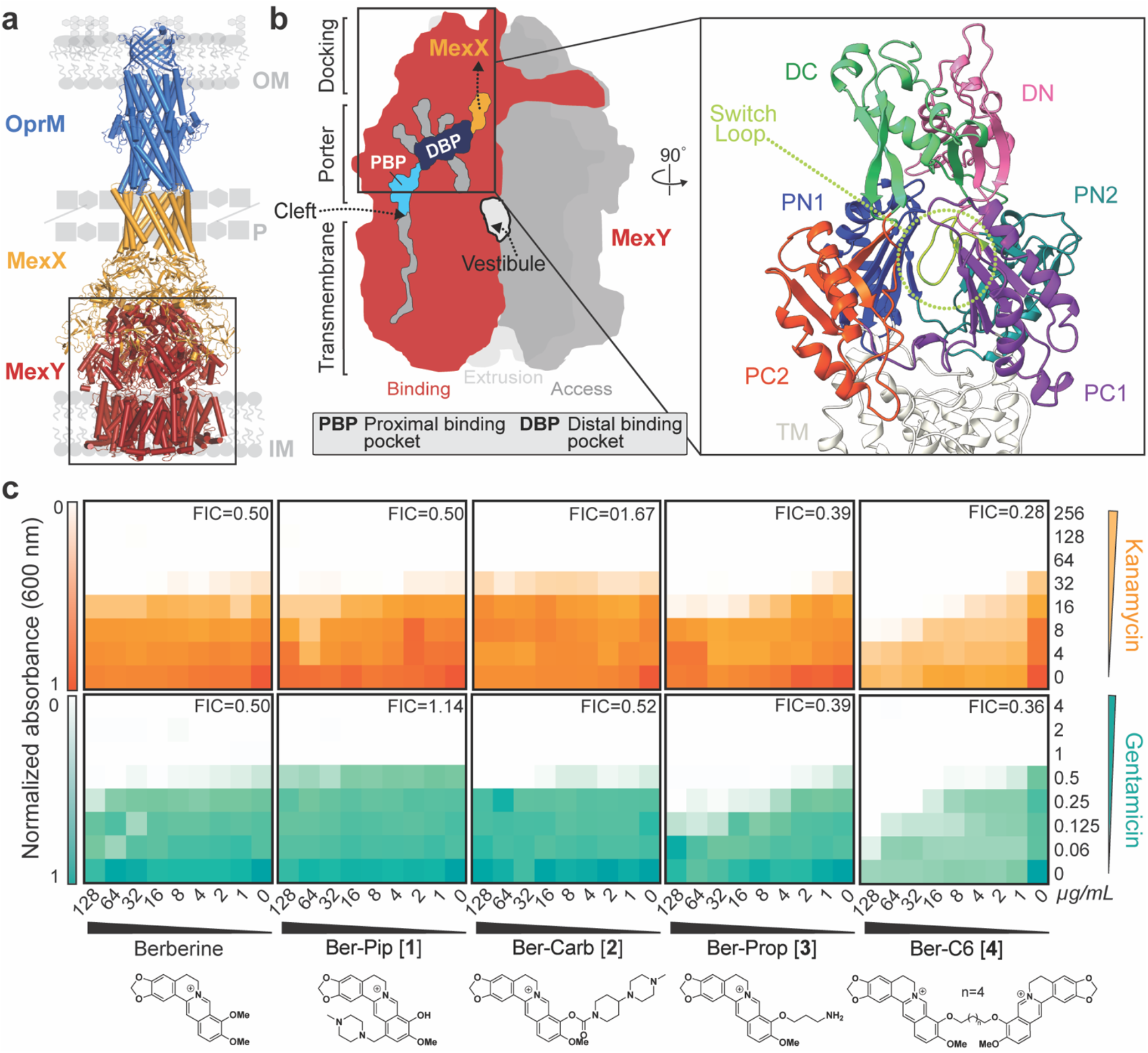
HTVS identifies berberine analogs as potential probes of MexY. (**a**) Homology model of the MexXY-OprM RND system comprising outer membrane (OM) protein OprM (blue), periplasmic (P) adaptor protein MexX (gold), and inner membrane (IM) transporter MexY (red). (**b**) Schematic of the MexY homotrimer in its three conformational states: access (dark grey), binding (red), and extrusion (light grey). The two putative main substrate entry channels (cleft and vestibule), primary substrate binding region (DBP and PBP), and path to MexX are indicated. A zoomed in view of the binding pocket structure (boxed) also highlights the MexX docking domain, DC (light green) and DN (pink), the porter domain, PN1 (blue), PN2 (teal), PC1 (purple), PC2 (orange), and the switch loop (lime green). (**c**) Checkerboard synergy assays in *P. aeruginosa* PAO1 with berberine and selected analogs from HTVS (*Generation 1*). Increased synergy is observed for Ber-Prop **[3]** and Ber-C6 **[4]** with Kan (top, orange) and Gen (bottom, teal) compared to berberine. Data are shown as the normalized mean of the optical density (OD_600_) of two biological replicates (0 is no growth, and 1 is maximum growth). The lowest Fractional Inhibitory Concentration (FIC) score for each compound-antibiotic pair is given in the upper right corner.

To provide insight into substrate recognition and/ or potential EPI binding, we aimed to identify potential molecular probes with distinct binding sites in the PBP /DBP region of MexY. Using the natural isoquinoline alkaloid berberine, a specific but weak MexY EPI^31,32^, as the starting scaffold in virtual screening against a homology model of MexY, we identified four berberine analogs as starting scaffolds. Through iterative rounds of structure-guided docking, molecular dynamics simulations, organic synthesis, and microbiological assays, we identified di-berberine conjugates that displayed increased synergy with aminoglycosides compared to natural berberine in *P. aeruginosa* PAO1, PA7, PA14, and pan-aminoglycoside resistant clinical isolates (**Supplementary Table 1**). Through alteration of the conjugate linker properties, we also demonstrate the applicability of these new berberine derivatives to identify binding pocket residues and substrate conformational properties which may play a role in specific substrate recognition by MexY.

## Results

### Virtual screening predicts four diverse berberine scaffolds as MexY chemical probes

*In silico* high-throughput virtual screening (HTVS) in the Schrödinger software was used to predict berberine analogs with enhanced binding scores and/ or distinct binding sites within the MexY PBP and DBP. Berberine was first used as a query on PubChem to generate a custom compound set comprising ∼10,000 potential ligands with a Tanimoto coefficient threshold >0.8. This ligand set was next docked using HTVS Glide (Schrödinger) into a binding pocket comprising the PBP and DBP of a *P. aeruginosa* PAO1 MexY (MexY^PAO1^) homology model using the protomer found in the binding conformational state (based on chain F of PDB 6IOL–see Methods for details)^33^. Ligands were ranked and the top scoring 1000 were subsequently re-docked with increased precision using Glide SP (Schrödinger). The top 1000 ligands were also clustered using a K-means algorithm based on chemical similarity to ensure selection of diverse scaffolds for further experimental investigation. This process produced four chemically diverse ligands (**Figure 2**, *Generation 1*): berberine conjugated with piperazine (Ber-Pip **[1]** and Ber-Carb **[2]**) or propanamine moieties (Ber-Prop **[3]**), and a di-berberine conjugate with a hexane linker (Ber-C6 **[4]**).

**Figure 2.**
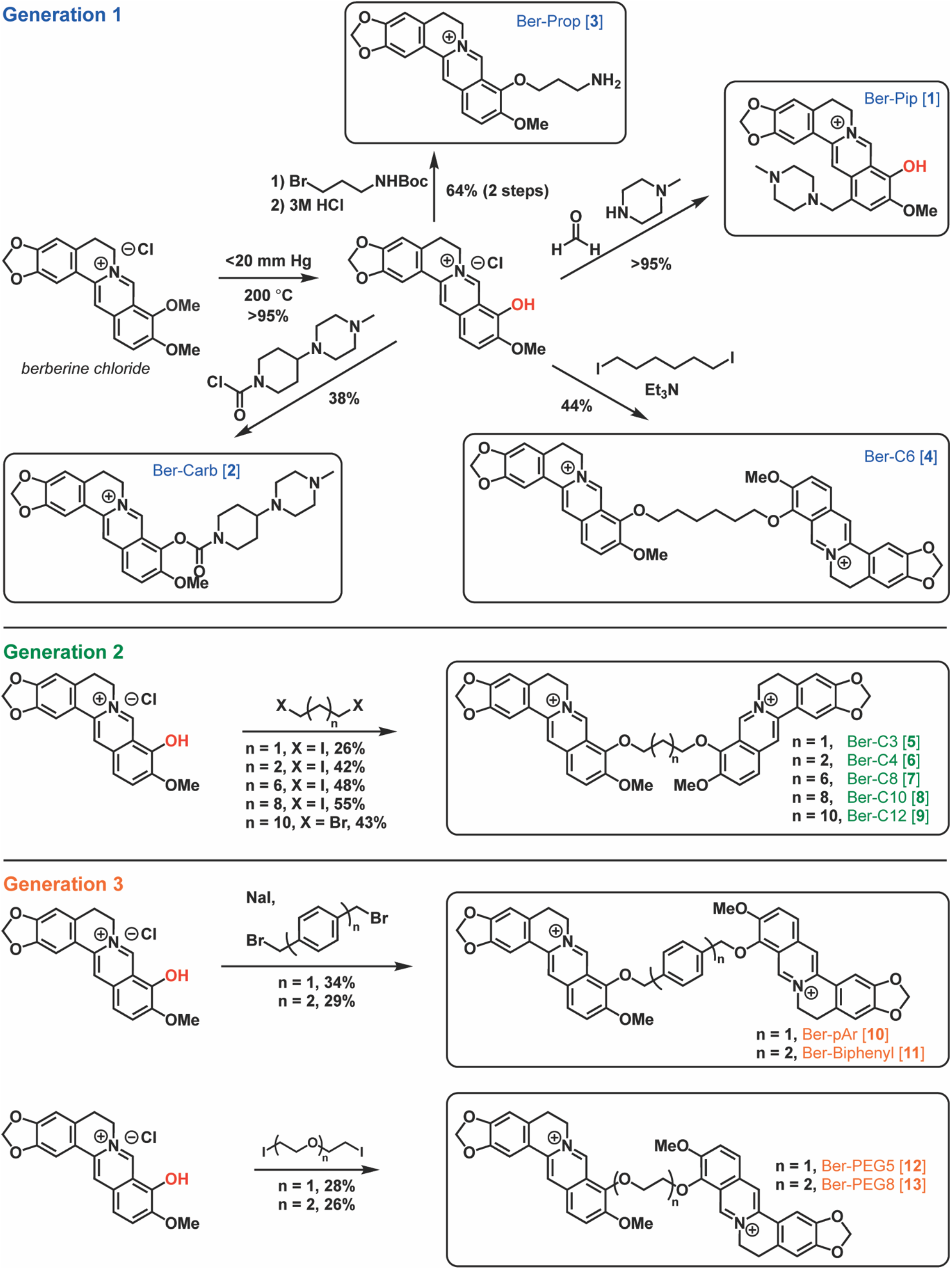
Overview of berberine analog synthesis. Overview of synthetic schemes to generate berberine and di-berberine conjugate analogs of Generation 1, Generation 2, and Generation 3. See Methods and Supplementary Materials for further details.

Our docking model predicted berberine binding to MexY^PAO1^ through multiple π-stacking interactions, including F610 in the switch loop and W177 in the DBP, as previously seen^31^ (**Figure 3a**). Furthermore, docking predicted three of the berberine analogs (Ber-Pip **[1]**, Ber-Prop **[3]**, Ber-C6 **[4]**) to have comparable or increased binding scores compared to berberine (**Supplementary Table 2**). In the DBP, Ber-Pip **[1]** and Ber-Carb **[2]** were predicted to have π-π stacking and cation-π interactions, respectively, while the terminal amino group of Ber-Prop **[3]** was positioned to interact with D124 (**Supplementary Figure 1a-c** and **Supplementary Table 3**). Ber-C6 **[4]** was predicted to bind in the DBP at the PC1/DN interface through multiple interactions, including π-stacking with Y752, cation-π interaction with K764, hydrogen bonding with the backbone of T176, and a salt bridge formed with E273 (**Supplementary Figure 1d** and **Supplementary Table 3**).

**Figure 3.**
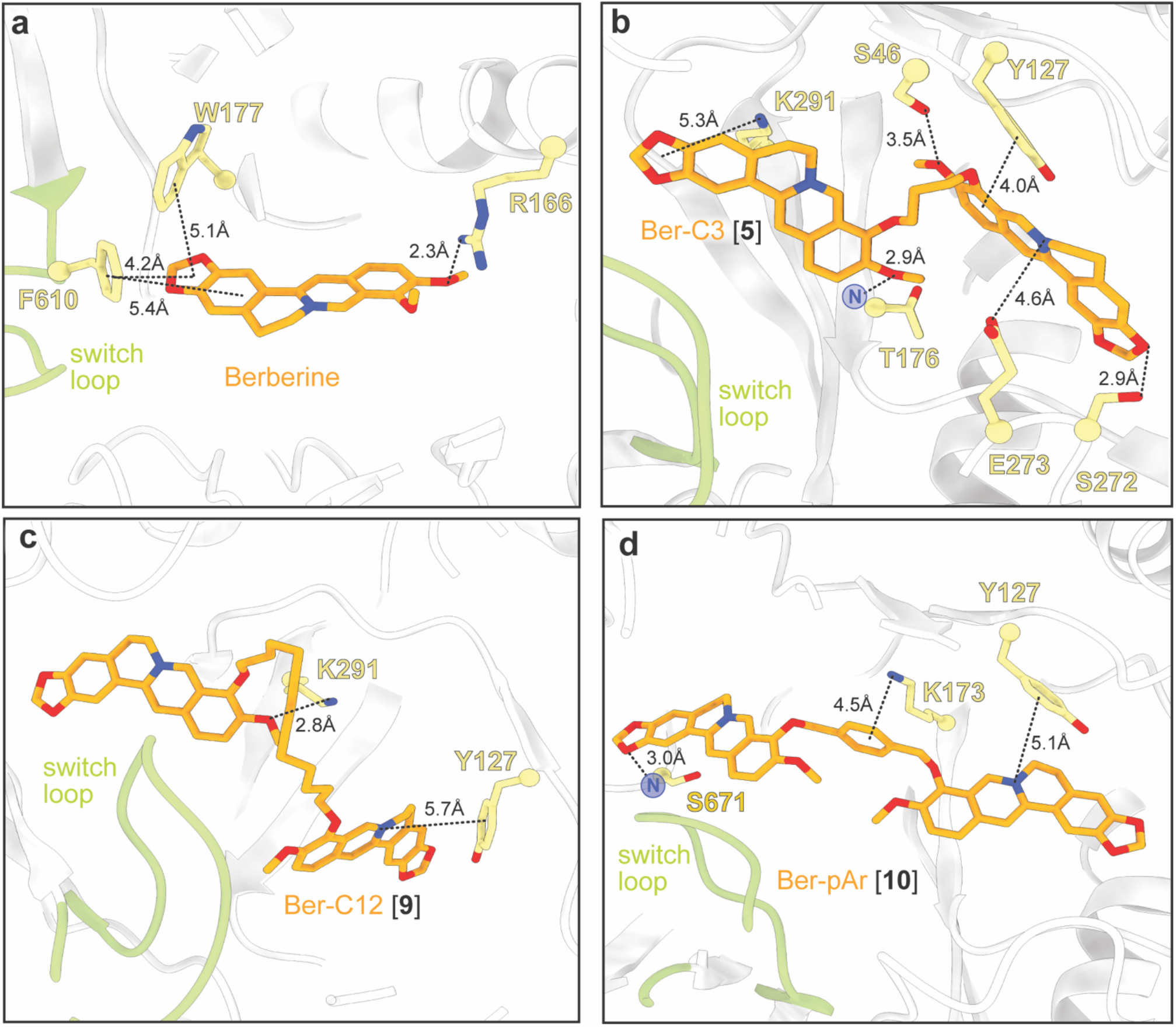
Predicted docking of select berberine analogs in the MexY^PAO1^ binding pocket reveals potential ligand-specific interactions. Docking of **(a)** berberine and the top three ligands of interest **(b)** Ber-C3 **[5]**, **(c)** Ber-C12 **[9]**, and **(d)** Ber-pAr **[10]** in the MexY^PAO1^ binding pocket spanning from the PBP to the DBP. Berberine docks in the DBP at the switch loop, Ber-C3 **[5]** and Ber-C12 **[9]** show preference for docking in the DBP, and Ber-pAr **[10]** spans the PBP and DBP.

Finally, to enhance the rigor of our ligand docking analyses and thus the inferences made from them, we generated a second independent MexY trimer homology model and again used the binding conformational state protomer (corresponding to chain L of PDB 6TA6) for docking of berberine, the four initially identified analogs, and all subsequently synthesized di-berberine conjugates. The periplasmic domains of both MexY models are very similar with an overall root mean square deviation of ∼0.8 Å, and, importantly, the trends observed for docking scores are consistent for the two models (**Supplementary Table 2**).

### Di-berberine conjugate has increased synergy with aminoglycosides

To test the validity of our computational modeling, a semisynthetic strategy was developed to obtain the top four ligands Ber-Pip **[1]**, Ber-Carb **[2]**, Ber-Prop **[3]** and Ber-C6 **[4]** (**Figure 2**–*Generation 1* and see Supplementary Methods). Synthesis began with selective C9-demethylation of commercial berberine chloride via vacuum pyrolysis to generate the natural product berberrubine chloride in high yield. The resulting phenol was appended to a commercially available tertiary amine scaffold via a carbamate linkage to give Ber-Carb **[2]**. Next, the electron-rich nature of the berberrubine ring system allowed for derivatization of C13 by double condensation of berberrubine chloride and N-methylpiperazine onto formaldehyde under acidic conditions, generating Ber-Pip **[1]**. Heating berberrubine chloride with 1,6-diiodohexane for 72 hours successfully generated the di-berberine conjugate Ber-C6 **[4]**, as well as a small amount of monohaloalkylated berberine. Because of the small amount of this isolated byproduct as well as its low reactivity, the propanamine derivative required an alternate approach in which berberrubine chloride was instead alkylated using 3-(Boc-amino)propyl bromide. Deprotection with hydrochloric acid then successfully afforded the desired ligand Ber-Prop **[3]** as the hydrochloride salt.

Checkerboard synergy assays were performed for berberine and the four synthesized analogs in *P. aeruginosa* PAO1 in the presence of two 4,6-deoxystreptamine (4,6-DOS) aminoglycosides, kanamycin (Kan) and gentamicin (Gen) (**Figure 1c**). Checkerboard assays confirmed berberine (4 µg/mL) is a weak inhibitor of the MexXY-OprM efflux pump, reducing the minimum inhibitory concentration (MIC) by two-fold and with a fractional inhibitory concentration (FIC) of 0.5, where FIC values < 0.5 are considered synergistic (**Figure 1c**, **Table 1**, and **Supplementary Table 4**). As MICs for berberine and analogs could not be experimentally determined, FIC value calculations used estimated MICs obtained via nonlinear regression of final culture OD_600_ (see Methods). Ber-Prop **[3]** and Ber-C6 **[4]** showed increased synergy with Kan and Gen, with Ber-C6 **[4]** reducing the MIC two-to eight-fold (FIC = 0.28-0.51). However, growth inhibition was observed in the presence of Ber-C6 **[4]** alone suggesting aminoglycoside-independent growth inhibition (**Figure 1c**, **Table 1**, and **Supplementary Table 4**). Piperazine ligands, Ber-Pip **[1]** and Ber-Carb **[2]** showed weaker or no synergy with the two aminoglycosides tested (**Figure 1c**,**Table 1**, and **Supplementary Table 4**). As a result, we selected the di-berberine conjugate, Ber-C6 **[4]**, as our focus for further development (**Figure 2**–*Generation 2*) due to its increased aminoglycoside synergism, higher predicted binding score, and more extensive and diverse predicted interactions within the DBP of MexY^PAO1^.

**Table 1.** Aminoglycoside minimum inhibitory concentration (MIC) against *P. aeruginosa* PAO1 in the presence of berberine and berberine analogs (64 µg/mL)

| Strain | Berberine analog | MIC; µg/mL<sup>1</sup> (fold change)<sup>2</sup> |  |  |  |
| --- | --- | --- | --- | --- | --- |
|  |  | Kan | Gen | Ami | Tob |
| PAO1 | - | 64 | 1 | 1 | 0.5 |
| PAO1Δ*mexXY* | - | 16 (4) | 0.25 (4) | 0.25 (4) | 0.25 (2) |
| PAO1 | Berberine | 32 (2) | 0.5 (2) | 0.5 (2) | 0.25 (2) |
|  | Ber-Pip [1] | 64 (-) | 1 (-) | *nd* | *nd* |
|  | Ber-Carb [2] | 32 (2) | 0.5 (2) | *nd* | *nd* |
|  | Ber-Prop [3] | 32 (2) | 0.5 (2) | *nd* | *nd* |
|  | Ber-C3 [5] | 16 (4) | 0.25 (4) | 0.25 (4) | 0.25 (2) |
|  | Ber-C4 [6] | 16 (4) | 0.25 (4) | 0.5 (2) | 0.25 (2) |
|  | Ber-C6 [4] | 16 (4) | 0.25 (4) | *nd* | *nd* |
|  | Ber-C8 [7] | 16 (4) | 0.125 (8) | *nd* | *nd* |
|  | Ber-C10 [8] | 8 (8) | 0.0625 (16) | 0.0125 (8) | 0.0625 (8) |
|  | Ber-C12 [9] | 8 (8) | 0.125 (8) | 0.125 (8) | 0.125 (4) |
|  | Ber-pAr [10] | 32 (2) | 0.25 (4) | 0.5 (2) | 0.25 (2) |
|  | Ber-Biph [11] | 32 (2) | 0.5 (2) | *nd* | *nd* |
|  | Ber-PEG5 [12] | 32 (2) | 0.5 (2) | 0.5 (2) | 0.25 (2) |
|  | Ber-PEG8 [13] | 32 (2) | 0.5 (2) | 0.5 (2) | 0.25 (2) |

<sup>1</sup>Determined from checkerboard and time-kill assays. Aminoglycosides: kanamycin (Kan), amikacin (Ami), gentamicin (Gen), and tobramycin (Tob); *nd*, not determined.
<sup>2</sup>Fold change relative to wild-type PAO1. (-) no fold change.

### Berberine alkane linker length defines MexXY-OprM inhibition efficacy and specificity

Di-berberine conjugates were designed with systematic variation in alkyl chain length connecting the two berberine groups to assess the role of linker length, flexibility, and overall size of the dimeric scaffold in predicted binding and MexXY-OprM inhibition. These *Generation 2* ligands with propane (Ber-C3 **[5]**), butane (Ber-C4 **[6]**), octane (Ber-C8 **[7]**), decane (Ber-C10 **[8]**), or dodecane (Ber-C12 **[9]**) linkers (**Figure 2**) were docked in MexY^PAO1^ binding pocket as before (**Figure 3b-c**, **Supplementary Figure 1e-g** and **Supplementary Table 2**).

An overall trend between longer linker length and increased predicted binding score was observed with the exception of the shortest alkane-linked conjugate, Ber-C3 **[5]** (**Supplementary Table 2**).The predicted lowest energy docked pose for Ber-C4 **[6]** differs from Ber-C6 **[4]**, with loss of hydrophobic interactions but gain of alternative stabilization via a salt bridge interaction with E175 in the DBP (**Supplementary Figure 1e** and **Supplementary Table 3**). In contrast, the di-berberine conjugate with the shortest linker, Ber-C3 **[5]**, is predicted to make a greater number of stabilizing interactions in the DBP through hydrogen bonding (to S46, T176 and S272), π-stacking/cation contacts (to Y127 and K291), and a salt bridge (with E273) (**Figure 3b** and **Supplementary Table 3**). Ber-C8 **[7]** was predicted to be stabilized through hydrogen bonding interactions with S46 and K79 in the PBP, while Ber-C10 **[8]** and Ber-C12 **[9]** had predicted interactions in the DBP with overlapping (K291), and distinct residues (E175 and Y127, respectively; **Figure 3d**, **Supplementary Figure 1f,g** and **Supplementary Table 3**). These docking results suggest that increased linker flexibility may allow the two berberine groups to independently adopt favorable conformations resulting in optimal interactions within the binding pocket, as no direct interactions were predicted with any alkane linker. Conversely, the Ber-C3 **[5]** analog can fortuitously adopt numerous stabilizing interactions within the DBP despite being more conformationally constrained.

Each of these new di-berberine conjugate analogs was obtained via reaction of berberrubine chloride with commercially available alkyl dihalides (**Figure 2**–*Generation 2* and see Supplementary Methods). Yields were moderate and increased consistently with alkyl chain length, suggesting that the dimerization reaction is slowed either by the steric repulsion of the bulky polycyclic rings or the electronic repulsion of the charged quaternary nitrogen atoms as they approach one another. This hypothesis is additionally supported by the observation that addition of a single berberrubine monomer to the alkyl halide was relatively rapid, leading to isolable monoalkylated product in less than one hour, but complete addition of a second berberrubine molecule took up to 72 hours.

Checkerboard synergy assays were performed using *P. aeruginosa* PAO1 with each di-berberine conjugate in combination with Kan and Gen (**Figure 4a** and **Supplementary Figure 2a**). Shortening the alkane linker in Ber-C3 **[5]** and Ber-C4 **[6]** reduced the MIC of Kan up to four-fold (FIC=0.30 and 0.38, respectively; **Figure 4a**, **Table 1**, **Supplementary Figure 2a** and **Supplementary Table 4**). Additionally, Ber-C4 reduced the MIC of Gen up to four-fold (FIC=0.38) (**Supplementary Figure 2a** and **Supplementary Table 4**). Ber-C3 at 64 µg/mL was able to reduce the MIC of Gen four-fold (**Table 1**), but at the highest concentration (128 µg/mL), reduced the Gen MIC eight-fold (FIC=0.30, **Figure 4a**). Increasing alkane linker length resulted in increased synergistic killing of PAO1 in combination with aminoglycosides, but with increased aminoglycoside-independent growth inhibition. Ber-C8 **[7]** reduced the MIC of Kan and Gen four-fold (FIC=0.28) and eight-fold (FIC=0.38), respectively, while Ber-C10 **[8]** and Ber-C12 **[9]** reduced the MICs of both aminoglycosides by up to 16-fold (**Figure 4a**, **Table 1**, **Supplementary Figure 2a** and **Supplementary Table 4**).

**Figure 4.**
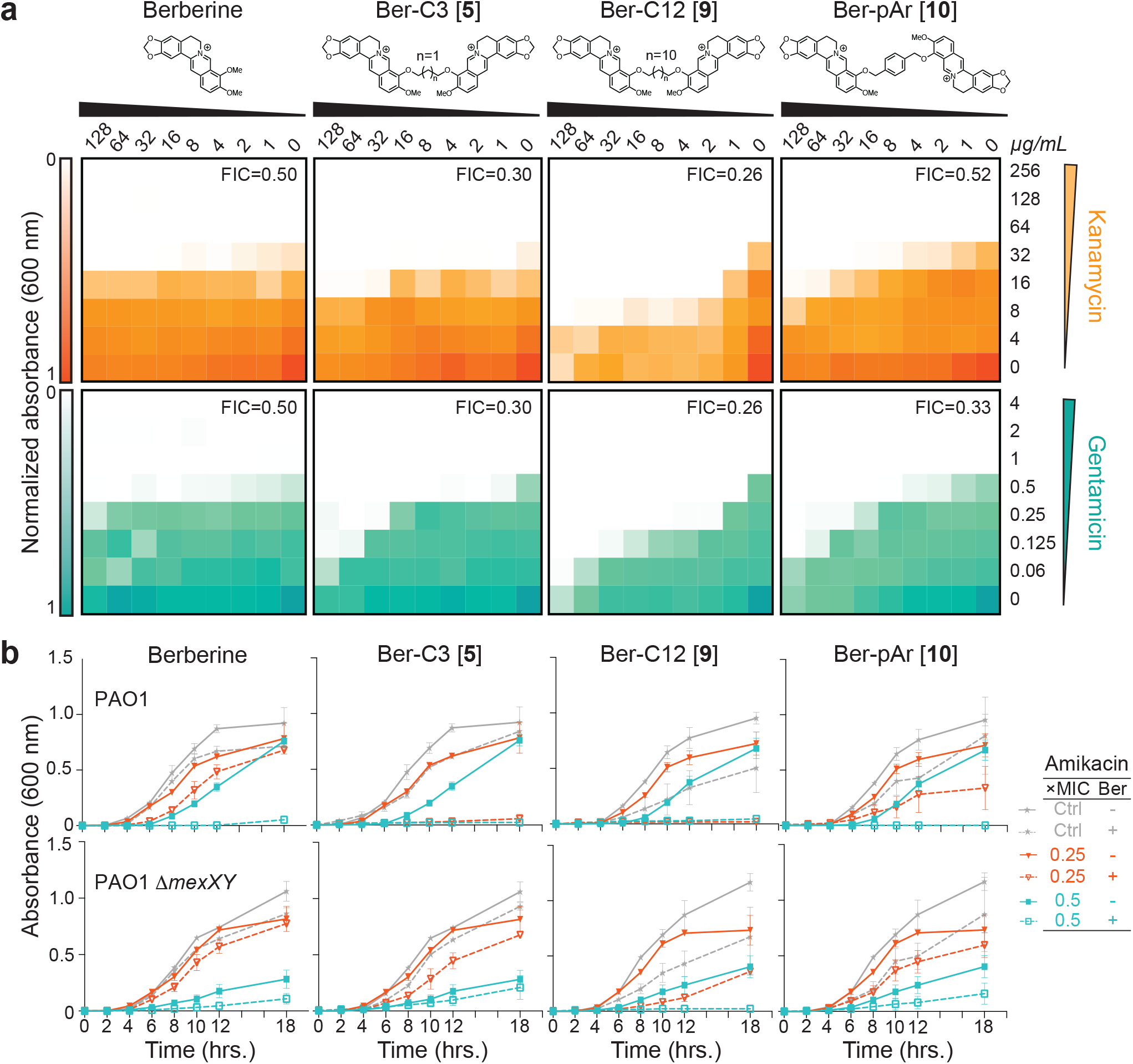
Top di-berberine conjugates exhibit increased synergy with aminoglycosides compared to berberine. **(a)** Checkerboard synergy assays with berberine and top selected di-berberine conjugates in *P. aeruginosa* PAO1. Increased synergy is observed with all ligands and kanamycin (top, orange) and gentamicin (bottom, teal) compared to natural berberine. The lowest fractional inhibitory concentration (FIC) score obtained is shown in the top right corner of each plot. Data are shown as the normalized mean of the optical density (OD_600_) of two biological replicates (0 is no growth, and 1 is maximum growth). **(b)** Time-kill assays for berberine and di-conjugate analogs at 64 µg/mL performed over 18 hours in both *P. aeruginosa* PAO1 (top) and its *mexXY* isogenic knockout (bottom). Synergy was observed with amikacin for all ligands tested, and Ber-C3 **[5]** and Ber-C12 **[9]** reduced the MIC four-fold (red open triangle) compared to no di-berberine conjugate (red filled triangle). Berberine and Ber-pAr **[10]** reduce the MIC by only two-fold (teal open square) at the tested concentration compared to no di-berberine conjugate (teal filled square). Significant aminoglycoside-independent growth inhibition is observed for Ber-C12 **[9]** (dashed grey line) compared to the no antibiotic growth control (grey line). Two-fold reduction of the MIC (black open square) for Ber-C12 **[9]** in *P. aeruginosa* PAO1Δ*mexXY* also suggests off-target activity. Data are shown as the average OD_600_ of biological replicates, each with two technical replicates, with error bars representing the standard deviation.

Next, to assess each di-berberine conjugate’s effect on growth phase and the potential for aminoglycoside specific interactions, we performed time-kill assays using *P. aeruginosa* PAO1 with two additional, structurally distinct 4,6-DOS aminoglycoside substrates, amikacin (Ami) and tobramycin (Tob; **Figure 4b**, **Supplementary Figure 3** and **Supplementary** Figure 4a,c). These assays were done in the presence of the di-berberine conjugates with the two shortest and two longest alkane linkers at 64 µg/mL as this concentration was predicted by checkerboard assays to allow differences to be observed across the range of linker lengths. As for Kan and Gen, berberine reduced the MIC two-fold for both Ami and Tob, indicating that its weak effect is independent of aminoglycoside substrate (**Table 1**, **Figure 4b**, and **Supplementary Figure 4a,c**). Consistent with its modestly tighter predicted binding, Ber-C3 **[5]** reduced the MIC for Ami four-fold (to 0.25 µg/mL) and Tob two-fold (to 0.25 µg/mL), while Ber-C4 **[6]** MIC reduction was equivalent to berberine (**Table 1**, **Figure 4b**,and **Supplementary Figure 4a,c**). For the longer alkyl-linked conjugate, Ber-C10 **[8]** reduced the MIC eight-fold for both Ami and Tob, whereas Ber-C12 **[9]** reduced the MIC of Ami eight-fold but the Tob MIC only four-fold (**Table 1**, **Figure 4b**, and **Supplementary Figure 4a,c**). Confirming the aminoglycoside-independent growth inhibition observed in previous checkerboard assays, growth curves of *P. aeruginosa* PAO1 in the presence of Ber-C10 **[8]** and Ber-C12 **[9]** showed a considerable extension of lag-phase growth switching to log-phase at 12 hours (compared to four hours) and a slower log-phase growth rate (**Figure 4b**, and **Supplementary Figure 4a,**c).

The observation of altered growth patterns in the time-kill assays with PAO1 next prompted further investigation of Ber-C3 **[5]**, Ber-C4 **[6]**, Ber-C10 **[8]**, and Ber-C12 **[9]** specificity for the MexXY-OprM efflux pump using equivalent experiments in a *P. aeruginosa* Δ*mexXY* strain (**Figure 4b**, **Supplementary Table 1**, and **Supplementary Figure 4b,d**). Loss of functional MexXY-OprM in the absence of berberine ligands, resulted in a four-fold MIC reduction for Ami and two-fold reduction for Tob (**Table 1**). The MICs for Ami and Tob in the presence of Ber-C3 **[5]** or Ber-C4 **[6]** do not exceed those observed in the Δ*mexXY* background suggesting these analogs with shorter alkane linkers are specific probes for the MexXY efflux system (**Table 1**). This conclusion was further validated in time-kill assays where addition of either compound in the Δ*mexXY* background did not alter growth patterns (**Figure 4b** and **Supplementary Figure 4b,d**). Of note, the Ber-C3 **[5]** compound reduced the MIC of both aminoglycosides to *mexXY* knockout levels, suggesting it may serve as a useful starting scaffold for MexXY-OprM EPIs. Consistent with previous checkerboard assays, Ber-C10 **[8]** and Ber-C12 **[9]** further reduced the MIC of Ami (eight-fold) and Tob (eight- and four-fold, respectively) to below those observed in the absence of MexXY-OprM (**Figure 4b**, **Table 1**, and **Supplementary Figure 4b,d**). Additionally, inactivation of MexXY-OprM did not influence the observed lag-phase extension or slower log-phase growth rate, suggesting the longer alkane conjugates are sensitizing the cell to aminoglycosides in both MexXY-dependent and -independent manners.

We considered the possibility that the longer alkane-linked conjugates might be disrupting membrane integrity, similar to amphipathic quaternary ammonium ligands with long hydrophobic chains^34,35^. However, using a sheep erythrocyte hemolysis assay, we found no evidence that berberine, Ber-C4 **[6]**, or either long alkane conjugates, Ber-C10 **[8]** or Ber-C12 **[9]**, is hemolytic at 128 µg/mL, a concentration at which MexXY-independent activity is clearly observed for the latter two analogs (**Supplementary Figure 5a**). Further, using a vancomycin susceptibility assay as a probe for membrane permeability^36^ in *P. aeruginosa* PAO1 and PAO1Δ*mexXY* resulted in identical vancomycin MICs with and without Ber-C12 **[9]** at 64 µg/mL, again suggesting that the off-target effect(s) of this analog are not related to membrane perturbation (**Supplementary Figure 5b,c**). The true additional target of the longer alkane conjugates is currently unknown and an area on-going investigation.

### Berberine and Ber-C3 **[5]** show antagonistic effects consistent with predicted overlapping binding in the MexY^PAO1^ DBP

Our structure-guided docking and checkerboard assays suggest that berberine and Ber-C3 **[5]** are specific inhibitors of the MexXY-OprM efflux system and bind at distinct but overlapping sites within the DBP (**Figure 3a-b**). If correct, berberine and Ber-C3 **[5]** should compete for binding in the DBP and would therefore not show synergistic activity when co-administered. Therefore, to provide an initial experimental validation of the ligand docking, this idea was tested in a three-way synergy assay using a fixed concentration of berberine (64 µg/mL) and a range of concentrations of Ber-C3 **[5]** (128-1 µg/mL) and Ami (4-0.125 µg/mL). Consistent with competition for binding, a weak antagonistic effect (two-fold increase in Ami MIC) or the same MIC as berberine alone was observed at most concentrations of Ber-C3 **[5]** (1-16 and 32-64 µg/mL, respectively; **Supplementary Table 5**). Only at 128 µg/mL was Ber-C3 **[5]** able to fully outcompete berberine for the overlapping binding pocket and resulting in an Ami MIC equivalent to analog the alone (0.25 µg/mL; **Supplementary Table 5**).

### Linker structure plays a role in MexXY-OprM inhibition efficacy and specificity

To further define how linker length, flexibility and chemical structure impact di-berberine conjugate activity and their potential as probes of MexXY function, four additional di-berberine conjugates were designed with alternative linkers: para-berberine substituents linked by one or two aryl groups, Ber-pAr **[10]** and Ber-Biph **[11]**, respectively, or by polyethylene glycol (PEG) linkers of different lengths, Ber-PEG5 **[12]** and Ber-PEG8 **[13]** (**Figure 2**–*Generation 3*). For example, if conjugate linker length is the primary determinant of efflux inhibition then di-berberine conjugates of a similar size with different linker types should maintain similar activity (Ber-PEG5 **[10]** *vs*. Ber-C4 **[6]**, Ber-pAr **[12]** *vs*. Ber-C6 **[4]**, Ber-PEG8 **[11]** *vs*. Ber-C8 **[7]**, and Ber-Biph **[13]** *vs*. Ber-C10 **[8]**. However, should flexibility or physico-chemical properties be critical, then conformationally more rigid aryl rings and/or more hydrophilic PEG moieties should show activity differences between the alkane, aryl, and PEG linkers of comparable length.

Computational docking of Ber-pAr **[10]**, Ber-Biph **[11]**, Ber-PEG5 **[12]**, and Ber-PEG8 **[13]** showed increased predicted binding scores for MexY^PAO1^ compared to their corresponding length alkane-linked conjugatess (**Supplementary Table 2**). Inspection of the lowest energy docking poses for the similarly sized analogs revealed Ber-pAr **[10]**, Ber-PEG5 **[12]**, and Ber-PEG8 **[13]** have unique predicted interactions in the binding pocket compared to their respective length-compared alkane-linked conjugates. Ber-pAr **[10]** and Ber-PEG5 **[12]** were both predicted to be stabilized by interactions spanning the PBP (via hydrogen bonding to S671) and DBP (via cation-π interaction with Y127) (compare **Figure 3d** and **Supplementary Figure 1d**, and **Supplementary Figure 1** panels **e** and **i**, respectively; and see **Supplementary Table 3**). Additionally, Ber-pAr **[10]** formed an additional linker-mediated cation-π interaction with K173 (**Figure 3d** and **Supplementary Table 3**). The longer PEG linker conjugate, Ber-PEG8 **[13]**, was predicted to be stabilized only in the DBP through a cation-π interaction (with Y127) and salt bridge (with E175) (compare **Supplementary** Figure 1 panels **f** and **j**; and see **Supplementary Table 3**). In contrast, Ber-Biph **[11]** shared predicted interactions (via salt bridge with E175 and hydrogen bonding to K291) with its corresponding length alkane-linked conjugate, Ber-C10 **[8]**, while gaining a linker-mediated cation-π interaction with K173 and an additional salt bridge with E273 (compare **Supplementary Figure 1** panels **g** and **h**; and see **Supplementary Table 3**). To complement these computational predictions with experimental assays of di-berberine conjugate activity, the four aryl- or PEG-linked conjugates were generated by chemical synthesis as before (**Figure 2**– *Generation* 3 and see Supplementary Methods). Synthesis of these *Generation 3* PEG-ylated analogs and aryl analogs proceeded similarly to the alkyl linkers over 72 hours despite the more activated benzylic electrophiles in the latter.

Checkerboard synergy assays were performed as before using *P. aeruginosa* PAO1 in combination with Kan and Gen (**Figure 4a** and **Supplementary Figure 2b**). Notably, increased predicted binding scores did not correlate with increased aminoglycoside synergy as previously observed for *Generation 1* and *Generation 2* analogs (**Table 1** and **Supplementary Table 2**). The Ber-pAr **[10]** analog showed synergy with only Gen, reducing the MIC up to four-fold (FIC = 0.33-0.52), and predicted FICs for both tested aminoglycosides were greater than the comparable Ber-C6 **[4]** analog (**Figure 4a** and **Supplementary Table 4**). Ber-Biph **[11]**, Ber-PEG5 **[12]**, and Ber-PEG8 **[13]** showed similar or decreased synergy with Kan and Gen compared to berberine, resulting in FIC values substantially higher than each corresponding length alkane-linked conjugate (**Supplementary Figure 2a,b** and **Supplementary Table 4**).

Time-kill assays in *P. aeruginosa* PAO1 showed only two-fold reduction in MIC for both Ami and Tob for Ber-pAr **[10]**, Ber-PEG5 **[12]**, and Ber-PEG8 **[13]** which is lower than the corresponding alkyl-linked conjugate in each case and comparable to berberine (**Figure 4b**, **Table 1**, and **Supplementary Figure 4a,c**). Ber-pAr **[10]** alone resulted a reduction in log-phase growth rate but neither Ber-PEG5 **[12]** nor Ber-PEG8 **[13]** affected overall growth rate (**Figure 4b** and **Supplementary** Figure 4a,c, respectively). To assess the specificity for the MexXY-OprM efflux pump, time-kill assays were performed in the isogenic Δ*mexXY* strain. While no difference in growth rate was again observed in the presence of the PEG-ylated conjugates, the reduced log-phase growth rate in the presence of Ber-pAr **[10]** observed in PAO1 was also observed in the Δ*mexXY* background (**Supplementary Figure 4b,d** and **Figure 4b**, respectively), suggesting this compound may exhibit off-target effects. However, as for the longer alkane-linked conjugates, we found no evidence for an effect on membrane integrity (**Supplementary Figure 5a**).

Overall, computational docking predicted increased binding affinities for *Generation 3* compounds compared to their similarly sized alkane-linked analogs, but this did not result in the anticipated increased synergy with aminoglycosides. We therefore propose the alkane linker flexibility may play an important role in ligand uptake into MexY and subsequent entry into the PBP/ DBP.

### Molecular dynamics (MD) simulations reveal important berberine probe characteristics

From the studies thus far, three ligands were selected as lead probes – Ber-C3 **[5]**, Ber-pAr **[10]**, and Ber-C12 **[9]** – reflecting the highest activity identified but with distinct linker flexibility, predicted binding pocket, and/ or specificity for MexXY-OprM. First, to more precisely define the nature and stability of predicted interactions made by berberine and these analogs with the MexY^PAO1^ DBP/ PBP, we performed five replica 50 ns MD simulations (Rep1 to Rep 5) in the Schrödinger Desmond module followed by MM/GBSA calculations on selected structures along each trajectory. The detailed analyses and following descriptions refer specifically to Rep1 (**Figure 5** and **Supplementary Figure 7a-c)**, but consistent results were obtained with all replica MD simulations and MM/GBSA calculations (**Supplementary Figure 6 and 7d**). Interestingly, berberine and Ber-C3 **[5]** showed greater overall conformational flexibility over the simulation, compared to Ber-pAr **[10]** and Ber-C12 **[9]** (**Figure 5a**). However, while similar total numbers of interactions were identified for each ligand pre- and post-simulation, only Ber-C3 **[5]** and Ber-pAr **[10]** maintained specific interactions over their trajectories (**Figure 5a** and **Supplementary Table 6**).

**Figure 5.**
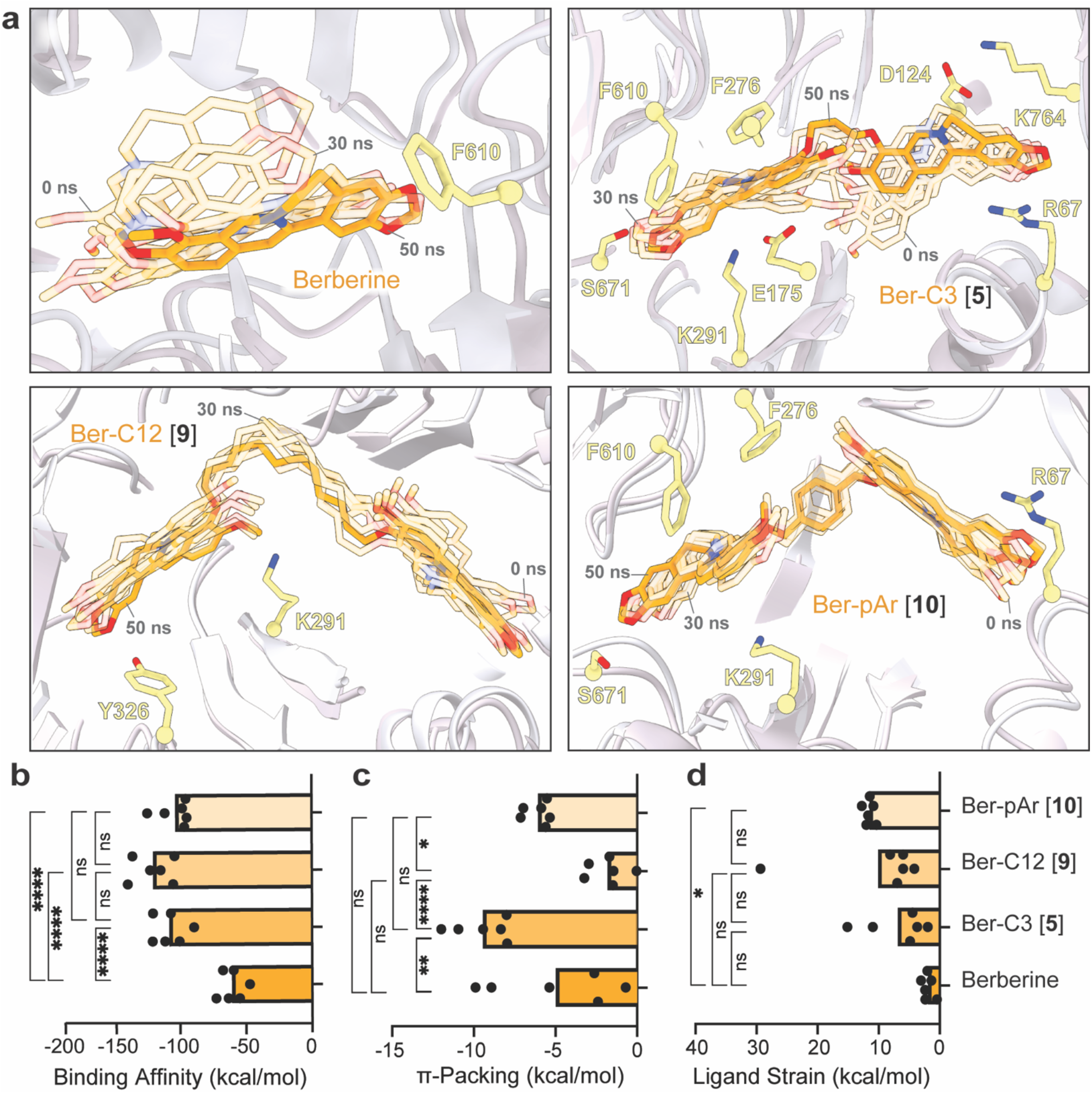
Molecular dynamics simulations of berberine and di-berberine conjugates reveal important characteristics for effective chemical probes of efflux function by MexY. **(a)** 50 ns molecular dynamics simulation (Rep1) of berberine (top left) and di-berberine conjugates, Ber-C3 **[5]** (top right), and Ber-C12 **[9]** (bottom left), and Ber-pAr **[10]** (bottom right) in the MexY^PAO1^ DBP and PBP. Conformations at the start (0 ns) and every 10 ns are shown as semi-transparent sticks (0 and 30 ns frames are indicated) and the final conformation (at 50 ns) is shown as solid sticks (orange). Parameters derived from MM/GBSA calculations: **(b)** binding affinity, **(c)** π-packing, and **(d)** ligand strain. Data shown are the individual calculated values for six frames (0 ns and one per 10 ns segment between 0-50 ns) with the average shown as a bar plot. Ber-C3 **[5]** was found to be an effective and specific efflux pump inhibitor for the MexXY-OprM efflux system compared to berberine, Ber-pAr **[9]** and Ber-C12 **[10]**. Statistical analysis was performed using one-way ANOVA on the mean for each ligand and a post-hoc Tukey test was used for pairwise comparison. Statistically significant results are shown (* pσ0.05, ** pσ0.01, *** pσ0.001, and **** pσ0.0001).

MM/GBSA calculations were performed on representative structures in each 10 ns window (100 frames) over the 50 ns simulations to obtain the relative free energy of each ligand in its binding pocket and thus reveal potentially important characteristics of a MexY inhibitor (**Figure 5b-d** and **Supplementary Figure 7a-c**). As expected, the overall average binding scores for Ber-C3 **[5]** (-116.0 kcal/mol), Ber-pAr **[9]** (-110.8 kcal/mol), and Ber-C12 **[10]** (-128.9 kcal/mol) were significantly greater than berberine (-65.23 kcal/mol; **Figure 5b**) where the total binding score is calculated from individual electrostatic, hydrogen bond, π-packing (π-mediated interactions), lipophilic, Van der Waals and solvation energetic scores.

The dimeric berberine ligands had significantly higher electrostatic interaction contributions than berberine, but there were no significant differences in hydrogen bonding contributions (**Supplementary Figure 7a-b**). Ber-C12 **[9]** was predicted to make significantly more lipophilic interactions in the binding pocket than Ber-C3 **[5]**, Ber-pAr **[10]**, and berberine, which contributed to its high predicted binding affinity (**Supplementary Figure 7c**). Reflecting its increased interactions with hydrophobic residues within the DBP during the MD simulation, Ber-C3 **[5]** has a significantly more favorable π-stacking mediated interactions, potentially due to enhanced flexibility seen within the simulation allowing it to position optimally to interact with the binding pocket (**Figure 5a,c**). This was further validated in the ligand strain energy calculation (the difference between the ligand’s free energy when bound compared to free in solution), where Ber-C3 **[5]** showed the lowest ligand strain energy (**Figure 5b,d**). MD simulations of berberine and lead analogs probes thus highlight characteristics, such as increased binding affinity, ability for multiple hydrophobic interactions, while maintaining flexibility and overall low ligand strain, as important for the development of MexXY-OprM specific efflux pump inhibitors.

### Top berberine alkane di-berberine conjugates are active against P. aeruginosa PA7 and PA14

To further investigate the efficacy and utility of our top lead probes, Ber-C3 **[5]**, Ber-pAr **[10]**, and Ber-C12 **[9]**, we evaluated synergy with Gen in two additional *P. aeruginosa* strains, PA7 and PA14 (**Supplementary Table 1** and **Table 2**). Alignment of the MexY protein sequence from *P. aeruginosa* PA7 (GenBank ABR84278; MexY^PA7^) and PA14 (GenBank ABJ11205; MexY^PA14^) with MexY^PAO1^ (GenBank OP868818) revealed 96.56% and 99.04% amino acid identity, respectively. This high conservation suggests that berberine and all analogs should maintain comparable predicted binding to the three MexY proteins. This was confirmed using SP Glide which produced binding scores that were broadly comparable to MexY^PAO1^ for both MexY^PA7^ and MexY^PA14^, with only a few larger differences (> 1 kcal/mol) and generally moderately tighter and weaker binding for MexY^PA7^ and MexY^PA14^, respectively (**Supplementary Table 2**). However, mapping the unique residues between the three *P. aeruginosa* strains on our MexY^PAO1^ homology model reveals that while most differences reside in the docking and transmembrane domains, four are located within the PBP/DBP: A589G (MexY^PAO1^), D149G (MexY^PA7^), S144A (MexY^PA7^) and E175Q (MexY^PA14^) (**Figure 6a**). Whether these differences result in differential activity of the berberine analogs was therefore explored with berberine and the three top selected analogs, Ber-C3 **[5]**, Ber-pAr **[10]**, and Ber-C12 **[9]**.

**Table 2.**
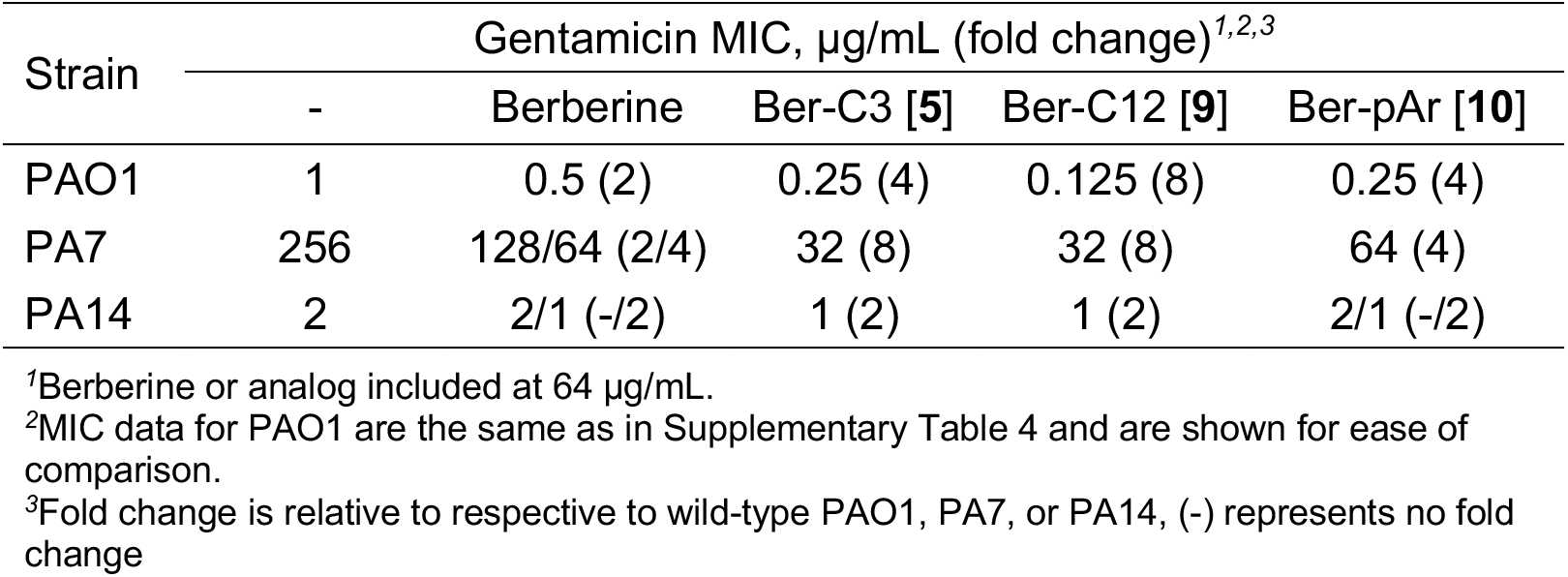
Gentamicin MIC against *P. aeruginosa* strains PA7 and PA14 in the absence and presence of berberine or select analogs.

**Figure 6.**
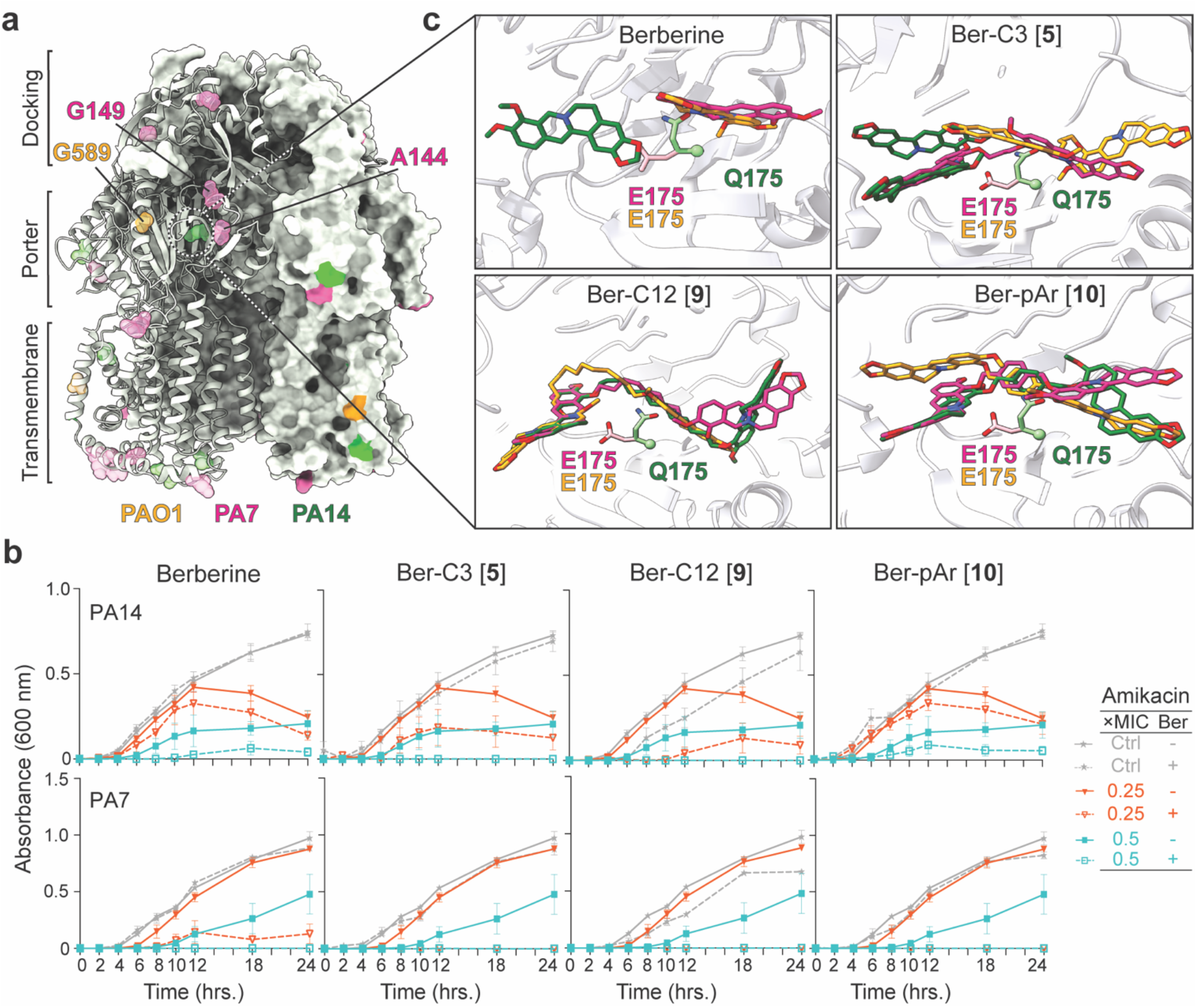
Berberine and di-berberine conjugates are active against *P. aeruginosa* PA7 and PA14. **(a)** Unique residues identified from alignment of MexY sequences from *P. aeruginosa* PAO1 (orange), PA7 (pink), and PA14 (green) mapped on a cartoon schematic of MexY^PAO1^ in the binding state. The three domains of the MexY transporter (docking, porter, and transmembrane) are indicated. **(b)** Time-kill assays for berberine and di-berberine conjugate analogs at 64 µg/mL performed over 18 hours in both *P. aeruginosa* PA14 (top) and PA7 (bottom). Synergy was observed in *P. aeruginosa* PA14 with Gen for Ber-C3 **[5]** and Ber-C12 **[9]** which reduced the MIC two-fold (teal open square) compared to no di-berberine conjugate present (teal filled square). In *P. aeruginosa* PA7, all dimer ligands tested reduced the MIC, at minimum, four-fold (red open triangle) at the tested concentration compared to no di-berberine conjugate (red filled triangle). No significant aminoglycoside-independent growth inhibition was observed for berberine or any di-berberine conjugate (dashed grey line) compared to the no antibiotic growth control (grey line). Data shown are the average OD_600_ measurement with error bars representing the standard deviation. **(c)** A zoomed in view of E175 conformation in MexY^PAO1^ (orange) and MexY^PA7^ (pink) which permits binding of berberine (top left), Ber-C3 **[5]** (top right), Ber-pAr **[10]** (bottom left), and Ber-C12 **[9]** (bottom right) in the DBP. In contrast, steric hinderance due to the Q175 conformation in MexX^PA14^ results in altered predicted ligand to bind in this region.

Time-kill assays were performed for berberine, Ber-C3 **[5]**, Ber-C12 **[9]**, and Ber-pAr **[10]**, at 64 μg/mL in combination with Gen against *P. aeruginosa* PA7 and PA14 (**Figure 6b**). Berberine reduced the MIC for Gen in PA7 two to four-fold and up to two-fold in PA14 (**Table 2** and **Figure 6b**). Alkane-linked conjugates, Ber-C3 **[5]** and Ber-C12 **[9]**, exhibited similar activity in each strain, reducing the Gen MIC eight-fold in PA7 and two-fold in PA14 (**Table 2** and **Figure 6b**). These results contrast to PAO1 where Ber-C3 **[5]** and Ber-C12 **[9]** differentially reduced the MIC of Gen four-fold and eight-fold, respectively. For the aryl-linker analog, Ber-pAr **[10]** reduced the MIC of Gen four-fold in PA7, comparable to PAO1, but reduced the MIC up to 2-fold in PA14. (**Table 2** and **Figure 6b**). Overall, changes in Gen MICs in PAO1 were comparable to those observed in PA7, but all ligands were less effective in PA14 (**Table 2**).

We next used computational docking to investigate whether any of the binding pocket residue changes identified in the MexY sequence alignment (**Figure 6a**) might be responsible for the observed differences in synergism in PAO1/ PA7 compared to PA14. Docking of Ber-C3 ****[5]****, Ber-pAr ****[10]****, and Ber-C12 ****[9]**** in homology models of MexY^PAO1^, MexY^PA7^, and MexY^PA14^ elucidated important differences in predicted binding poses of the di-berberine conjugates, suggesting that inherent differences between each MexY binding pocket may indeed exist (**Supplementary Figures 8** and **9**). In particular, a predicted interaction with MexY from three strains was observed at residue 175, i.e. MexY^PAO1^/ MexY^PA7^ E175 and MexY^PA14^ Q175 (**Figure 6c** and **Supplementary Figure 8b-d**). In MexY^PAO1^/ MexY^PA7^, E175 extends into the binding pocket, whereas Q175 of MexY^PA14^ adopts a distinct conformation resulting in potential clash with the berberine analogs (**Figure 6c**). This observation suggests that the identity and thus positioning of the residue 175 side chain may influence the preferred ligand conformation and subsequent binding pocket interactions. For example, Ber-C3 ****[5]**** docks in an ‘extended’ conformation in MexY^PAO1^/ MexY^PA7^ but a ‘compact’ conformation in MexY^PA14^, resulting in a potential loss of important cation-π/ π-π stacking interactions (with Y127 and K291) and salt bridges (to E273 in MexY^PAO1^ and E175 in MexY^PA7^). These predicted differences in interactions resulting from the change at residue 175 have the potential to explain the decreased fold-change in Gen MIC observed for PA14, but require future testing *in vivo* (**Figure 6c**, **Table 2**, and **Supplementary Figures 8b** and **9**). Thus, these results thus exemplify the potential of di-berberine conjugates can be used as functional probes of the MexY binding pocket to elucidate important residues involved in ligand recognition.

### Di-berberine conjugates are active against pan-aminoglycoside resistant clinical isolates and show preferential synergy with aminoglycoside substrates of MexXY-OprM

Having defined activity in *P. aeruginosa* PAO1, PA7 and PA14, we extended these studies to further explore the potential utility of the top three di-berberine conjugate analogs by assessing efflux inhibition activity in two pan-aminoglycoside resistant *P. aeruginosa* clinical isolates, K2156 and K2161^19^ (**Supplementary Table 1**). Alignment of the MexY protein sequence from *P. aeruginosa* K2156 (GenBank OP868819; MexY^K2156^) and K2161 (GenBank OP868820; MexY^K2161^) with MexY^PAO1^ (GenBank OP868818) revealed 99.52% and 99.62% amino acid identity, respectively. Unique residues from both clinical isolates were primarily located in the transmembrane region of the MexY^PAO1^ homology model, with no observed changes in the PBP/ DBP binding pocket (**Supplementary Figure 10**). However, both MexY^K2156^ and MexY^K2161^ have Q840E (cleft) and T543A (transmembrane) amino acid substitutions, with additional unique V980I (transmembrane) changes in MexY^K2156^ and A33V (vestibule) and D428N (central cavity) in MexY^K2161^ (**Supplementary Figure 10**).

MIC assays were performed for clinical isolates K2156 and K2161 in the presence of 4,6-DOS aminoglycosides (Kan, Ami, Gen, and Tob) and 64 µg/mL of berberine or di-berberine conjugate (**Table 3**). To compare berberine ligand synergy with aminoglycosides, MICs for the isogenic Δ*mexXY* knockout for each clinical isolate were also determined (**Table 3** and **Supplementary Table 1**). Berberine, Ber-C3 ****[5]****, and Ber-pAr ****[10]**** exhibited increased synergy against K2156, with a eight-to 16-fold reduction in all MICs, where 16-fold reduction represented complete MexXY-OprM inhibition. Ber-C12 ****[9]**** reduced the MIC of Kan below Δ*mexXY* knockout levels (32-fold reduction; **Table 3**). Additionally, berberine and the tested di-berberine conjugates all showed increased synergy of four to eight-fold MIC reduction with Tob, which were below the two-fold reduction observed in the Δ*mexXY* background (**Table 3**). In contrast, a low to moderate change (zero to four-fold) reduction in Kan, Ami and Gen MICs was observed for K2161 in the presence of berberine and di-berberine conjugates, of which Ber-C3 ****[5]**** and Ber-C12 ****[9]**** showed increased synergy compared to berberine and Ber-pAr ****[10]****. Interestingly, Ber-C3 ****[5]**** and Ber-C12 ****[9]**** inhibited Tob to Δ*mexXY* knockout levels (four-fold reduction), but none of the berberine compounds inhibited below observed knockout levels, suggesting off-target effects may also be strain dependent (**Table 3**). The variable synergies exhibited with pan-aminoglycoside clinical isolates in the presence of berberine, Ber-C3 ****[5]****, Ber-C12 ****[9]****, and Ber-pAr ****[10]**** in combination with homology modeling suggests these chemical probes could potentially be used to further understand efflux substrate interactions at entry channels (e.g., cleft, vestibule, central cavity) into MexY.

**Table 3.** Antibiotic minimum inhibitory concentration (MIC) in absence or presence of select berberine analogs for *P. aeruginosa* PAO1 and two clinical pan-aminoglycoside resistant strains (K2156 and K2161)

| Genotype<br>Analog<sup>1</sup> | Antibiotic MIC;µg/mL<sup>3</sup> (fold change)<sup>4</sup> |  |  |  |  |  |
| --- | --- | --- | --- | --- | --- | --- |
|  | WT | ΔXY<sup>2</sup> | WT |  |  |  |
|  | - | - | Berberine | Ber-C3 [5] | Ber C12 [9] | Ber-pAr [10] |
| *P. aeruginosa* PAO1 |  |  |  |  |  |  |
| Kan | 64 | 16 (4) | 32 (2) | 16 (4) | 8 (8) | 32 (2) |
| Ami | 1 | 0.25 (4) | 0.5 (2) | 0.25 (4) | 0.125 (8) | 0.5 (2) |
| Gen | 1 | 0.25 (4) | 0.5 (2) | 0.25 (4) | 0.125 (8) | 0.25 (4) |
| Tob | 0.5 | 0.25 (2) | 0.25 (2) | 0.25 (2) | 0.125 (4) | 0.25 (2) |
| Tig | 4 | 0.5 (8) | 4 (-) | 2 (2) | 2 (2) | 4 (-) |
| Cef | 1 | 0.5 (2) | 1 (-) | 0.5 (2) | 2 (+2) | 1 (-) |
| Cip | 0.125 | 0.125 (-) | 0.125 (-) | 0.125 (-) | 0.5 (+4) | 0.25 (+2) |
| Imi | 4 | 4 (-) | 4 (-) | 4 (-) | 2 (2) | 2 (2) |
| Pan-aminoglycoside resistant *P. aeruginosa* clinical isolate K2156 |  |  |  |  |  |  |
| Kan | 256 | 16 (16) | 16 (16) | 16 (16) | 8 (32) | 16 (16) |
| Ami | 8 | 0.5 (16) | 0.5 (16) | 0.5 (16) | 0.5 (16) | 0.5 (16) |
| Gen | 8 | 0.5 (16) | 1 (8) | 0.5 (16) | 0.5 (16) | 1 (8) |
| Tob | 2 | 1 (2) | 0.5 (4) | 0.25 (8) | 0.5 (4) | 0.5 (4) |
| Tig | 4 | 0.5 (8) | 4 (-) | 2 (2) | 1 (4) | 4 (-) |
| Cef | 2 | 1 (2) | 2 (-) | 2 (-) | 2 (-) | 2 (-) |
| Cip | 1 | 1 (-) | 1 (-) | 1 (-) | 4 (+4) | 2 (+2) |
| Imi | 2 (-) | 2 (-) | 2 (-) | 2 (-) | 2 (-) | 2 (-) |
| Pan-aminoglycoside resistant *P. aeruginosa* clinical isolate K2161 |  |  |  |  |  |  |
| Kan | 256 | 64 (4) | 256 (-) | 128 (2) | 128 (2) | 256 (-) |
| Ami | 16 | 2 (8) | 8 (2) | 4 (4) | 8 (2) | 8 (2) |
| Gen | 16 | 2 (8) | 8 (2) | 4 (4) | 4 (4) | 8 (2) |
| Tob | 8 | 2 (4) | 4 (2) | 2 (4) | 2 (4) | 4 (2) |
| Tig | 4 | 1 (4) | 4 (-) | 4 (-) | 2 (2) | 4 (-) |
| Cef | 0.25 | 0.125 (2) | 0.5 (+2) | 0.25 (-) | 1 (+4) | 1 (+4) |
| Cip | 0.125 | 0.06 (2) | 0.25 (+2) | 0.25 (+2) | 0.5 (+4) | 0.25 (+2) |
| Imi | 2 | 2 (-) | 2 (-) | 2 (-) | 1 (2) | 2 (-) |

<sup>1</sup>Berberine or dimeric analog included at a fixed concentration of 64 µg/mL where indicated.
<sup>2</sup>Isogenic MexXY knockout of the corresponding parental strain, PAO1 or clinical isolate.
<sup>3</sup>Antibiotics include MexXY-OprM substrates kanamycin (Kan), amikacin (Ami), gentamicin (Gen), tobramycin (Tob), tigecycline (Tig), cefepime (Cef), and ciprofloxacin (Cip), and the nonsubstrate imipenem (Imi).
<sup>4</sup>Fold decrease relative to respective wild type PAO1, K2156, or K2161. (-) indicates no change. (+) indicates an increase in MIC.

Finally, to assess the ability of these dimeric berberine probes to provide insight into binding preferences of other, non-aminoglycoside substrates and their routes through the MexY transporter, we determined MICs for a representative panel of additional antibiotics: tigecycline (Tig), cefepime (Cef), ciprofloxacin (Cip); and the non-MexY-substrate imipenem (Imi) (**Table 3**). Ber-C3 ****[5]**** and Ber-C12 ****[9]**** reduced the MIC of Tig two-to four-fold in PAO1 and K2156, whereas Ber-C12 ****[9]**** produced only a two-fold reduction in K2161 (**Table 3**). Additionally, Ber-C3 ****[5]**** reduced the MIC two-fold for the substrate Cef in PAO1. All other interactions between non-aminoglycoside substrates (Cef or Cip), *P. aeruginosa* strains (PAO1, K2156, or K2161), and ligands (berberine, Ber-C3 ****[5]****, Ber-C12 ****[9]****, or Ber-pAr ****[10]****) resulted in either no difference in observed MIC or an increase in resistance (**Table 3**). Such increase in resistance is most likely due to increased efflux of these substrates by another RND-efflux system in *P.* aeruginosa. Additionally, a two-fold decrease in the MIC of non-substrate Imi was observed in the presence of Ber-C12 ****[9]**** (PAO1 and K2161) and Ber-pAr (PAO1) (**Table 3**). These data suggest that the berberine alkyl conjugates can be used as aminoglycoside-specific probes for MexY efflux function across different *P. aeruginosa* strains. Additionally, our di-berberine conjugates may serve as useful tool for understanding crosstalk between multiple RND systems in *Pseudomonas*.

## Discussion

The development of efflux pump inhibitors (EPIs) remains challenging due to limitations in our understanding of substrate recognition and efflux mechanisms, as well as the resource-intensive nature of screening chemical libraries *in vitro* for potential hits^37^. Although no EPIs have progressed to clinical approval, those discovered such as phenylalanine arginyl β-napthylamide (PAβN) and carbonyl cyanide-m-chlorophenylhydrazone (CCCP) have provided valuable tools to study efflux function ^38–42^. However, these general inhibitors suffer significant limitations such as being substrates for multiple RND systems within the same organism or having off-target effects, both of which limit their use as tools to study specific RND efflux transporter function^42^. Therefore, we aimed to identify a pathway to new chemical probes, specific for the MexXY-OprM efflux system, to elucidate efflux function and inhibition.

Here, based on its known MexXY-OprM EPI activity, we used the naturally occurring and well-studied alkaloid berberine as a starting scaffold for a structure-guided computational screen^31,32,43–48^. HTVS identified four berberine analogs (*Generation 1*) as potential leads with diverse predicted binding preferences in the MexY PBP/ DPB. Among the *Generation 1* ligands, addition of piperazine moieties to monomeric berberine did not enhance inhibition as previously shown for other pyridylpiperazine-based allosteric inhibitors of RND efflux systems^49^. However, our studies identified an alkane-linked di-berberine conjugate, Ber-C6 ****[4]****, as an alternative promising lead for MexY. Di-berberine conjugates have been shown to bind preferentially to G-quadruplex DNA or ribosomal RNA but, to our knowledge, have yet to be characterized as protein inhibitors^50–54^.

Using di-berberine conjugates of variable linker length (*Generation 2*) and chemical nature (*Generation 3*), we identified linker characteristics that affect inhibition of the MexXY-OprM efflux system. Generally, alkyl linker length correlated with increased binding scores, and these predictions were corroborated by biological data showing improved synergy with aminoglycosides. However, the exception to this trend was observed for the analog with the shortest linker, Ber-C3 ****[5]****), which had higher predicted binding scores, due to the increased predicted hydrophobic interactions. Again, however, biological assays corroborated these computational predictions, revealing that this ligand had increased synergy with aminoglycosides and reduced the MIC to Δ*mexXY* knockout levels, a two-to four-fold decrease. This range is recognized by the Clinical and Laboratory Standards Institute as the aminoglycoside clinical breakpoint (from sensitive to resistant), highlighting the importance of MexXY efflux inhibition for favorable treatment outcomes. Importantly, Ber-C3 ****[5]**** is non-toxic to pseudomonal cells and appears specific for the MexXY-OprM system, unlike previously identified berberine analogs^47,48^ Of note, longer alkyl analogs did exhibit clear aminoglycoside-independent growth inhibition, but contrary to expectation our data suggest that this phenomenon is not attributable to membrane disruption. Berberine was identified to negatively impact DNA topoisomerase I/II and DNA, causing cell cycle arrest and DNA damage^55–59^. In the absence of membrane effects, we therefore speculate that the longer alkyl-linked di-berberine conjugates may act similarly and studies are currently ongoing to define the basis of these off-target effects.

Initial docking studies of berberine in the MexY^PAO1^ binding pocket identified interactions with the switch loop as has previously been observed^31^. Our docking of additional berberine ligands was also validated through a three-way synergy assay (berberine, Ber-C3 ****[5]****, and amikacin) in PAO1 which supported the predicted overlapping binding sites for these ligands. Docking analyses with berberine alkyl-linked analogs predicted multiple binding sites within the substrate binding region of MexY, including the PBP, switch loop, DBP, and at the DBP-porter interface, highlighting the potential utility of these di-berberine conjugates as chemical probes for understanding the ligand translocation pathway and its kinetics. Although binding scores for analogs with alternative aryl or PEG linkers were higher compared to the comparably sized alkyl-linked analogs, these predictions were not borne out in the biological analyses for these ligands. This discrepancy may have arisen due to known issues with MM/GBSA binding energy over-estimation between sets of ligands^60^, or because docking analysis does not consider other processes that might impact activity. For example, analogs with more rigid and/ or hydrophobic linkers may be less readily able to reach the MexY PBP/ DBP. As such, further detailed analyses will be essential to fully cross-corroborate these *in-silico* predictions. However, based on docking and MD simulations, we propose that possessing a short but slightly flexible linker that allows for optimal π-stacking interactions in the binding pocket is an important characteristic for di-berberine conjugate binding to MexY.

Antimicrobial susceptibility testing (e.g., checkerboard assays, MICs, and time-kill assays) also revealed preferential inhibition by di-berberine conjugates for different drugs of the same class. For example, Ber-pAr ****[10]**** showed increased synergy with gentamicin compared to other aminoglycosides tested. Further development toward these specificities could thus allow berberine analogs to be used to probe specific binding regions for substrates the MexXY-OprM efflux system. Additionally, reduction in MIC for non-aminoglycoside substrates was possible, but inhibiting to the level of MexXY knockout in *P. aeruginosa* strains was inconsistent, suggesting that these substrates can more efficiently outcompete berberine analogs for binding or have alternative binding site(s), or that they are also effluxed by other RND systems^61–64^. Further, we found that Ber-pAr ****[10]**** and Ber-C12 ****[9]**** had a strain-dependent effect on enhanced susceptibility to non-efflux substrate imipenem, a phenomenon also previously observed in *P. aeruginosa* PAO12 with berberine^65^. Importantly, our top di-berberine conjugate analogs, Ber-C3 ****[5]****, Ber-C12 ****[9]****, and Ber-pAr ****[10]****, showed promising activity against two pan-aminoglycoside resistant clinical *P. aeruginosa* isolates with variable MexXY-OprM dependencies, K2156 and K2161, as well as *P. aeruginosa* PA7 and PA14 strains. As proof-of-concept for the utility of the di-berberine conjugates as chemical probes, we identified a putative *P. aeruginosa* strain specific residue (MexY^PAO1^/ MexY^PA7^ E175 *vs*. MexY^PA14^ Q175) which may influence inherent properties of the MexY binding pocket, as observed in the corresponding residue mutation, Q176K, of the homolog *Salmonella* AcrB transporter^66^, and therefore subsequent efflux inhibition. Through checkerboard synergy assays, we identified that Ber-C3 ****[5]**** has reduced synergy with aminoglycosides in PA14 compared to PAO1 and PA7. Docking of Ber-C3 ****[5]**** into all three strains MexY homology models predicted ligand conformational changes due to the position of the 175 residue for glutamic acid versus glutamine. These data highlight the functionality of di-berberine conjugates as probes to identify putative important residues involved in MexXY-OprM efflux inhibition in multiple *P. aeruginosa* strains.

In summary, using chemical biology and small-molecule discovery, we generated a panel of potential probes of the transporter component of the MexXY-OprM efflux pump in *P. aeruginosa*. As RND efflux pumps are ubiquitous in Gram-negative pathogens and contribute extensively to multidrug resistance, understanding the chemical and mechanical properties of efflux pumps and their respective substrates is imperative. Further development of chemical probes, such as our lead di-berberine conjugates, can shed light on mechanisms that govern efflux pump substrate recognition in RND efflux pumps and help drive new strategies for specific EPI development.

## Methods

### Homology modeling of MexY and MexY-Per

The amino acid sequences of the MexY from *P. aeruginosa* PAO1, PA7 and PA14 were retrieved from The *Pseudomonas* Genome Database^67^ (accession codes GCF_000006765.1, GCF_000017205.1, and GCF_000404265.1, respectively). Homology models of the asymmetric MexY homotrimer were constructed using two independent cryo-EM structures of MexB as the template (PDB 6IOL chains E, F and G, and 6TA6 chains J, K and L) on the Swiss Model server (https://swissmodel.expasy.org). Although at modest resolution compared to some other available structures of MexB alone, these templates were selected with the expectation that the trimeric MexB structure within the full efflux pump would best represent the relevant conformational states of the transporter protein^68^. Models built with the Swiss Model server were minimized in MacroModel (Schrödinger LLC 2022-3) using the OPLS4 force field and Polak-Ribier Conjugate Gradient (PRCG) minimization method with 2500 iterations. A MexY^PAO1^ model was also generated with the transmembrane regions removed and the remaining periplasmic components joined with Gly/Ser linker (“MexY-per”, comprising residues: MexY^1–38^-GSGSGGSGGS (linker)-MexY^558–863^). The MexY-per model was constructed using the full *P. aeruginosa* PAO1 MexY homology model as the starting template in Biologics (Schrödinger LLC 2022-3) and with enhanced loop sampling for the linker region.

### High-throughput virtual screening (HTVS) and molecular docking

The PyMol Caver plugin^69^ was used to identify potential functional pockets in the three conformationally distinct protomers of each trimeric MexY^PAO1^ homology model. For HTVS and docking studies, we selected the binding conformation protomer defined by the largest combined volume of PBP and DBP identified using Caver (i.e., corresponding to 6IOL chain F and 6TA6 chain L). The MexY^PAO1^ model from *P. aeruginosa* PAO1 was prepared for docking in Protein Preparation Wizard and energy minimized prior to grid creation with OPLS4 force field (Schrodinger LLC 2022-3). For HTVS, a ligand set (∼10,000 berberine analogs) was created using berberine as the query in PubChem. To ensure chemical diversity within this ligand set, we applied a Tanimoto coefficient cutoff of 0.8 and resultant ligands were prepared in LigPrep with the OPLS4 force field (Schrödinger LLC 2022-3). A region comprising the combined PBP and DBP of MexY^PAO1^ was targeted using a docking grid centered between MexY residues F610 and F623 with a 30 Å^3^ outer grid (where the ligands can dock) and 15 Å^3^ inner grid (where the ligand centroid can be placed). HTVS Glide docking module (Schrödinger LLC 2022-3) was used for screening of the ligand set and the top 1000 scoring ligands were re-docked with SP Glide (Schrödinger LLC 2022-3). The top 100 scoring ligands from SP Glide were selected based on chemical similarity (Tanimoto coefficient) using K-means clustering algorithm in the Discovery Informatics module (Schrödinger LLC 2022-3).

For additional docking studies using SP Glide, models of MexY^PA7^ and MexY^PA14^ were prepared as for MexY^PAO1^, above. In total, docking of berberine and identified di-conjugate analogs was performed with four structural models: MexY^PAO1^ (based on PDB 6IOL), MexY^PAO1^ (based on PDB 6TA6), MexY^PA7^ (based on PDB 6IOL), and MexY^PA14^ (based on PDB 6IOL) (**Supplementary Table 2**). Due to the flexibility of Generation 2 and Generation 3 di-berberine conjugate linkers, conventional docking could not sample all potential ligand conformations. Therefore, we used ConfGen module with mixed torsional/ low mode sampling method with 100 steps/ rotatable bond and 1000 iterations (Schrödinger LLC 2022-3) to generate all potential conformers of these ligands. These conformers were subsequently used as the input for SP Glide docking (Schrödinger LLC 2022-3) in MexY^PAO1^, MexY^PA7^ and MexY^PA14^.

### Berberine analog synthesis

#### Instrumentation, chemicals, and general notes

Nuclear magnetic resonance (NMR) spectra were recorded on either a Varian INOVA 600 (600/150 MHz) or Varian INOVA 500 (500/125 MHz) spectrometer. Chemical shifts are quoted in parts per million (ppm) relative to tetramethylsilane standard and with the indicated solvent as an internal reference. The following abbreviations are used to describe signal multiplicities: s (singlet), d (doublet), t (triplet), q (quartet), m (multiplet), br (broad), dd (doublet of doublets), dt (doublet of triplets), etc. Accurate mass spectra were recorded on an Agilent 6520 Accurate-Mass Q-TOF LC/MS.

Non-aqueous reactions were performed under an atmosphere of argon, in flame-dried glassware, with HPLC-grade solvents dried by passage through activated alumina. Brine refers to a saturated aqueous solution of sodium chloride, sat. NaHCO_3_ refers to a saturated aqueous solution of sodium bicarbonate, sat. NH_4_Cl refers to a saturated aqueous solution of ammonium chloride, etc. “Column chromatography,” unless otherwise indicated, refers to purification in a gradient of increasing ethyl acetate concentration in hexanes or methanol concentration in dichloromethane on a Biotage® flash chromatography purification system. All chemicals were used as received from Oakwood, TCI America, Sigma-Aldrich, Alfa Aesar, or CombiBlocks.

#### Experimental procedures and characterization data

Berberrubine chloride (2.4 eq.) was combined with either A. diiodide (1 eq.) or B. dibromide (1 eq.)/sodium iodide (2 eq.) in N,N-dimethylformamide (0.1 M) in a sealed reaction vessel and heated to 70°C. This solution was stirred under a static argon atmosphere until thin layer chromatography (TLC) analysis showed reaction completion (48-72 hours). Solvent was removed *in vacuo* and product was purified via silica gel column chromatography. All synthesized analogs used in biological analysis were purified to >95% purity by high-performance liquid chromatography (HPLC). Additional details of all synthetic procedures and results of chemical analyses are provided in the Supplementary Information.

### Bacterial strains and growth conditions

Strains used in this study (**Supplementary Table 1**) were previously reported *P.* aeruginosa PAO1 (K767), PAO1Δ*mexXY* (K1525), and pan-aminoglycoside resistant clinical isolates K2156 and K2161^14,19,70^. *P. aeruginosa* PA7 and PA14 were kindly provided by Dr. Joanna Goldberg (Department of Pediatrics, Emory University, Georgia, USA). Bacterial strains were maintained on Lysogeny Broth (LB)-Miller or LB-Miller agar (Merck) and grown aerobically at 37°C overnight. Strains were stored in LB-Miller broth supplemented with 16% glycerol at -80°C.

### DNA methods

Chromosomal DNA was extracted from *P. aeruginosa* PAO1 (K767), PAO1ΔmexXY (K1525), or clinical isolates K2156 and K2161 using Qiagen DNeasy Blood and Tissue Kit according to manufacturer protocol. The *mexY* gene from each strain was subsequently amplified using five primers that encompass the 5’ (ATGGCTCGTTTCTTCATT) and 3’ ends (TCAGGCTTGCTCCGTG) of the gene and three internal primers (Int1: GCAGTTCGGCGAAATTCCGC; Int2: GGTTCAACCGCGCCTTC; and, Int3: CCTGGGCATCGACGACATC) to encompass all 3.1 kb. Purified PCR products were sent for Sanger Sequencing (GeneWiz). Resultant FASTA sequences were aligned to *P. aeruginosa* PAO1 (accession GCF_000006765.1) and deposited to NCBI GenBank under the accessions OP868818 (K767), OP868819 (K2156), and OP868820 (K2161). Disruption of *mexXY* in *P. aeruginosa* PAO1Δ*mexXY* (K1525) was validated by colony PCR.

### Antibiotic susceptibility assays

#### Growth conditions

For all antimicrobial susceptibility assays, *P. aeruginosa* strains were grown in cation-adjusted Mueller-Hinton broth (CA-MHB) prepared using premixed MHB (Sigma) supplemented with 20 mg/L calcium chloride (OmniPur) and 10 mg/L magnesium chloride (Sigma). Strains were cultured aerobically, shaking vigorously, at 37°C for 18 hours.

#### Minimum inhibitory concentration (MIC)

All MICs were determined using serial two-fold microdilution method in 96-well microtiter plates as done previously^71^. Antibiotics used in this study were: kanamycin (Kan; OmniPur), amikacin (Ami; ChemImpex), gentamicin (Gen; Sigma), tobramycin (Tob; Alfa Aesar), cefepime (Cef; ChemImpex), ciprofloxacin (Cip; Enzo), tigecycline (Tig; Astatech), imipenem (Imi; Combi-Blocks), and vancomycin (Van; Sigma). MIC values were determined at 18 hours post-inoculation by reading optical density at 600 nm (OD_600_) on a Synergy Neo2 Multi-Mode Plate Reader (Agilent BioTek), and inhibition of growth defined as OD_600_<0.5 above background. All MIC assays were performed at least twice starting from fresh streak plates on separate days and each assay was performed as two technical replicates. Assays were performed for all *P. aeruginosa* strains to determine starting MICs in the absence of berberine or berberine analogs. All MIC assays with berberine ligands (Ber, Ber-C3 ****[5]****, Ber-C12 ****[9]****, or Ber-pAr ****[10]****) were adjusted for a final concentration of 1% DMSO in each well.

#### Checkerboard

Checkerboard assays were performed with *P. aeruginosa* PAO1 (K767) using Kan or Gen, and all berberine ligands. Two-fold serial dilutions for a final concentration of Kan [256-4 µg/mL] or Gen[4-0.06 µg/mL] in CA-MHB were dispensed across the rows of a 96-well microtiter plate. Berberine and berberine analogs were first resuspended in 100% DMSO and then diluted with water to yield stocks [1280-10 µg/mL] in 20% DMSO. Finally, the berberine and berberine analogs were dispensed down the columns of the 96-well microtiter plate to give final two-fold decreasing concentrations (128-1 µg/mL) and 1% DMSO final concentration. *P. aeruginosa* PAO1 (K767) was grown to OD_600_ of 0.1 in CA-MHB and was diluted to a final concentration of 10^5^ cells. MIC values were determined as above at 18 hours post-inoculation by reading OD_600_, where inhibition of growth defined as OD_600_<0.5 above background. All checkerboard assays were performed at least twice starting from fresh streak plates on separate days. OD_600_ readings each replicate between antibiotic and berberine analog interaction were averaged and normalized from 0-1.

For analysis of antibiotic-berberine interactions, fractional inhibitory concentration (FIC) was determined as previously described^72^, where a FIC value < 0.5 indicates synergy between antibiotic and ligand. For berberine and berberine analogs without an experimentally measurable MIC value (i.e., greater than 128 µg/mL), the final OD_600_ values from cultures at 18 hours after exposure to a range of ligand concentrations (128-0 µg/mL) were plotted and nonlinear regression of the log transformed OD_600_ values was used to extrapolate a calculated MIC value.

#### Time-kill

Growth curves for *P. aeruginosa* PAO1 (K767) and PAO1ΔmexXY (K1525) were measured in the absence or presence of amikacin or tobramycin at concentrations ranging from ¼ to 2× the MIC and with select berberine ligands (64 µg/mL). Berberine ligands were prepared and added to 96-well microtiter plate as described above for the checkerboard assays. Microtiter plates containing technical replicates for each antibiotic-berberine pair were incubated for 18 hours with vigorous shaking using in Cytation5 Imaging Reader (Agilent BioTek). OD_600_ values were recorded every 15 minutes and select time points (0, 2, 4, 6, 8, 10, 12, 18, and 24 hours) were plotted. Growth curves were performed in duplicate at least twice from freshly streaked plates on separate days.

#### Three-way synergy

A three-way synergy assay was performed using *P. aeruginosa* PAO1 (K767) in the presence of berberine (64 µg/m), Ber-C3 ****[5]**** (128-1 µg/mL) and Ami (4-0.06 µg/mL). The 96-well microtiter plate for Ber-C3 ****[5]**** and amikacin was set up as a checkerboard synergy assay (as described above). Berberine was added to all wells to a final concentration of 64 µg/mL and the total concentration of berberine and Ber-C3 ****[5]**** was adjusted for a final concentration of 1% DMSO. MIC values were determined as above at 18 hours post-inoculation by reading OD_600._ The assay was performed twice using two fresh streak plates for starter cultures on two different days.

### Hemolysis assay

Hemolysis was performed on mechanically defibrinated sheep blood (Hemostat Labs). One mL of erythrocytes was centrifuged for 10 minutes at 3,800 rpm. Supernatant was discarded, cells were washed in 1 mL of 1× phosphate-buffered saline (PBS), and pelleted. This washing step was repeated five times. The final pellet was diluted 20-fold in 1× PBS. Berberine and berberine ligands were resuspended in 100% DMSO and the final starting concentration in each well was 2.5%. Berberine ligands were two-fold serially diluted in DMSO from 128-0.5 µg/mL in a 96-well microtiter plate. 100 µL of sheep blood suspension was added to 100 µL of berberine ligand, 100 µL 1% Triton-X (positive control), or 100 µL 1× PBS (negative control). Samples were incubated for one hour at 37°C with shaking at 200 rpm, and then pelleted. The OD_540_ of the resulting supernatant was recorded using Synergy Neo2 Multi-Mode Plate Reader (Agilent BioTek). The concentration inducing 20% red blood cell lysis was calculated for each ligand based on positive and negative control absorbances. Hemolysis assays were performed in triplicate.

### MD simulation and MM/GBSA calculation

We used the same strategy and rationale for using a reduced model of *P. aeruginosa* PAO1 MexY-per as previously described for AcrB^73–75^. The MexY-per SP Glide docking model with chemically diverse berberine probes bound to the PBP and DBP binding pocket was used as a starting structure for MD simulations. Simulations were run in five replicates with each run initiated using a different initial velocity with OPLS4 force field, producing essentially identical results. Each system was first neutralized by adding sodium ions around the protein using the System Builder module. The neutralized protein was placed in TIP3P water, and random water molecules were substituted to obtain an ionic strength of 150 mM. Each solvated system was relaxed using a series of restrained minimization stages each of 1 ns duration: 1) all heavy atoms with force constant 50 kcal/molÅ^2^, all protein atoms with 2) 10 kcal/molÅ^2^ and 3) 5 kcal mol kcal/molÅ^2^, and finally 4) with no constraints. Unrestrained MD simulations were then performed for 60 ns, comprising 10 ns for equilibration followed by a 50 ns production run in the isothermal-isobaric (NPT) ensemble using Nose-Hoover chain thermostat and Martyna-Tobias-Klein barostat with relaxation times of 1 and 2 ps, respectively. The equations of motion were integrated using multiple time steps for short-range (2 fs) and long-range (6 fs) interactions with a 10 Å cutoff applied for non-bonded interactions. MM/GBSA calculations were performed in the Prime module (Schrödinger LLC 2022-3) with the OPLS4 force field and allowing protein side chain flexibility within a 5 Å radius of the bound ligand.

## Abbreviations

4,6-DOS: 4,6-deoxystreptamine
CA-MHB: cation-adjusted Mueller Hinton broth
DPB: Distal binding pocket
EPI: efflux pump inhibitor
FIC: fractional inhibitory concentration
HTVS: high-throughput virtual screening
PBP: proximal binding pocket
PEG: polyethylene glycol
RND: Resistance-Nodulation-Division.
*Antibiotics*: Ami: amikacin
Cef: cefepime
Cip: ciprofloxacin
Gen: gentamicin
Kan: kanamycin
Imi: imipenem
Tig: tigecycline
Tob: tobramycin.

## Data Availability

Sequences generated during this analysis are available on NCBI GenBank under accession number BankIt2645695 OP868818-20. Other underlying data are available upon request.

## Acknowledgements

We thank Dr. Keith Poole in the Department of Biomedical and Molecular Sciences, Queens University, Ontario, Canada for providing *P. aeruginosa* PAO1, clinical isolates, and respective *mexXY* knockout strains. We thank Dr. Joanna Goldberg in the Department of Pediatrics, Emory University, Atlanta, Georgia, USA, for providing *P. aeruginosa* PA7 and PA14 isolates. This work was supported by the National Institute of General Medical Sciences (R35 GM119426 to W.M.W), National Institute of General Medical Sciences NRSA Pre-Doctoral Fellowship (F31 GM143891 to L.G.K), Cystic Fibrosis Foundation Student Traineeship (Kavana21H0 to L.G.K), an ACS MEDI Pre-doctoral Fellowship (A.R.M.), and Emory University through an Accelerator Grant from the Biological Discovery through Chemical Innovation (BDCI) initiative (to G.L.C. and W.M.W.). L.G.K. also gratefully acknowledges support from Dr. John and Linda McGowan from the Atlanta Chapter ARCS Foundation.

## Author Contributions

**L.G.K.** and **A.R.M.** designed and performed experiments, analyzed data, and wrote the original draft and edited the manuscript; **D.D.** performed computational analyses; **G.L.C.** and **W.M.W.** conceived the study, supervised the project, and edited the manuscript. All authors jointly discussed the results and implications at all stages of the work and edited the final manuscript. **L.G.K.** and **A.R.M.** are formally recognized co-first authors of this work. Correspondence may be addressed to either **G.L.C.** or **W.M.W.** (co-corresponding authors).

