## Supplemental Information for "Di-berberine conjugates as chemical probes of *Pseudomonas aeruginosa* MexXY-OprM efflux function and inhibition"

**This file contains:**

Supplementary Tables S1-S6

Supplementary Fig. S1-S10

Supplementary Methods: Details of chemical synthesis and compound analysis

Supplementary References

Appendix: $^{1}H$ and $^{13}C$ NMR spectra for all synthesized intermediates and berberine analogs.

### Supplementary Tables

**Table S1. Strains used in this study.**

| Strain | Characteristics | Source (Reference) |
|---|---|---|
| PAO1 (K767) | Wild-type PAO1; lab evolved strain | Poole Lab (Sobel et al. 2003)[1] |
| PAO1Δ*mexXY* (K1525) | K767Δ*mexXY* | Poole Lab (Sobel et al. 2003)[1] |
| K2156 | Pan-aminoglycoside resistant clinical isolate | Poole Lab (Sobel et al. 2003)[1] |
| K2161 | Pan-aminoglycoside resistant clinical isolate | Poole Lab (Sobel et al. 2003)[1] |
| PA14 | Virulent clinical isolate | Goldberg Lab |
| PA7 | MDR clinical isolate | Goldberg Lab |

**Table S2. SP Glide docking scores for berberine and analogs in the *P. aeruginosa* PAO1, PA7, and PA14 MexY binding pocket.**

| Compound | SP Glide Docking score (Average)[a,b] | | | |
|---|---|---|---|---|
| | $MexY^{PAO1}$ | | $MexY^{PA7}$ | $MexY^{PA14}$ |
| | 6IOL (Chain B) | 6TA6 (Chain C) | 6IOL (Chain B) | 6IOL (Chain B) |
| Berberine | -6.46 (-6.04) | -5.70 (-5.38) | -6.41 (-6.13) | -5.75 (-5.01) |
| Ber-Pip [**1**] | -6.03 (-5.82) | -5.82 (-5.20) | -5.74 (-4.41) | -5.02 (-4.94) |
| Ber-Carb [**2**] | -4.73 (-4.68) | -5.75 (-5.75) | -5.41 (-3.37) | -4.23 (-4.00) |
| Ber-Prop [**3**] | -8.10 (-7.97) | -8.36 (-8.01) | -8.17 (-7.99) | -7.87 (-7.44) |
| Ber-C6 [**4**] | -8.99 (-7.39) | -9.32 (-8.79) | -9.33 (-7.74) | -8.10 (-6.72) |
| Ber-C3 [**5**] | -9.18 (-7.74) | -9.28 (-8.97) | -9.77 (-7.47) | -8.37 (-5.85) |
| Ber-C4 [**6**] | -8.88 (-7.51) | -9.50 (-9.11) | -9.06 (-6.68) | -8.09 (-6.21) |
| Ber-C8 [**7**] | -9.28 (-7.38) | -9.32 (-8.74) | -10.30 (-8.37) | -8.48 (-6.91) |
| Ber-C10 [**8**] | -9.55 (-7.35) | -9.77 (-9.13) | -9.73 (-8.23) | -8.63 (-7.22) |
| Ber-C12 [**9**] | -9.77 (-7.14) | -9.53 (-8.71) | -10.10 (-7.80) | -8.45 (-6.99) |
| Ber-pAr [**10**] | -9.37 (-7.67) | -9.56 (-9.29) | -10.01 (-6.47) | -8.60 (-6.96) |
| Ber-Biph [**11**] | -10.36 (-10.39) | -10.26 (-9.65) | -11.08 (-10.23) | -10.56 (-10.26) |
| Ber-PEG5 [**12**] | -9.73 (-7.76) | -9.43 (-9.13) | -9.87 (-7.52) | -8.70 (-6.80) |
| Ber-PEG8 [**13**] | -9.90 (-7.89) | -9.75 (-9.36) | -10.10 (-7.72) | -8.94 (-7.47) |

[a]Docking scores correspond to the top scored pose (shown in figures, where applicable) and the average of all docking poses for each compound in parenthesis.
[b]The PDB code noted (with chain ID in parenthesis) indicates to the structure used to generate the $MexY^{PAO1}$, $MexY^{PA7}$, and $MexY^{PA14}$ homology models used for docking studies (see Methods for details).

**Table S3. Contact residues for berberine and analogs docked in the PBP/ DBP of MexY from *P. aeruginosa* strains PAO1, PA7, and PA14 using SP Glide.**

| Ligand | PAO1 | PA7 | PA14 |
|---|---|---|---|
| Berberine | R166, W177, F610 | Y127, F610 | Q175, K291 |
| Ber-Pip [**1**] | Y127 | Y127, K173 | S88, K291 |
| Ber-Carb [**2**] | K173, K291 | K173, K291 | K173 |
| Ber-Prop [**3**] | D124 | D124 | D133,K291,Y326 |
| Ber-C6 [**4**] | T176, E273, Y752, K764 | T176, K273, Y752, K764 | S46, Y127, S272, E273, K291, K764 |
| Ber-C3 [**5**] | S46, Y127, T176, S272, E273, K291 | Y127, E175, K291 | N44, D133, K291 |
| Ber-C4 [**6**] | E175 | Y127, F276 | N44, Y127, K291 |
| Ber-C8 [**7**] | S46, K79 | Y127, K173, T176, K291 | R116, S272, E273, K291 |
| Ber-C10 [**8**] | E175, K291 | E175, E273, K764 | N44, S272, E273, K764 |
| Ber-C12 [**9**] | Y127, K291 | E175, K291, K764 | E273, K291 |
| Ber-pAr [**10**] | Y127, K173, S671 | K173, E175, K291 | S46 |
| Ber-Biph [**11**] | K173, E175, E273, K291 | K173, E175, E273, K291 | K173, K291 |
| Ber-PEG5 [**12**] | Y127, S671 | Y127, S671 | K173, Q175, K291 |
| Ber-PEG8 [**13**] | Y127, E175 | S88, Y127,F276, K291, S671 | S46, S88, Y127, D133, Q175, E273 |

**Table S4. Predicted minimum inhibitory concentration (MIC) and fractional inhibitory concentration (FIC) for berberine and analogs.**

| | Compound | Predicted MIC (μg/mL)[a] | Antibiotic[b] | FIC | |
|---|---|---|---|---|---|
| | | | | Range | Average |
| Generation 1 | Berberine | 979 | Kan | 0.50 | 0.50 |
| | | | Gen | 0.50 | 0.50 |
| | Ber-Pip [**1**] | 3549 | Kan | 0.50 | 0.50 |
| | | | Gen | 1.14 | 1.14 |
| | Ber-Carb [**2**] | 760 | Kan | 1.67 | 1.67 |
| | | | Gen | 0.52 | 0.52 |
| | Ber-Prop [**3**] | 947 | Kan | 0.39-0.50 | 0.44 |
| | | | Gen | 0.39-0.50 | 0.44 |
| | Ber-C6 [**4**] | 287 | Kan | 0.28-0.51 | 0.38 |
| | | | Gen | 0.36-0.51 | 0.48 |
| Generation 2 | Ber-C3 [**5**] | 584 | Kan | 0.30-0.50 | 0.40 |
| | | | Gen | 0.30-0.50 | 0.38 |
| | Ber-C4 [**6**] | 508 | Kan | 0.38-0.5 | 0.44 |
| | | | Gen | 0.38-0.5 | 0.44 |
| | Ber-C8 [**7**] | 250 | Kan | 0.28-0.5 | 0.39 |
| | | | Gen | 0.38-0.5 | 0.40 |
| | Ber-C10 [**8**] | 200 | Kan | 0.17-0.51 | 0.31 |
| | | | Gen | 0.17-0.51 | 0.33 |
| | Ber-C12 [**9**] | 180 | Kan | 0.26-0.51 | 0.36 |
| | | | Gen | 0.26-0.77 | 0.46 |
| Generation 3 | Ber-pAr [**10**] | 400 | Kan | 0.52-0.57 | 0.55 |
| | | | Gen | 0.33-0.52 | 0.43 |
| | Ber-Biph [**11**] | 502 | Kan | 0.50-0.51 | 0.51 |
| | | | Gen | 0.52 | 0.52 |
| | Ber-PEG5 [**12**] | 545 | Kan | 0.56 | 0.56 |
| | | | Gen | 0.48-0.53 | 0.51 |
| | Ber-PEG8 [**13**] | 622 | Kan | 0.55 | 0.55 |
| | | | Gen | 0.46-0.53 | 0.49 |

[a]Predicted berberine analog MIC was determined by plotting final $OD_{600}$ values from cultures at 18 hours after exposure to a range of ligand concentrations (128-0 μg/mL) and using nonlinear regression of the log transformed $OD_{600}$ values to obtain the MIC value (y intercept = 0).
[b]Aminoglycosides: kanamycin (Kan) and gentamicin (Gen).

**Table S5. Three-way synergy assay with berberine, Ber-C3 [5], and amikacin.**

| Berberine (μg/mL) | Ber-C3 [**5**] (μg/mL) | Ami MIC (μg/mL) |
|---|---|---|
| 0 | 0 | 1 |
| | 64 | 0.25 |
| 64 | 0 | 0.5 |
| | 1 | 1 |
| | 2 | 1 |
| | 4 | 1 |
| | 8 | 1 |
| | 16 | 1 |
| | 32 | 0.5 |
| | 64 | 0.5 |
| | 128 | 0.25 |

**Table S6. Contact residues for berberine and select analogs in the PBP and DBP of *P. aeruginosa* PAO1 MexY before and after MD simulation.**

| Ligand | Before MD (0 ns)[a] | After MD (50 ns)[b] |
|---|---|---|
| Berberine | - | F610 |
| Ber-C3 [**5**] | R116, Y127, K173, E175, K291, F610 | R67, D124, **E175**, F276, **K291**, **F610**, S671, K764 |
| Ber-C12 [**9**] | Y127, Q163, R763 | K291, Y326 |
| Ber-pAr [**10**] | R166, E273, F276, F610, S671, Q761 | R67, **F276**, K291, **F610**, **S671** |

[a]The predicted interactions noted are after the initial equilibration (10 ns) period immediately prior to the 50 ns MD production run, and may therefore differ from initial docking (**Table S1**).
[b]Interactions with residues in bold are present before and after the MD simulation.

### Supplemental Figures

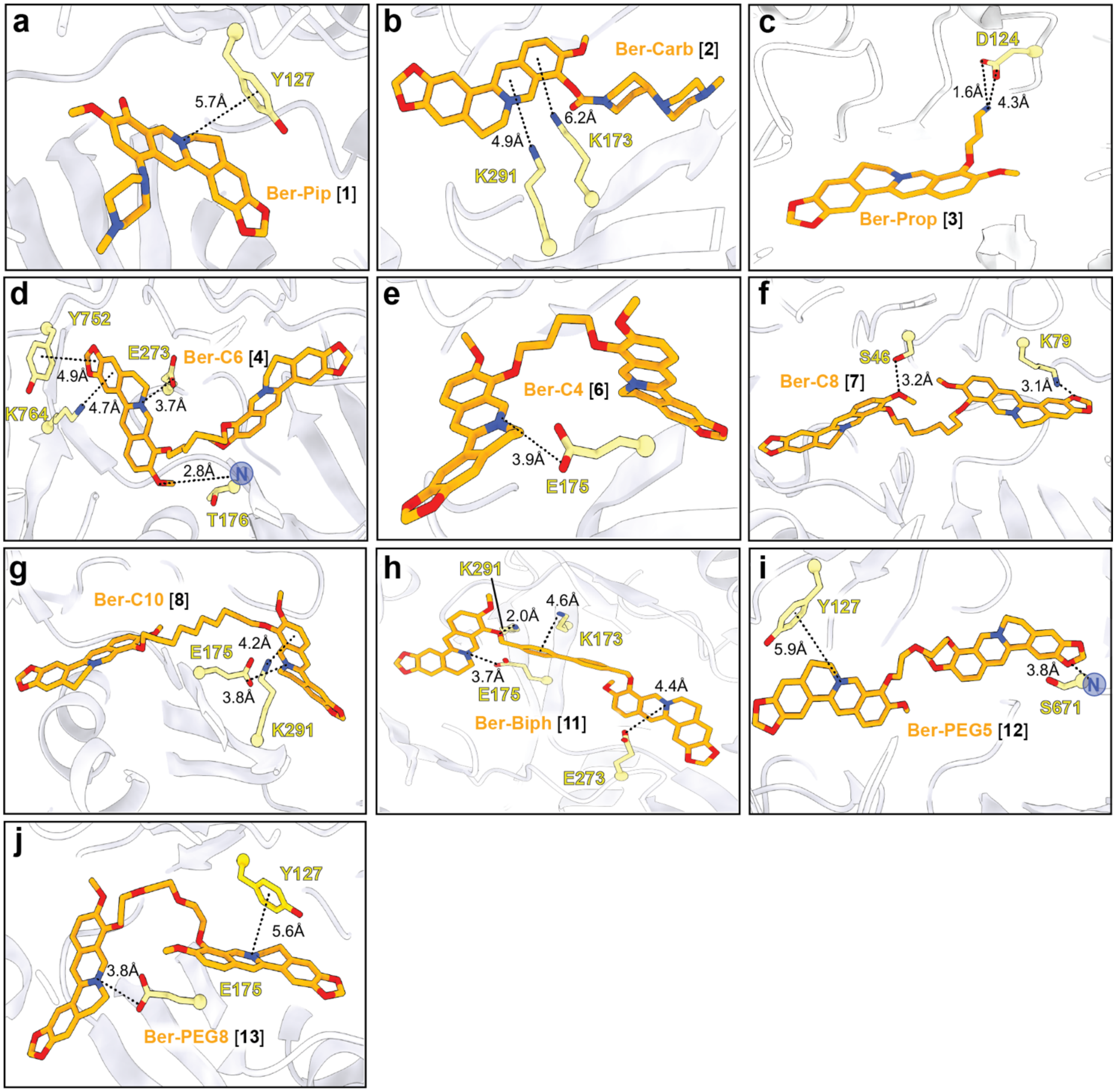


**Supplemental Fig. S1. Lowest energy docking poses of berberine analogs in the MexY$^{PAO1}$ binding pocket.** Docking of all berberine analogs (orange) not included in **Fig.2**. *Generation 1* analogs: **(a)** Ber-Pip [**1**], **(b)** Ber-Carb [**2**], **(c)** Ber-Prop [**3**], **(d)** Ber-C6 [**4**]. All *Generation 1* analogs docked preferentially in the MexY$^{PAO1}$ DBP, with Ber-C6 [**4**] docked in the DBP at the PC1/DN interface. *Generation 2* analogs: **(e)** Ber-C4 [**6**], **(f)** Ber-C8 [**7**], and **(g)** Ber-C10 [**8**]. Ber-C4 [**6**] and Ber-C10 [**8**] showed preference for the DBP. In contrast, Ber-C8 [**7**] showed preference for the PBP. *Generation 3* analogs**:(h)** Ber-Biph [**11**], **(i)** Ber-PEG5 [**12**], and **(j)** Ber-PEG8 [**13**]. Ber-Biph [**11**] and Ber-PEG8 [**13**] showed preference for the DBP, while Ber-PEG5 [**12**] spanned the PBP to DBP.

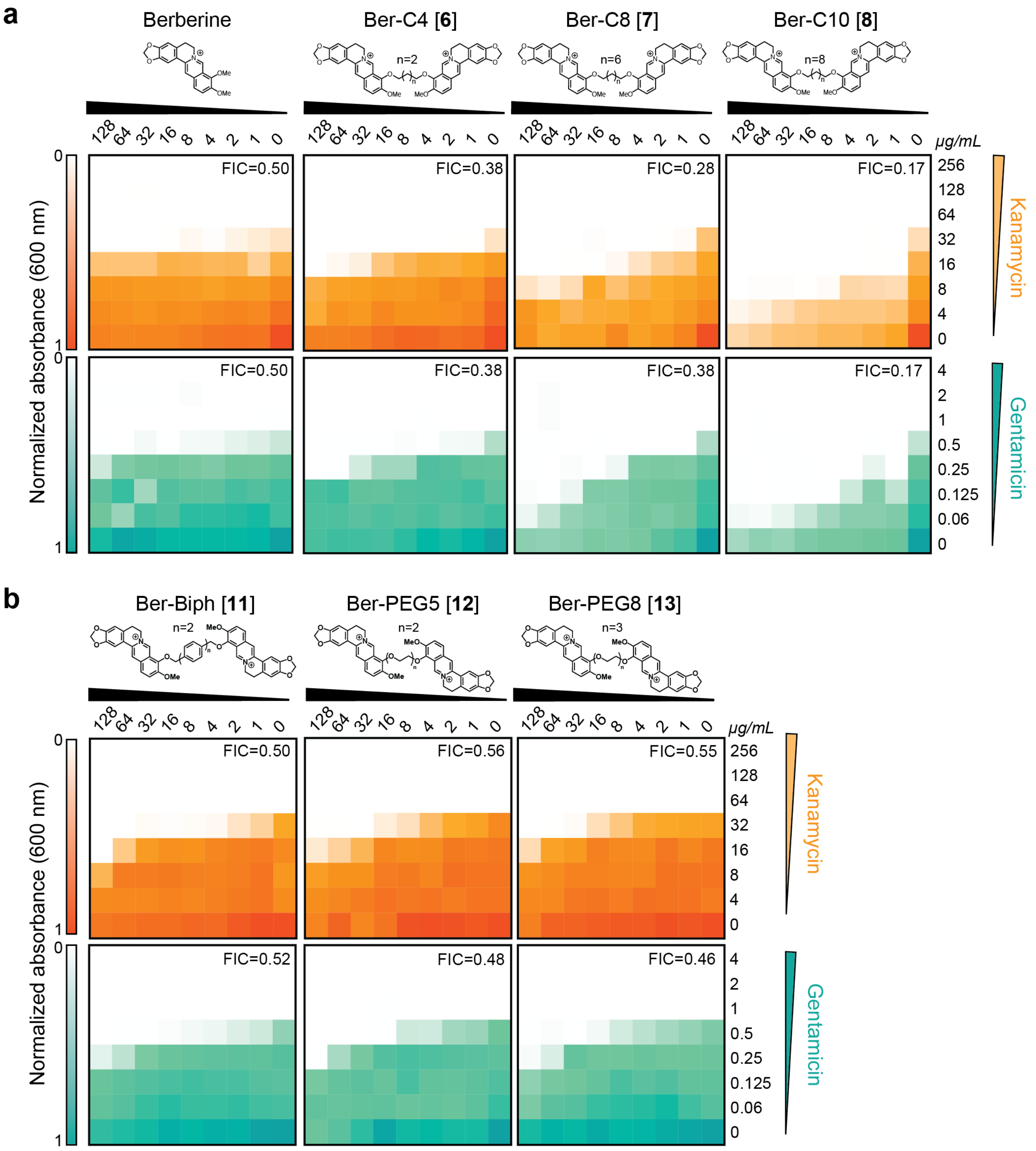


**Supplemental Fig. S2. Checkerboard synergy assays for additional di-berberine conjugates.** Checkerboard synergy assays in *P. aeruginosa* PAO1 with selected analogs (those not shown in **Fig. 1** and **Fig. 3**): **(a)** berberine (same as shown in other figures but included here for ease of comparison) and *Generation 2* ligands Ber-C4 [**6**], Ber-C8 [**7**], and Ber-C10 [**8**]. Increased synergy compared to berberine is observed for all tested compounds with both Kan (top, orange) and Gen (bottom, teal). **(b)** *Generation 3* compounds Ber-Biph [**11**], Ber-PEG5 [**12**], and Ber-PEG8 [**13**], which all showed similar synergy with Kan (top, orange) and Gen (bottom, teal) as berberine. All data are shown as the normalized mean of the optical density ($OD_{600}$) of two biological replicates, (0 is no growth, 1 is maximum growth). The lowest FIC score for each compound-antibiotic pairing is given in the upper, right corner.

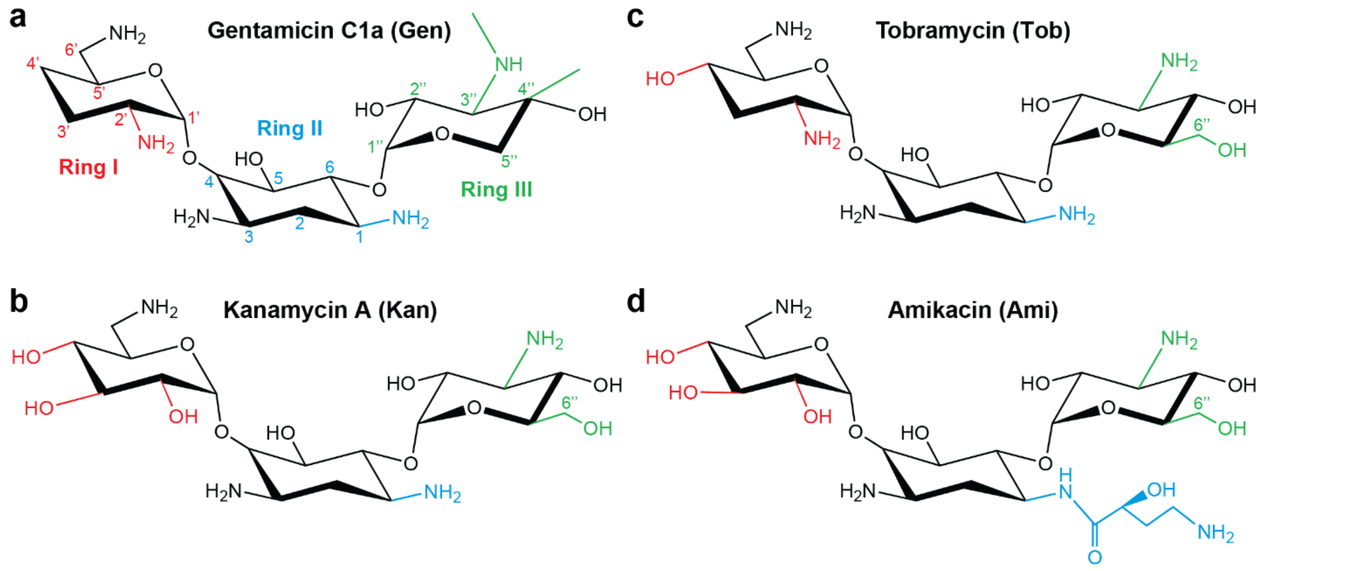


**Supplemental Fig. S3. Structures of aminoglycosides used in this study**. Chemical structures of (**a**) gentamicin C1a (Gen), (**b**) kanamycin A (Kan), (**c**) tobramycin (Tob), and (**d**) amikacin (Ami), highlighting the differences between the four structures color coded by ring. Ring and carbon atom numbers within each aminoglycoside are indicated on the structure of gentamicin (except Ring III 6'' which is only present in the three other aminoglycosides). The deoxystreptamine ring (Ring II) is appended with a (S)-4-amino-2-hydroxubutyrate group on the amino group of position 1 (commonly referred to as L-HABA, or, L-hydroxyaminobuteroyl amide).

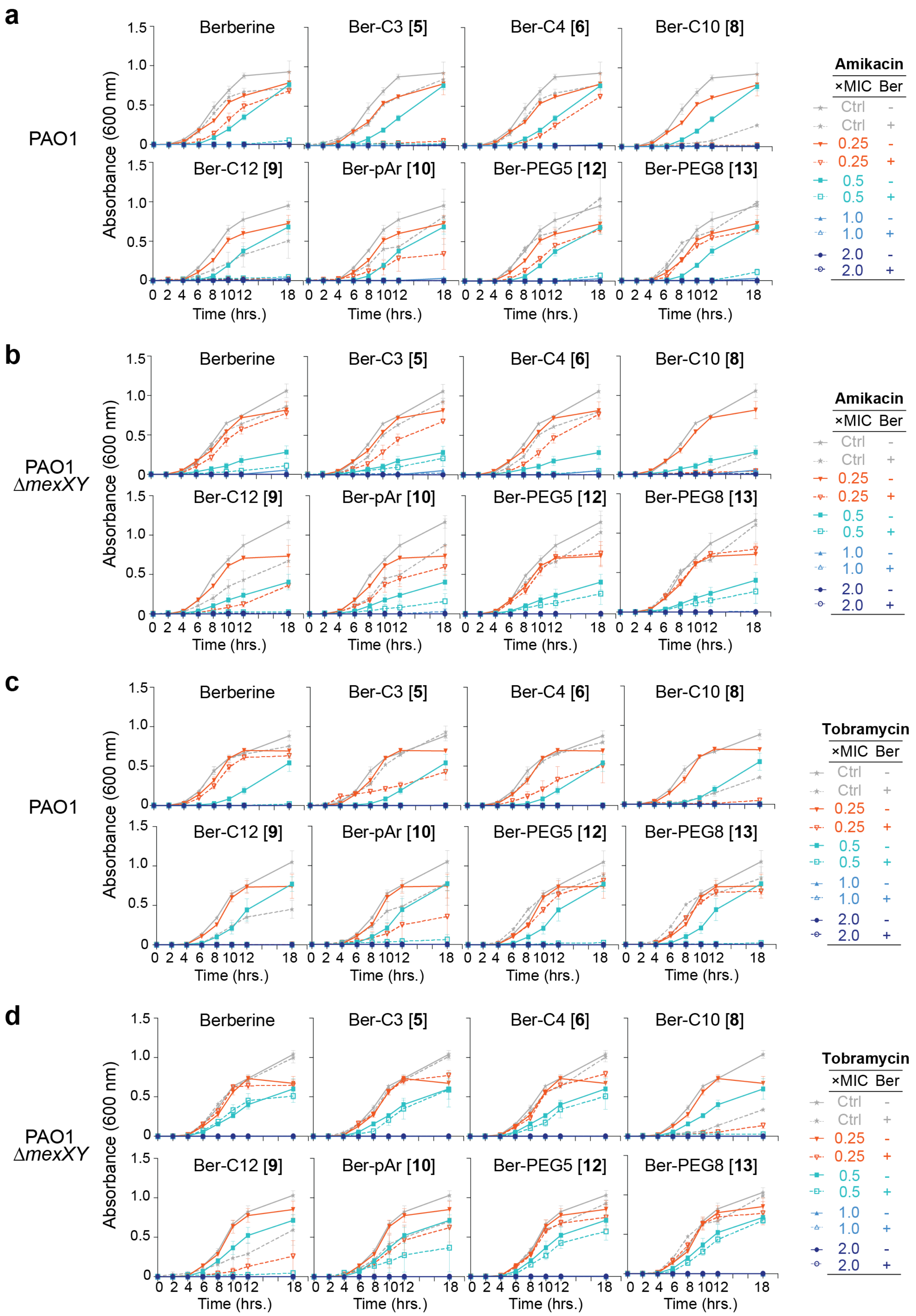


**Supplemental Fig. S4.** (See next page for full legend.)

**Supplemental Fig. S4. Growth curves with berberine and select analogs with amikacin and tobramycin in *P. aeruginosa* PAO1 and isogenic PAO1Δ*mexXY*.** Time-kill assays for berberine and analogs from *Generation 2* (Ber-C3 [**5**], Ber-C4 [**6**], Ber-C10 [**8**], and Ber-C12 [**9**]) and *Generation 3* (Ber-pAr [**10**], Ber-PEG5 [**12**], and Ber-PEG8 [**13**]) at 64 µg/mL are shown with (**a,b)** Ami and **(c,d)** Tob over 18 hours in both *P. aeruginosa* PAO1 and its corresponding *mexXY* isogenic knockout (PAOΔ*mexXY*). Data are shown as the average $OD_{600}$ of biological replicates, each with two technical replicates; error bars are standard deviation.

**a**

| Compound | RBC Hemolysis |
|---|---|
| Berberine | >128 µg/mL |
| Ber-C4 [**6**] | >128 µg/mL |
| Ber-C10 [**8**] | >128 µg/mL |
| Ber-C12 [**9**] | >128 µg/mL |
| Ber-pAr [**10**] | >128 µg/mL |
| Triton-X (+) | <1% |

**b**

| Conc. (µg/mL) | PAO1 | | PAO1Δ*mexXY* | |
|---|---|---|---|---|
| Ber-C12 [**9**] | 0 | 64 | 0 | 64 |
| Vancomycin | 128 | 128 | 128 | 128 |

**c**

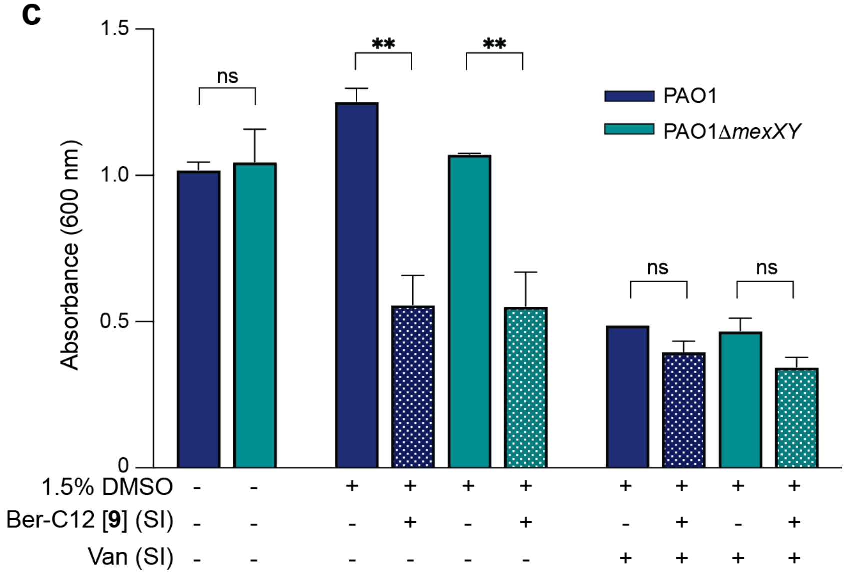


**Supplemental Fig. S5. Berberine and di-berberine conjugates do not disrupt *P. aeruginosa* membrane. (a)** Hemolysis assay was performed with sheep red blood cells in the presence of 128-0 µg/mL of berberine and berberine analogs (Ber-C4 [**6**], Ber-C10 [**8**], Ber-C12 [**9**], and Ber-pAr [**10**]). The reported concentration of berberine analog or Triton-X (positive control) is that needed to lyse ≥ 20% of red blood cell as determined by absorbance measurement ($OD_{540}$). Berberine and analogs did not lyse ≥ 20% of red blood cells at any concentration tested (up to 128 µg/mL). The average from triplicate assays are reported. **(b)** The vancomycin (Van) MIC was determined in *P. aeruginosa* PAO1 and *P. aeruginosa* PAO1Δ*mexXY* in the absence and presence of 64 µg/mL of Ber-C12 [**9**] and did not change under any conditions tested. The assay was performed in duplicate. **(c)** Final absorbance ($OD_{600}$) for PAO1 (blue) and PAO1Δ*mexXY* (teal) in the absence (filled bar) and presence (stippled bar) of 64 µg/mL Ber-C12 [**9**] and a sub-inhibitory concentration (SI) of Van (64 µg/mL). Data are plotted as average and error bars represent the standard deviation. Statistical significance was determined by unpaired T-test between each condition (* $p \leq 0.05$, ** $p \leq 0.01$, *** $p \leq 0.001$, and **** $p \leq 0.0001$). A difference was observed in the final absorbance for both PAO1 and PAO1Δ*mexXY* in the presence of 64 µg/mL, but this difference was not apparent upon the addition of SI of Van.

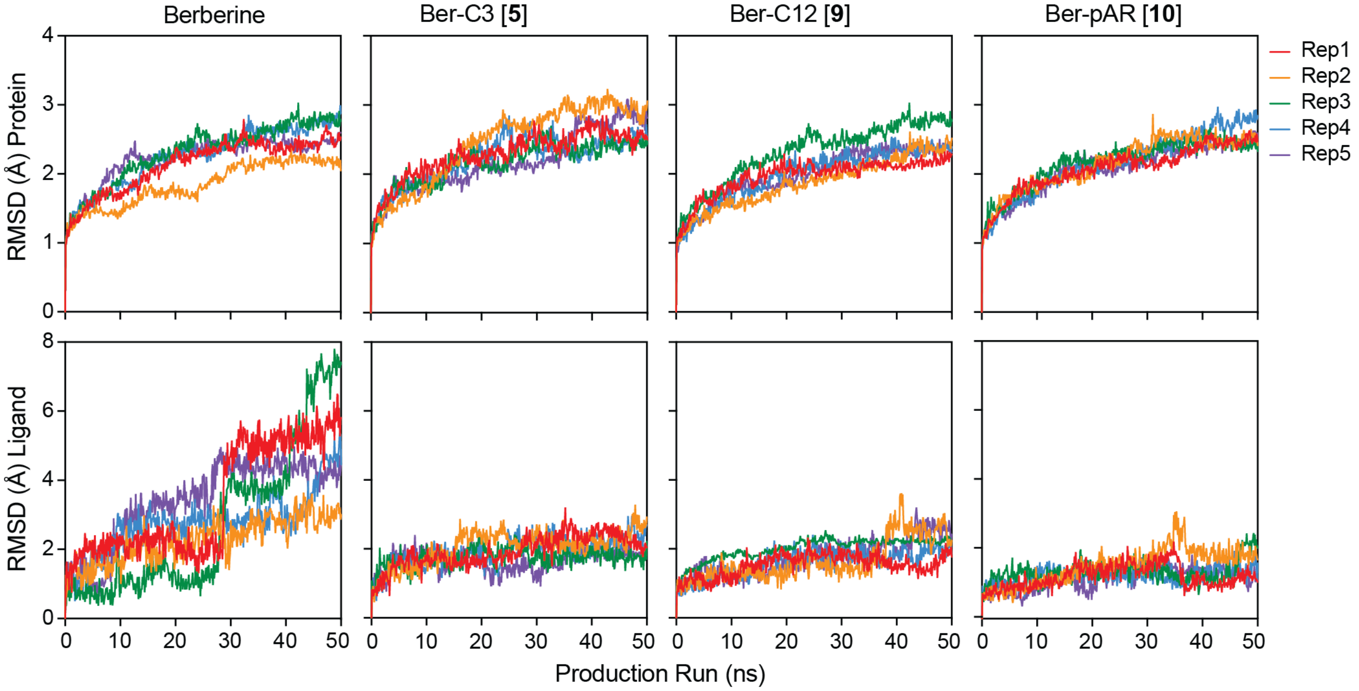


**Supplemental Fig. S6. Protein and ligand RMSD plots from replicate MD similations of MexY with berberine and select analogs.** RMSD during the five replicate 50 ns MD simulations (Rep1-5) for the protein (top)-ligand (bottom) pairs indicated above each pair of plots: berberin, Ber-C3 [**5**], Ber-C12 [**9**], and Ber-pAR [**10**].

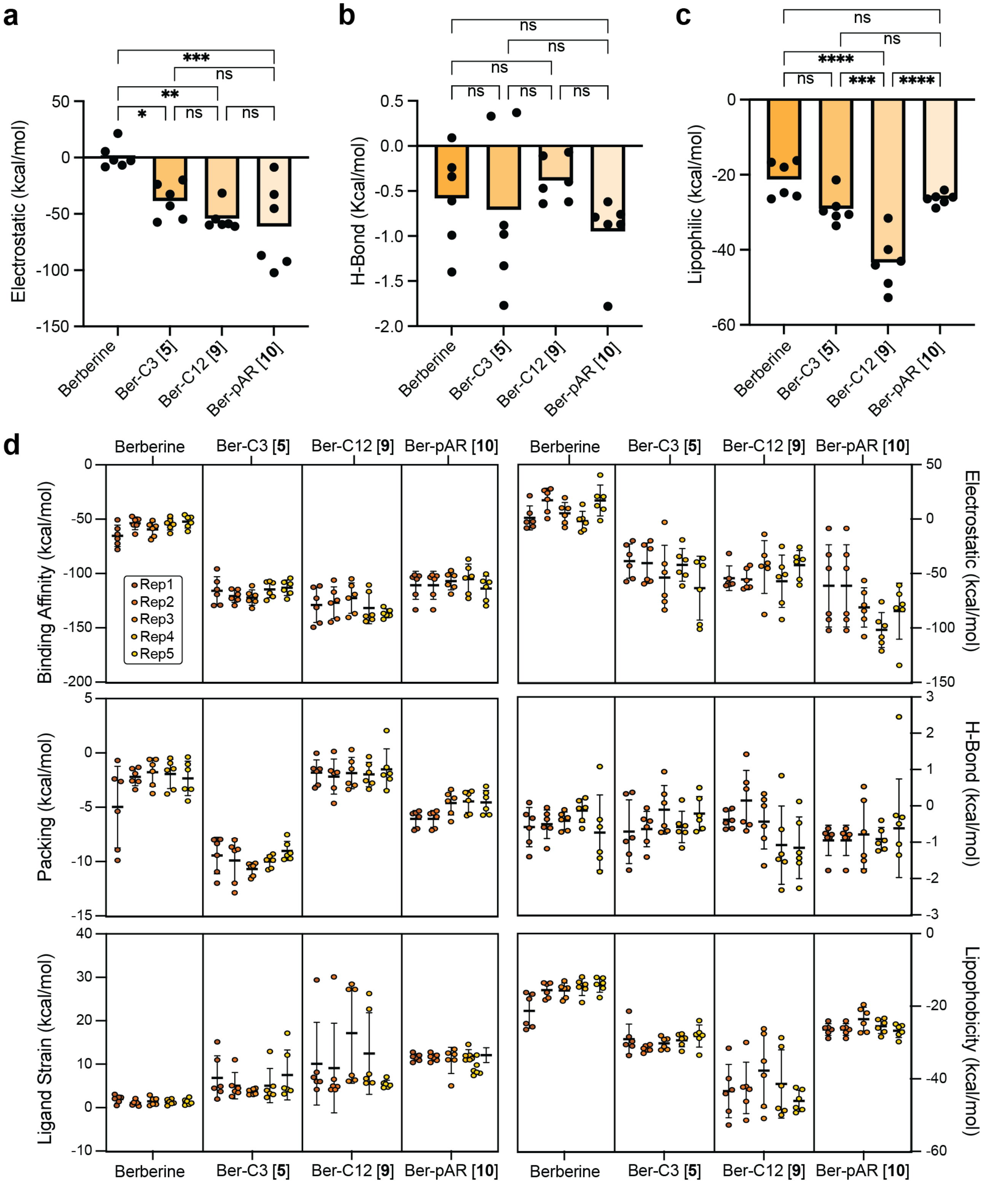


**Supplemental Fig. S7. MM/GBSA calculations from replicate MD simulations.** Additional parameters derived from MM/GBSA calculations for MD Rep1 (also see **Fig.4**): **(a)** electrostatic interaction, **(b)** hydrogen bonding (H-bond), and **(c)** lipophilic interaction. Data shown are the individual calculated values for six frames (one per 10 ns from 0-50 ns) with the average shown as a bar plot. Statistical analysis was performed using one-way ANOVA on the mean for each ligand and a post-hoc Turkey test was used for pairwise comparison. Statistically significant results are shown (* p≤0.05, ** p≤0.01, *** p≤0.001, and **** p≤0.0001). **(d)** Summary of all MM/GBSA calculations for the five replicate simulations (Rep1-5) with berberine and top di-berberine conjugates. Data shown are the individual calculated values for six frames as for panels (a)-(c).

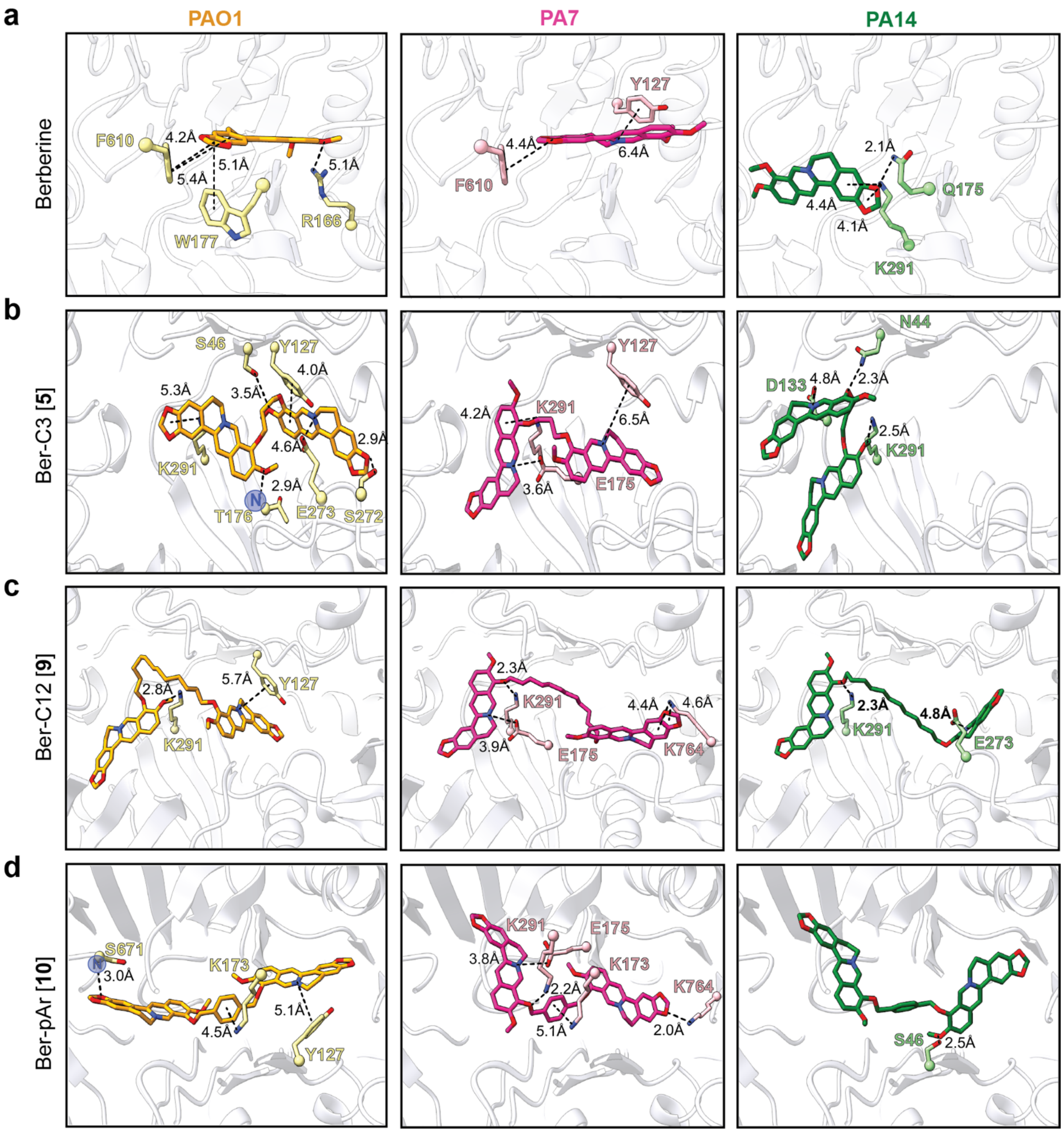


**Supplemental Fig. S8. Preferred docking conformation for berberine and select analogs in the binding pockets of MexY$^{PAO1}$, MexY$^{PA7}$, and MexY$^{PA14}$.** Docking of **(a)** berberine, and di-berberine conjugate analogs **(b)** Ber-C3 [**5**], **(c)** Ber-C12 [**9**], and **(d)** Ber-pAr [**10**] in the PBP/DBP of MexY$^{PAO1}$ (orange, left), MexY$^{PA7}$ (pink, middle), and MexY$^{PA14}$ (green, right). Berberine and Ber-C3 [**5**] adopt similar conformations in MexY$^{PAO1}$/ MexY$^{PA7}$. In contrast, Ber-C12 [**9**] and Ber-pAr [**10**] adopt similar conformations in MexY$^{PA7}$/ MexY$^{PA14}$.

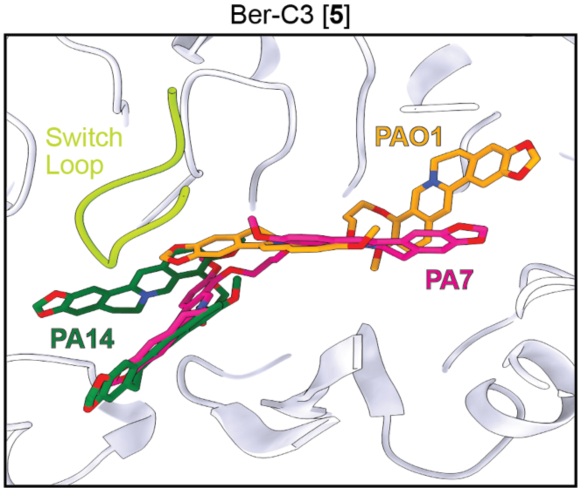


**Supplemental Fig. S9. Distinct predicted docking conformations of Ber-C3 [5] in the binding pocket of MexY from different *P. aeruginosa* strains.** Predicted docking shows that Ber-C3 [**5**] preferentially binds in the DBP of MexY in an 'extended' conformation in MexY$^{PAO1}$ (orange) and MexY$^{PA7}$ (pink), but in a 'compact' conformation in MexY$^{PA14}$ (green).

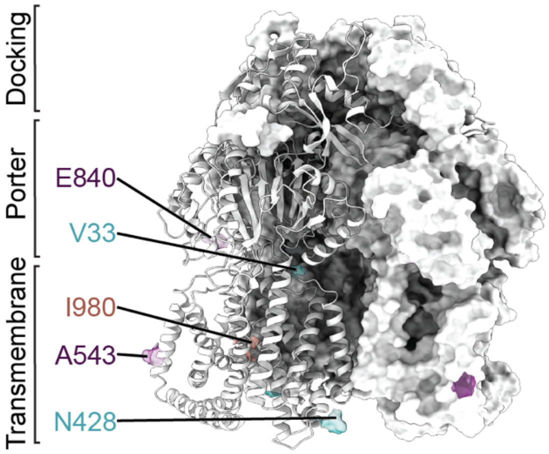


**Supplemental Fig. S10. Unique amino acid changes in pan-aminoglycoside resistant *P. aeruginosa* clinical isolate K2161.** Unique residues identified from alignment of MexY sequences from *P. aeruginosa* PAO1 and clinical isolates K2156 (salmon) and K2161 (teal) mapped on a cartoon of MexY$^{PAO1}$ in the binding state. Residues with the same amino acid substitution in both K2156 and K2161 are shown in purple. The three domains of the MexY transporter (docking, porter, and transmembrane) are indicated.

### Supplementary Methods: Details of chemical synthesis and compound analysis

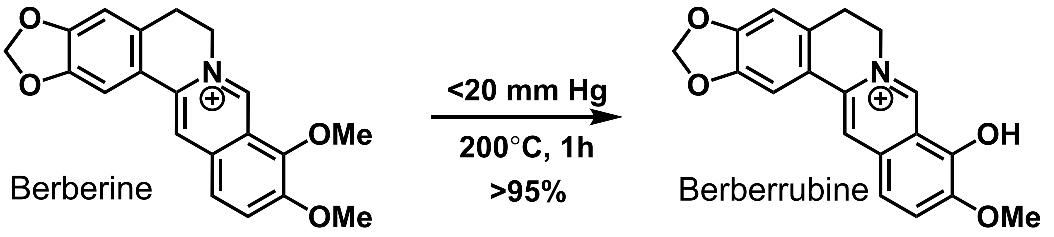


**Berberrubine chloride.**[2] Berberine chloride (500 mg, 1.34 mmol, 1 eq.) was heated to 200°C under vacuum (<20 mm Hg) for 45 minutes with stirring, causing a color change from yellow to red. Solid was filtered, washing with chloroform. Filtrate was purified by silica column chromatography (0-10% MeOH in DCM) to yield the title compound as a dark red solid (480 mg, >95% yield). **$^1$H NMR** (600 MHz, Chloroform-*d*) δ 9.16 (s, 1H), 7.54 (s, 1H), 7.21 (d, *J* = 7.9 Hz, 1H), 7.20 (s, 1H), 6.71 (s, 1H), 6.46 (d, *J* = 7.9 Hz, 1H), 6.02 (s, 2H), 4.36 (s, 2H), 3.87 (s, 3H), 3.08 – 3.02 (m, 2H). **$^{13}$C NMR** (151 MHz, $CDCl_3$) δ 167.92, 150.81, 148.99, 148.07, 145.61, 132.90, 131.27, 128.08, 122.16, 120.36, 120.01, 117.48, 108.32, 104.47, 102.84, 101.77, 56.03, 53.19, 28.57. **HRMS** Accurate mass (ES+): Found 322.10777 (+1.21 ppm), $C_{19}H_{16}NO_4$ ($M^+$) requires 322.10738.

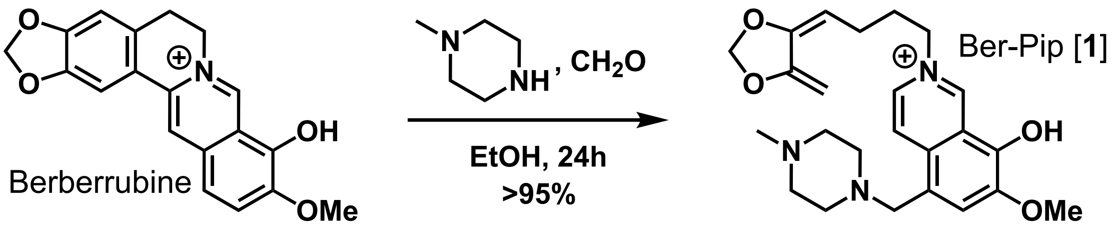


**Ber-Pip [1]: 9-hydroxy-10-methoxy-12-((4-methylpiperazin-1-yl)methyl)-5,6-dihydro-[1,3]dioxolo[4,5-g]isoquinolino[3,2-a]isoquinolin-7-ium chloride.**[3] To a solution of berberrubine (24 mg, 0.067 mmol, 1 eq.) in anhydrous EtOH (1 mL) was added N-methylpiperazine (0.037 mL, 0.335 mmol, 5 eq.) and formaldehyde (37% aq., 0.035 mL, 0.335 mmol, 5 eq.). The solution stirred at room temperature for 24 hours and was then concentrated *in vacuo* and purified by silica column chromatography (0-5% MeOH/DCM) to afford the title compound as a yellow powder (30.5 mg, >95% yield). **$^1$H NMR** (600 MHz, $CDCl_3$) δ 9.21 (s, 1H), 8.03 (s, 1H), 7.21 (s, 1H), 7.18 (s, 1H), 6.70 (s, 1H), 6.02 (s, 2H), 4.38 (t, *J* = 5.9 Hz, 2H), 3.86 (s, 3H), 3.65 (s, 2H), 3.04 (t, *J* = 6.0 Hz, 2H), 2.52 (s, 8H), 2.29 (s, 3H). **$^{13}$C NMR** (151 MHz, $CDCl_3$) δ 167.15, 149.90, 149.15, 148.21, 145.75, 132.96, 129.71, 128.33, 123.82, 122.55, 120.47, 115.07, 110.00, 108.45, 104.71, 101.87, 64.39, 59.75, 56.33, 55.19, 53.30, 52.68, 45.81, 28.64, 25.49. **HRMS** Accurate mass (ES+): Found 434.20693 (-1.17 ppm), $C_{25}H_{28}N_3O_4$ ($M^+$) requires 434.20743.

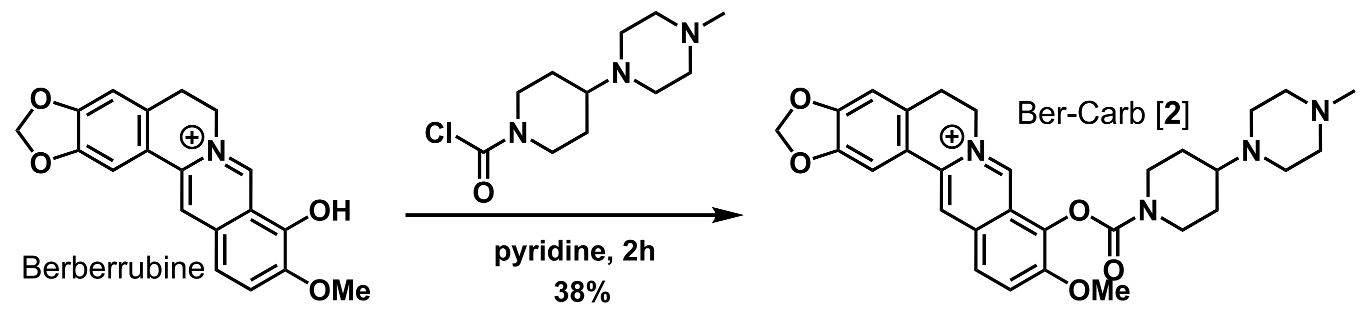


**Ber-Carb [2]: 10-methoxy-9-((4-(4-methylpiperazin-1-yl)piperidine-1-carbonyl)oxy)-5,6-dihydro-[1,3]dioxolo[4,5-g]isoquinolino[3,2-a]isoquinolin-7-ium chloride formate.**[4] Berberrubine (96.4 mg, 0.300 mmol, 1 eq.) was dissolved in pyridine (1 mL) and added in one portion to 4-(4-methylpiperazin-1-yl)piperidine-1-carbonyl chloride (369 mg, 1.50 mmol, 5 eq.)**.** The solution was stirred at room temperature for 2 hours, then was concentrated *in vacuo* and purified by silica column chromatography (0-5% MeOH/DCM) to yield the title compound as a yellow solid (64.6 mg, 38% yield) **$^1$H NMR** (600 MHz, $D_2O$) δ 9.24 (s, 1H), 8.41 (s, 1H), 8.15 (s, 1H), 7.84 (d, *J* = 19.0 Hz, 2H), 7.14 (s, 1H), 6.80 (s, 1H), 6.00 (s, 2H), 4.65 (s, 2H), 4.40 (s, 1H), 4.11 (d, *J* = 12.9 Hz, 1H), 3.92 (s, 3H), 3.68 – 3.52 (m, 2H), 3.22 – 3.16 (m, 1H), 3.09 – 2.97 (m, 4H), 2.62 (br s, 4H), 2.27 (s, 4H), 2.05 (br s, 2H), 1.56 (d, *J* = 8.5 Hz, 1H), 1.47 (d, *J* = 10.4 Hz, 1H). **$^{13}$C NMR** (151 MHz, $D_2O$) δ 171.02, 153.34, 150.46, 150.15, 147.64, 142.69, 138.04, 133.70, 132.89, 130.14, 126.43, 125.25, 121.36, 119.60, 108.28, 104.98, 102.28, 60.58, 56.89, 55.97, 53.54, 47.84, 27.57, 27.23, 26.40. **HRMS** Accurate mass (ES+): Found 531.26012 (-0.14 ppm), $C_{30}H_{35}N_4O_5$ ($M^+$) requires 531.26020.

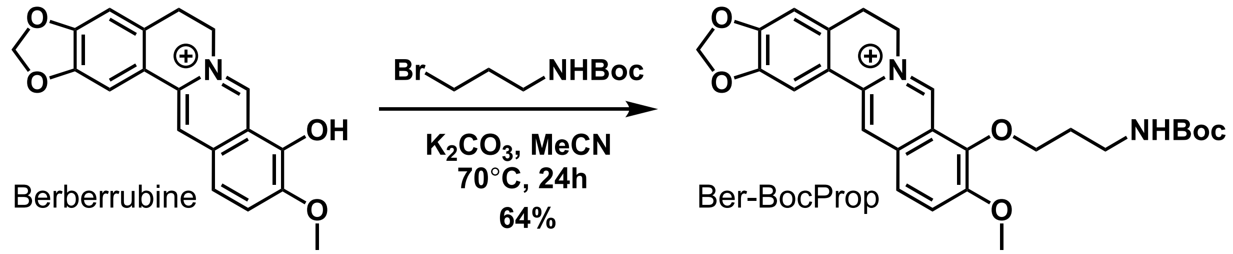


**9-(3-((tert-butoxycarbonyl)amino)propoxy)-10-methoxy-5,6-dihydro-[1,3]dioxolo[4,5-g]isoquinolino[3,2-a]isoquinolin-7-ium chloride.** To a solution of berberrubine (50 mg, 0.134 mmol, 1 eq.) and $K_2CO_3$ (18.6 mg, 0.403 mmol, 3 eq.) in acetonitrile (1 mL) preheated to 70°C was added tert-butyl (3-bromopropyl)carbamate (64 mg, 0.269 mmol, 2 eq.). Additional acetonitrile (0.5 mL) was added, the vessel was sealed, and the reaction continued to stir at 70°C for 24 hours. The crude reaction mixture was cooled to RT and purified by silica column chromatography (0-10% MeOH/DCM), yielding the title compound as a yellow powder (45.9 mg, 64%) **$^1$H NMR** (500 MHz, DMSO-$d_6$) δ 9.83 (s, 1H), 8.93 (s, 1H), 8.19 (d, *J* = 9.2 Hz, 1H), 7.97 (d, *J* = 9.0 Hz, 1H), 7.79 (s, 1H), 7.09 (s, 1H), 6.97 (t, *J* = 5.7 Hz, 1H), 6.16 (s, 2H), 5.00 – 4.89 (m, 2H), 4.28 (t, *J* = 6.0 Hz, 2H), 4.04 (s, 3H), 3.24 – 3.15 (m, 4H), 1.94 (p, *J* = 5.9, 5.5 Hz, 2H), 1.34 (s, 9H). **$^{13}$C NMR** (151 MHz, DMSO-$d_6$) δ 155.87, 150.28, 149.80, 147.67, 145.43, 142.73, 137.37, 132.98, 130.60, 126.63, 123.34, 121.53, 120.43, 120.16, 108.43, 105.43, 102.10, 77.65, 71.80, 57.06, 55.30, 54.95, 36.68, 30.28, 28.25, 26.34. **HRMS** Accurate mass (ES+): Found 479.21776 (-0.14 ppm), $C_{27}H_{31}N_2O_6$ ($M^+$) requires 479.21766.

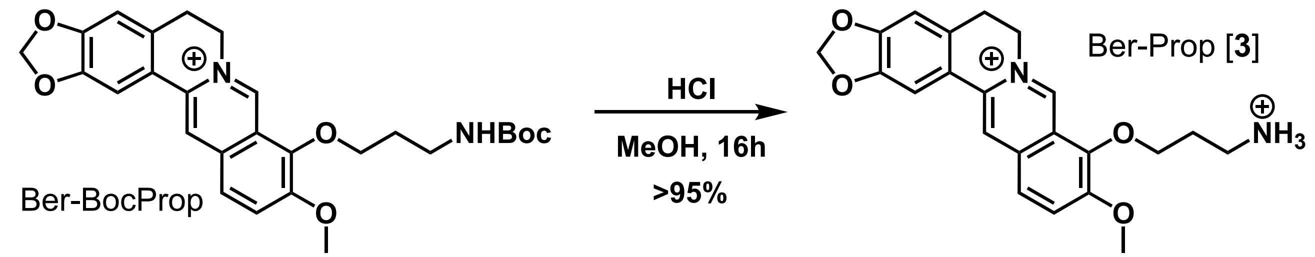


**Ber-Prop [3]: 9-(3-ammoniopropoxy)-10-methoxy-5,6-dihydro-[1,3]dioxolo[4,5-g]isoquinolino[3,2-a]isoquinolin-7-ium dichloride.** Carbamate **Ber-BocProp** (10 mg, 0.019 mmol, 1 eq.) was dissolved in MeOH (4 mL) and HCl (12M, 2.0 mL, 24 mmol) was added. Reaction was stirred at room temperature for 24 hours, then solvent was removed *in vacuo* to afford the pure HCl salt product as a yellow powder (8.5 mg, >95% yield). $^{1}$**H NMR** (600 MHz, DMSO-$d_6$) δ 9.98 (s, 1H), 8.94 (s, 1H), 8.24 – 8.17 (m, 4H), 7.98 (d, $J$ = 9.1 Hz, 1H), 7.79 (s, 1H), 7.08 (s, 1H), 6.16 (s, 2H), 5.01 (t, $J$ = 6.2 Hz, 2H), 4.37 (t, $J$ = 5.8 Hz, 2H), 4.07 (s, 3H), 3.22 – 3.17 (m, 2H), 3.10 (q, $J$ = 6.7, 6.3 Hz, 2H), 2.18 (p, $J$ = 6.3 Hz, 2H). $^{13}$**C NMR** (151 MHz, DMSO) δ 150.66, 150.32, 148.18, 145.96, 143.14, 137.97, 133.44, 131.21, 127.11, 123.97, 121.93, 120.93, 120.67, 108.93, 105.91, 102.58, 72.04, 57.60, 55.60, 36.64, 28.30, 26.80. **HRMS** Accurate mass (ES-): Found 379.16512 (-0.31 ppm), $C_{22}H_{23}N_2O_4$ ($M^{2+}$ -$H^+$) requires 379.16523.

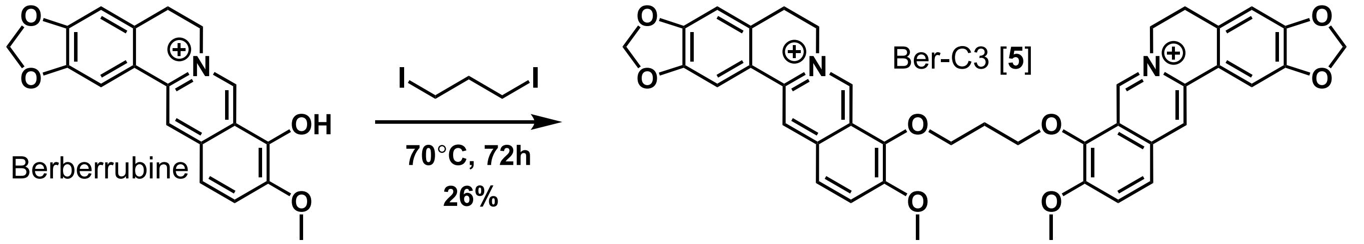


**Ber-C3 [5]: 9,9'-(propane-1,4-diylbis(oxy))bis(10-methoxy-5,6-dihydro-[1,3]dioxolo[4,5-g]isoquinolino[3,2-a]isoquinolin-7-ium) dichloride.** Following general procedure A, diiodide (16.6 mg. 0.056 mmol) yielded the title compound as a yellow solid (11.2 mg, 26% yield). $^{1}$**H NMR** (600 MHz, DMSO-$d_6$) δ 9.85 (s, 2H), 8.97 (s, 2H), 8.22 (d, $J$ = 9.2 Hz, 2H), 8.01 (d, $J$ = 9.1 Hz, 2H), 7.81 (s, 2H), 7.11 (s, 2H), 6.19 (s, 4H), 4.97 – 4.92 (m, 4H), 4.59 (t, $J$ = 6.4 Hz, 4H), 4.04 (s, 6H), 3.24 – 3.19 (m, 4H), 2.57 – 2.52 (m, 2H). $^{13}$**C NMR** (151 MHz, DMSO) δ 150.86, 150.36, 148.21, 145.77, 143.24, 138.02, 133.54, 131.17, 127.24, 123.98, 122.05, 120.90, 120.74, 108.92, 105.93, 102.61, 72.15, 57.60, 55.76, 30.92, 26.80. **HRMS** Accurate mass (ES+): Found 342.12269 (-1.01 ppm), $C_{41}H_{36}N_2O_8$ ($M^{2+}$) requires 342.12303.

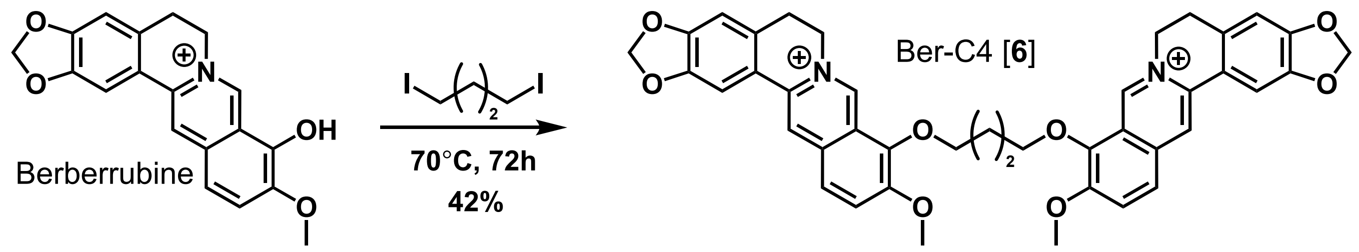


**Ber-C4 [6]: 9,9'-(butane-1,4-diylbis(oxy))bis(10-methoxy-5,6-dihydro-[1,3]dioxolo[4,5-g]isoquinolino[3,2-a]isoquinolin-7-ium) dichloride.** Following general procedure A, diiodide (17.4 mg. 0.056 mmol) yielded the title compound as a yellow solid (18.3 mg, 42% yield). **$^{1}$H NMR** (500 MHz, DMSO-$d_6$) δ 9.82 (s, 2H), 8.96 (s, 2H), 8.21 (d, *J* = 7.9 Hz, 2H), 8.00 (d, *J* = 7.9 Hz, 2H), 7.81 (s, 2H), 7.10 (s, 2H), 6.18 (s, 4H), 4.96 (s, 4H), 4.41 (s, 4H), 4.05 (s, 6H), 3.20 (s, 4H), 2.14 (s, 4H). **$^{13}$C NMR** (151 MHz, DMSO-$d_6$) δ 150.90, 150.32, 148.18, 145.76, 143.17, 137.97, 133.47, 131.16, 127.10, 123.92, 122.11, 120.91, 120.73, 108.92, 105.90, 102.61, 76.98, 74.41, 57.51, 55.78, 26.58. **HRMS** Accurate mass (ES+): Found 349.13082 (-0.11 ppm), $C_{42}H_{38}N_2O_8$ ($M^{2+}$) requires 349.13086.

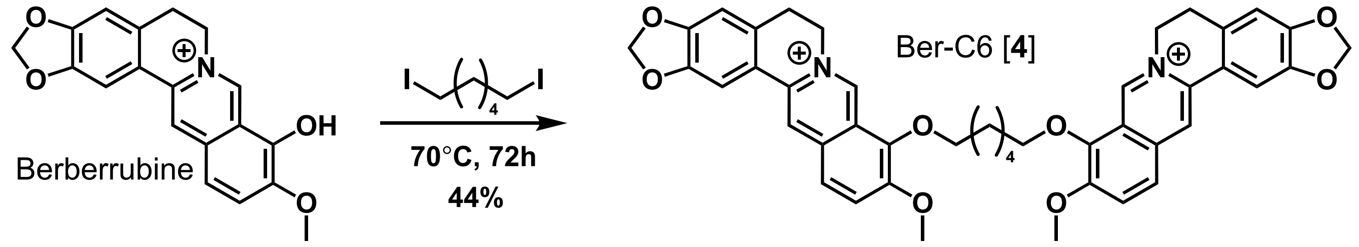


**Ber-C6 [4]: 9,9'-(hexane-1,4-diylbis(oxy))bis(10-methoxy-5,6-dihydro-[1,3]dioxolo[4,5-g]isoquinolino[3,2-a]isoquinolin-7-ium) dichloride.** Following general procedure A, diiodide (38.0 mg. 0.112 mmol) yielded the title compound as a yellow solid (39.3 mg, 44% yield). **$^{1}$H NMR** (600 MHz, DMSO-$d_6$) δ 9.75 (s, 2H), 8.93 (s, 2H), 8.20 (d, *J* = 9.2 Hz, 2H), 7.99 (d, *J* = 9.1 Hz, 2H), 7.78 (s, 2H), 7.09 (s, 2H), 6.17 (s, 4H), 4.94 (t, *J* = 6.1 Hz, 4H), 4.31 (t, *J* = 6.7 Hz, 4H), 4.04 (s, 6H), 3.19 (t, *J* = 6.1 Hz, 4H), 1.94 (p, *J* = 6.2 Hz, 4H), 1.60 (p, *J* = 4.0 Hz, 4H). **$^{13}$C NMR** (151 MHz, DMSO) δ 150.41, 149.84, 147.69, 145.20, 142.83, 137.47, 133.03, 130.64, 126.70, 123.34, 121.66, 120.41, 120.23, 108.40, 105.39, 102.11, 74.23, 57.03, 55.26, 29.48, 26.31, 25.13. **HRMS** Accurate mass (ES+): Found 363.14631 (-0.55 ppm), $C_{44}H_{42}N_2O_8$ ($M^{2+}$) requires 363.14651.

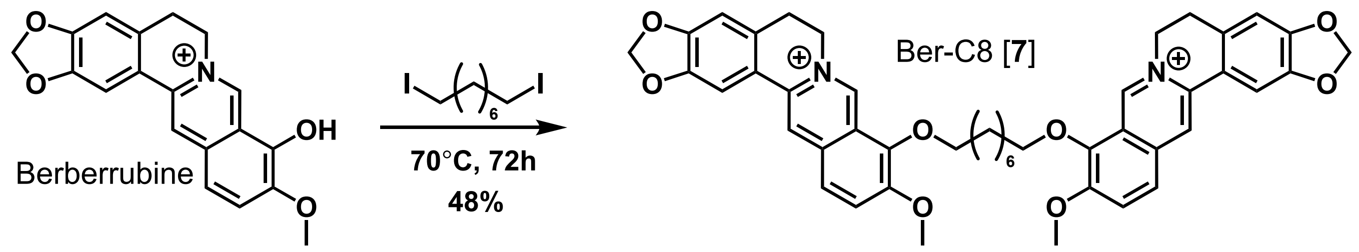


**Ber-C8 [7]: 9,9'-(octane-1,8-diylbis(oxy))bis(10-methoxy-5,6-dihydro-[1,3]dioxolo[4,5-g]isoquinolino[3,2-a]isoquinolin-7-ium) dichloride.** Following general procedure A, diiodide (20.5 mg. 0.056 mmol) yielded the title compound as a yellow solid (22.2 mg, 48% yield). $^{1}$**H NMR** (600 MHz, DMSO-$d_6$) δ 9.75 (s, 2H), 8.94 (s, 2H), 8.20 (d, *J* = 9.2 Hz, 2H), 8.00 (d, *J* = 9.1 Hz, 2H), 7.79 (s, 2H), 7.09 (s, 2H), 6.18 (s, 4H), 4.98 – 4.93 (m, 4H), 4.30 (t, *J* = 6.6 Hz, 4H), 4.04 (s, 6H), 3.23 – 3.18 (m, 4H), 1.91 (dt, *J* = 14.3, 6.9 Hz, 4H), 1.55 – 1.49 (m, 4H), 1.44 (s, 4H). $^{13}$**C NMR** (151 MHz, DMSO) δ 150.88, 150.30, 148.16, 145.71, 143.34, 137.94, 133.51, 131.13, 127.19, 123.79, 122.14, 120.91, 120.71, 108.88, 105.89, 102.58, 74.76, 57.54, 55.78, 29.99, 29.33, 26.80, 25.76. **HRMS** Accurate mass (ES+): Found 377.16182 (-0.90 ppm), $C_{46}H_{46}N_2O_8$ ($M^{2+}$) requires 377.16216.

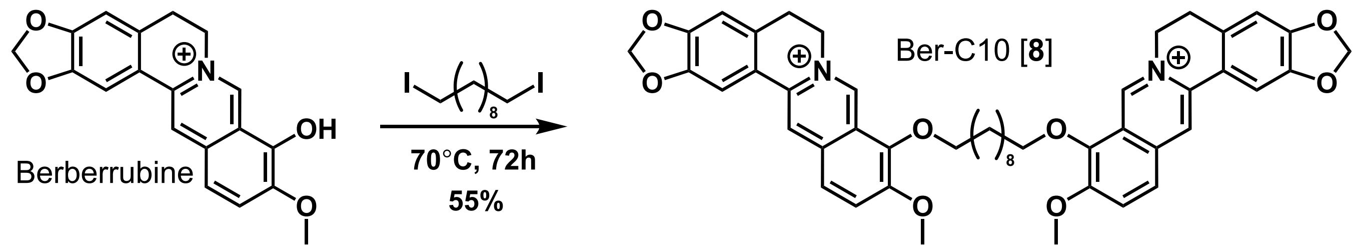


**Ber-C10 [8]: 9,9'-(decane-1,4-diylbis(oxy))bis(10-methoxy-5,6-dihydro-[1,3]dioxolo[4,5-g]isoquinolino[3,2-a]isoquinolin-7-ium) dichloride.** Following general procedure A, diiodide (23.0 mg. 0.058 mmol) yielded the title compound as a yellow solid (27.3 mg, 55% yield). $^{1}$**H NMR** (600 MHz, DMSO-$d_6$) δ 9.75 (s, 2H), 8.95 (s, 2H), 8.20 (d, *J* = 9.1 Hz, 2H), 8.00 (d, *J* = 9.0 Hz, 2H), 7.80 (s, 2H), 7.09 (s, 2H), 6.18 (s, 4H), 4.98 – 4.93 (m, 4H), 4.29 (t, *J* = 6.7 Hz, 4H), 4.05 (s, 6H), 3.21 (d, *J* = 11.3 Hz, 4H), 1.89 (dt, *J* = 14.4, 6.9 Hz, 4H), 1.51 – 1.46 (m, 4H), 1.41 – 1.22 (m, 22H). $^{13}$**C NMR** (151 MHz, DMSO) δ 150.89, 150.30, 148.17, 145.74, 143.35, 137.94, 133.51, 131.15, 127.18, 123.78, 122.14, 120.92, 120.70, 108.89, 105.90, 102.58, 74.74, 57.54, 55.79, 29.99, 29.54, 29.36, 26.81, 25.78. **HRMS** Accurate mass (ES+): Found 391.1773 (-1.31 ppm), $C_{48}H_{90}N_2O_8$ ($M^{2+}$) requires 391.17781.

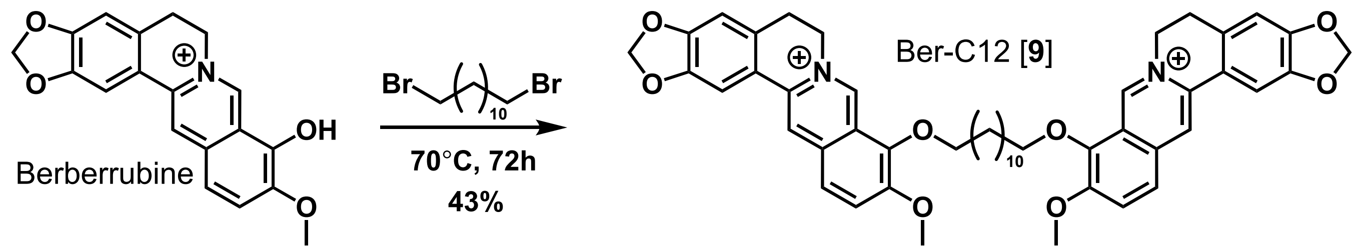


**Ber-C12 [9]: 9,9'-(dodecane-1,4-diylbis(oxy))bis(10-methoxy-5,6-dihydro-[1,3]dioxolo[4,5-g]isoquinolino[3,2-a]isoquinolin-7-ium) dichloride.** Following general procedure B, dibromide (15.3 mg. 0.047 mmol) yielded the title compound as a yellow solid (17.8 mg, 43% yield). **$^{1}$H NMR** (600 MHz, DMSO-*d*$_6$) δ 9.75 (s, 2H), 8.94 (s, 2H), 8.20 (d, *J* = 9.1 Hz, 2H), 7.99 (d, *J* = 9.0 Hz, 2H), 7.80 (s, 2H), 7.09 (s, 2H), 6.18 (s, 4H), 4.99 – 4.91 (m, 4H), 4.28 (t, *J* = 6.5 Hz, 4H), 4.05 (s, 6H), 3.23 – 3.18 (m, 4H), 1.93 – 1.82 (m, 4H), 1.52 – 1.43 (m, 4H), 1.40 – 1.21 (m, 22H). **$^{13}$C NMR** (151 MHz, DMSO) δ 150.89, 150.31, 148.18, 145.75, 143.35, 137.94, 133.51, 131.15, 127.19, 123.78, 122.15, 120.93, 120.72, 108.90, 105.90, 102.58, 74.74, 57.53, 55.79, 29.99, 29.57, 29.50, 29.37, 26.82, 25.77. **HRMS** Accurate mass (ES+): Found 405.19332 (-0.33 ppm), $C_{50}H_{54}N_2O_8$ ($M^{2+}$) requires 405.19346.

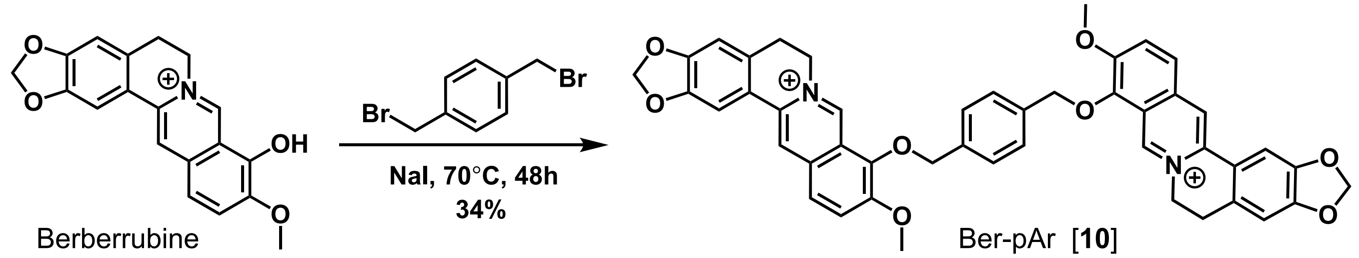


**Ber-pAr [10]: 9,9'-((1,4-phenylenebis(methylene))bis(oxy))bis(10-methoxy-5,6-dihydro-[1,3]dioxolo[4,5-g]isoquinolino[3,2-a]isoquinolin-7-ium) dichloride.** Following general procedure B, dibromide (29.2 mg. 0.111 mmol) yielded the title compound as a yellow solid (34.0 mg, 34% yield). **$^{1}$H NMR** (600 MHz, DMSO-*d*$_6$) δ 9.70 (s, 2H), 8.91 (s, 2H), 8.21 (d, *J* = 9.2 Hz, 2H), 8.00 (d, *J* = 9.0 Hz, 2H), 7.75 (s, 2H), 7.60 (s, 4H), 7.05 (s, 2H), 6.16 (s, 4H), 5.34 (s, 4H), 4.91 (t, *J* = 6.2 Hz, 4H), 4.07 (s, 6H), 3.22 – 3.14 (t, *J* = 6.2 Hz, 4H). **$^{13}$C NMR** (151 MHz, DMSO) δ 150.67, 149.82, 147.66, 145.16, 141.79, 137.38, 136.58, 132.87, 130.59, 128.85, 126.52, 123.79, 121.82, 120.33, 120.17, 108.35, 105.35, 102.11, 75.02, 57.03, 55.25, 26.30. **HRMS** Accurate mass (ES+): Found 373.13073 (-0.35 ppm), $C_{46}H_{38}N_2O_8$ ($M^{2+}$) requires 373.13086.

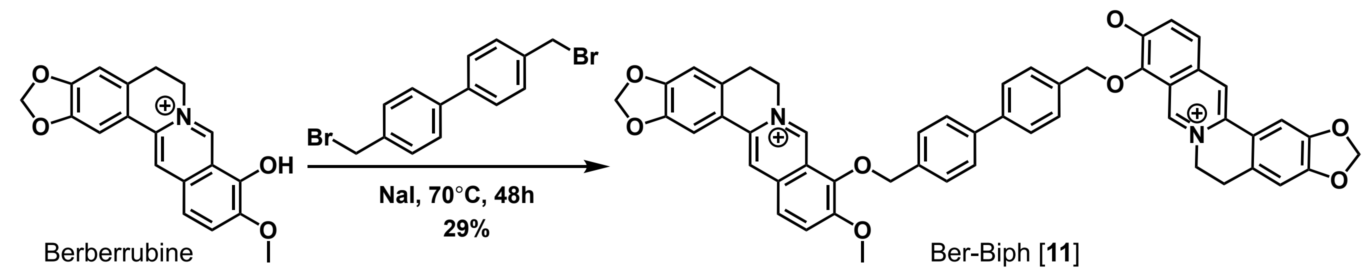


**9,9'-Ber-Biph [11]: (([1,1'-biphenyl]-4,4'-diylbis(methylene))bis(oxy))bis(10-methoxy-5,6-dihydro-[1,3]dioxolo[4,5-g]isoquinolino[3,2-a]isoquinolin-7-ium) dichloride.** Following general procedure B, dibromide (38.2 mg, 0.112 mmol) yielded the title compound as a yellow solid (28.5 mg, 29% yield). **$^{1}$H NMR** (600 MHz, DMSO-$d_6$) δ 9.80 (s, 2H), 8.95 (s, 2H), 8.24 (d, J = 9.2 Hz, 2H), 8.02 (d, J = 9.1 Hz, 2H), 7.80 (s, 2H), 7.74 (d, J = 8.3 Hz, 4H), 7.70 (d, J = 8.4 Hz, 4H), 7.10 (s, 2H), 6.18 (s, 4H), 5.41 (s, 4H), 4.94 (t, J = 6.4 Hz, 4H), 4.11 (s, 6H), 3.20 (t, J = 6.4, 4H). **$^{13}$C NMR** (151 MHz, DMSO-$d_6$) δ 150.69, 149.88, 147.72, 145.35, 142.06, 139.62, 137.48, 135.91, 132.97, 130.68, 129.65, 129.38, 126.63, 123.79, 121.80, 120.42, 120.28, 108.44, 105.44, 102.12, 75.05, 57.09, 55.34, 26.35. **HRMS** Accurate mass (ES+): Found 411.14652 (+0.02 ppm), $C_{52}H_{42}N_2O_8$ ($M^{2+}$) requires 411.14651.

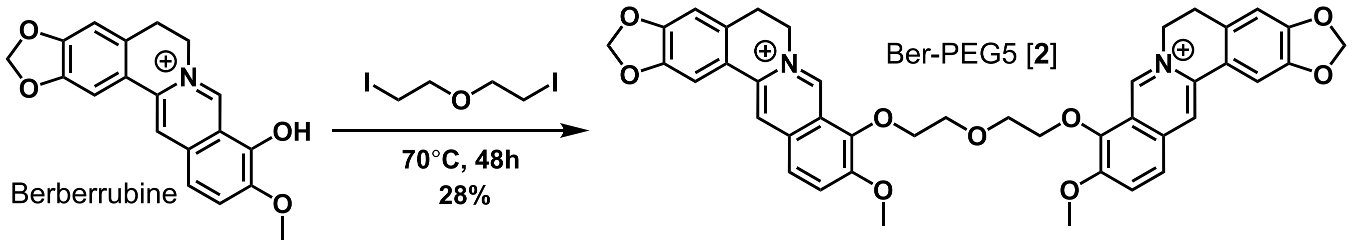


**Ber-PEG5 [12]: 9,9'-((oxybis(ethane-2,1-diyl))bis(oxy))bis(10-methoxy-5,6-dihydro-[1,3]dioxolo[4,5-g]isoquinolino[3,2-a]isoquinolin-7-ium) dichloride.** Following general procedure A, diiodide (15.0 mg. 0.046 mmol) yielded the title compound as a yellow solid (12.3 mg, 28% yield). **$^{1}$H NMR** (600 MHz, DMSO-$d_6$) δ 9.54 (s, 2H), 8.75 (s, 2H), 8.09 (d, *J* = 9.2 Hz, 2H), 7.93 – 7.85 (m, 2H), 7.39 (s, 2H), 7.04 (s, 2H), 6.13 (s, 4H), 4.88 (t, *J* = 5.7 Hz, 4H), 4.30 – 4.23 (m, 4H), 3.92 (s, 6H), 3.88 – 3.84 (m, 4H), 3.18 – 3.11 (m, 4H). **$^{13}$C NMR** (151 MHz, DMSO) δ 150.14, 149.70, 147.50, 145.08, 142.58, 137.04, 132.77, 130.19, 126.10, 123.56, 121.82, 120.11, 119.95, 108.22, 105.04, 102.14, 72.83, 69.48, 56.79, 55.06, 26.16. **HRMS** Accurate mass (ES+): Found 357.12815 (-0.47 ppm), $C_{42}H_{38}N_2O_9$ ($M^{2+}$) requires 357.12832.

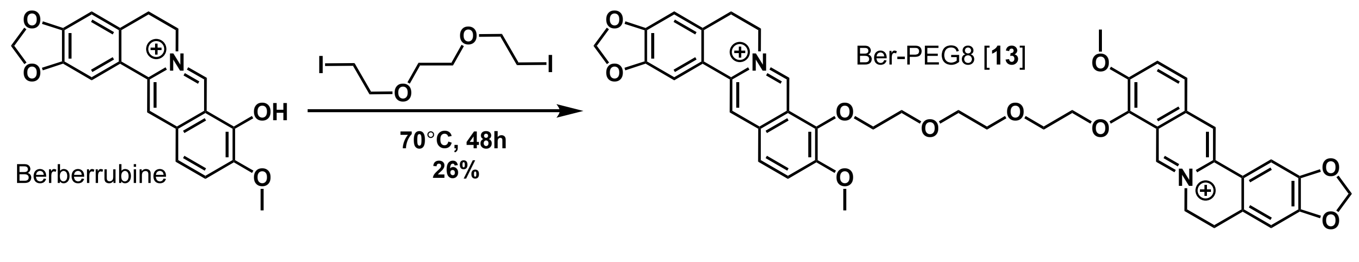


**Ber-PEG8 [13]: 9,9'-(((ethane-1,2-diylbis(oxy))bis(ethane-2,1-diyl))bis(oxy))bis(10-methoxy-5,6-dihydro-[1,3]dioxolo[4,5-g]isoquinolino[3,2-a]isoquinolin-7-ium) dichloride.** Following general procedure A, diiodide (17.0 mg. 0.047 mmol) yielded the title compound as a yellow solid (12.1 mg, 26% yield). **$^1$H NMR** (600 MHz, DMSO-$d_6$) δ 9.69 (s, 2H), 8.86 (s, 2H), 8.11 (d, $J$ = 9.2 Hz, 2H), 7.92 (d, $J$ = 9.1 Hz, 2H), 7.68 (s, 2H), 7.05 (s, 2H), 6.15 (s, 4H), 4.92 (t, $J$ = 6.1 Hz, 4H), 4.35 – 4.31 (m, 4H), 3.95 (s, 6H), 3.83 – 3.79 (m, 4H), 3.67 (s, 4H), 3.22 – 3.17 (m, 4H). **$^{13}$C NMR** (151 MHz, DMSO) δ 150.77, 150.24, 148.08, 145.70, 142.80, 137.79, 133.28, 130.96, 126.79, 123.98, 122.10, 120.79, 120.55, 108.78, 105.73, 102.60, 73.66, 70.13, 69.90, 57.34, 55.89, 26.78. **HRMS** Accurate mass (ES+): Found 379.1414 (-0.07 ppm), $C_{44}H_{42}N_2O_{10}$ ($M^{2+}$) requires 379.14142.


### Appendix

The following pages show $^1$H NMR (*top*) and $^{13}$C NMR (*bottom*) spectra for all synthesized intermediates and berberine analogs. All spectra were taken in DMSO-$d_6$ with the exception of berberrubine ($CDCl_3$) and Ber-Carb [**2**] ($D_2O$).

**Supplementary Methods: Details of chemical synthesis and compound analysis**

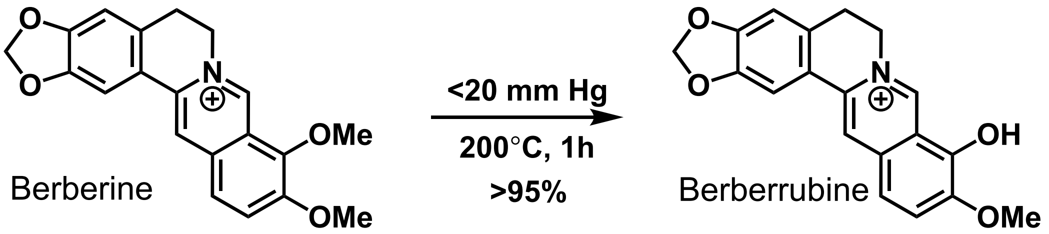


**Berberrubine chloride.**[2] Berberine chloride (500 mg, 1.34 mmol, 1 eq.) was heated to 200°C under vacuum (<20 mm Hg) for 45 minutes with stirring, causing a color change from yellow to red. Solid was filtered, washing with chloroform. Filtrate was purified by silica column chromatography (0-10% MeOH in DCM) to yield the title compound as a dark red solid (480 mg, >95% yield). **$^1$H NMR** (600 MHz, Chloroform-*d*) δ 9.16 (s, 1H), 7.54 (s, 1H), 7.21 (d, *J* = 7.9 Hz, 1H), 7.20 (s, 1H), 6.71 (s, 1H), 6.46 (d, *J* = 7.9 Hz, 1H), 6.02 (s, 2H), 4.36 (s, 2H), 3.87 (s, 3H), 3.08 – 3.02 (m, 2H). **$^{13}$C NMR** (151 MHz, $CDCl_3$) δ 167.92, 150.81, 148.99, 148.07, 145.61, 132.90, 131.27, 128.08, 122.16, 120.36, 120.01, 117.48, 108.32, 104.47, 102.84, 101.77, 56.03, 53.19, 28.57. **HRMS** Accurate mass (ES+): Found 322.10777 (+1.21 ppm), $C_{19}H_{16}NO_4$ ($M^+$) requires 322.10738.

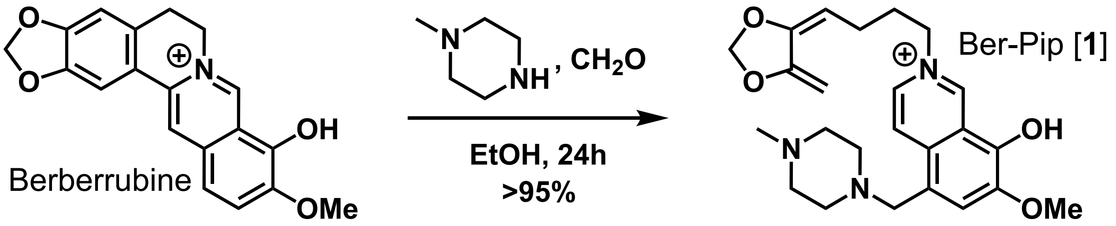


**Ber-Pip [1]: 9-hydroxy-10-methoxy-12-((4-methylpiperazin-1-yl)methyl)-5,6-dihydro-[1,3]dioxolo[4,5-g]isoquinolino[3,2-a]isoquinolin-7-ium chloride.**[3] To a solution of berberrubine (24 mg, 0.067 mmol, 1 eq.) in anhydrous EtOH (1 mL) was added N-methylpiperazine (0.037 mL, 0.335 mmol, 5 eq.) and formaldehyde (37% aq., 0.035 mL, 0.335 mmol, 5 eq.). The solution stirred at room temperature for 24 hours and was then concentrated *in vacuo* and purified by silica column chromatography (0-5% MeOH/DCM) to afford the title compound as a yellow powder (30.5 mg, >95% yield). **$^1$H NMR** (600 MHz, $CDCl_3$) δ 9.21 (s, 1H), 8.03 (s, 1H), 7.21 (s, 1H), 7.18 (s, 1H), 6.70 (s, 1H), 6.02 (s, 2H), 4.38 (t, *J* = 5.9 Hz, 2H), 3.86 (s, 3H), 3.65 (s, 2H), 3.04 (t, *J* = 6.0 Hz, 2H), 2.52 (s, 8H), 2.29 (s, 3H). **$^{13}$C NMR** (151 MHz, $CDCl_3$) δ 167.15, 149.90, 149.15, 148.21, 145.75, 132.96, 129.71, 128.33, 123.82, 122.55, 120.47, 115.07, 110.00, 108.45, 104.71, 101.87, 64.39, 59.75, 56.33, 55.19, 53.30, 52.68, 45.81, 28.64, 25.49. **HRMS** Accurate mass (ES+): Found 434.20693 (-1.17 ppm), $C_{25}H_{28}N_3O_4$ ($M^+$) requires 434.20743.

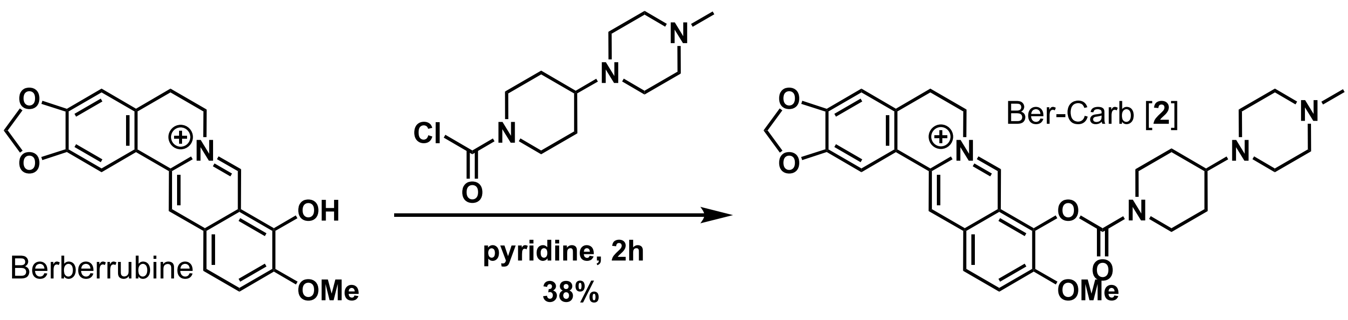


**Ber-Carb [2]: 10-methoxy-9-((4-(4-methylpiperazin-1-yl)piperidine-1-carbonyl)oxy)-5,6-dihydro-[1,3]dioxolo[4,5-g]isoquinolino[3,2-a]isoquinolin-7-ium chloride formate.**[4] Berberrubine (96.4 mg, 0.300 mmol, 1 eq.) was dissolved in pyridine (1 mL) and added in one portion to 4-(4-methylpiperazin-1-yl)piperidine-1-carbonyl chloride (369 mg, 1.50 mmol, 5 eq.)**.** The solution was stirred at room temperature for 2 hours, then was concentrated *in vacuo* and purified by silica column chromatography (0-5% MeOH/DCM) to yield the title compound as a yellow solid (64.6 mg, 38% yield) **$^1$H NMR** (600 MHz, $D_2O$) δ 9.24 (s, 1H), 8.41 (s, 1H), 8.15 (s, 1H), 7.84 (d, *J* = 19.0 Hz, 2H), 7.14 (s, 1H), 6.80 (s, 1H), 6.00 (s, 2H), 4.65 (s, 2H), 4.40 (s, 1H), 4.11 (d, *J* = 12.9 Hz, 1H), 3.92 (s, 3H), 3.68 – 3.52 (m, 2H), 3.22 – 3.16 (m, 1H), 3.09 – 2.97 (m, 4H), 2.62 (br s, 4H), 2.27 (s, 4H), 2.05 (br s, 2H), 1.56 (d, *J* = 8.5 Hz, 1H), 1.47 (d, *J* = 10.4 Hz, 1H). **$^{13}$C NMR** (151 MHz, $D_2O$) δ 171.02, 153.34, 150.46, 150.15, 147.64, 142.69, 138.04, 133.70, 132.89, 130.14, 126.43, 125.25, 121.36, 119.60, 108.28, 104.98, 102.28, 60.58, 56.89, 55.97, 53.54, 47.84, 27.57, 27.23, 26.40. **HRMS** Accurate mass (ES+): Found 531.26012 (-0.14 ppm), $C_{30}H_{35}N_4O_5$ ($M^+$) requires 531.26020.

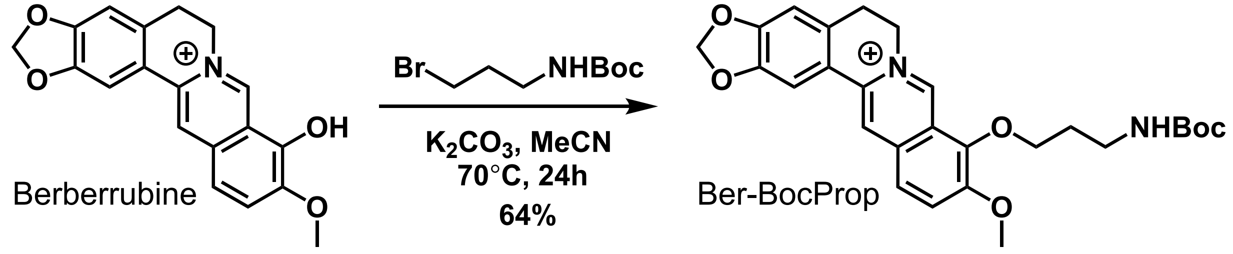


**9-(3-((tert-butoxycarbonyl)amino)propoxy)-10-methoxy-5,6-dihydro-[1,3]dioxolo[4,5-g]isoquinolino[3,2-a]isoquinolin-7-ium chloride.** To a solution of berberrubine (50 mg, 0.134 mmol, 1 eq.) and $K_2CO_3$ (18.6 mg, 0.403 mmol, 3 eq.) in acetonitrile (1 mL) preheated to 70°C was added tert-butyl (3-bromopropyl)carbamate (64 mg, 0.269 mmol, 2 eq.). Additional acetonitrile (0.5 mL) was added, the vessel was sealed, and the reaction continued to stir at 70°C for 24 hours. The crude reaction mixture was cooled to RT and purified by silica column chromatography (0-10% MeOH/DCM), yielding the title compound as a yellow powder (45.9 mg, 64%) **$^1$H NMR** (500 MHz, DMSO-$d_6$) δ 9.83 (s, 1H), 8.93 (s, 1H), 8.19 (d, *J* = 9.2 Hz, 1H), 7.97 (d, *J* = 9.0 Hz, 1H), 7.79 (s, 1H), 7.09 (s, 1H), 6.97 (t, *J* = 5.7 Hz, 1H), 6.16 (s, 2H), 5.00 – 4.89 (m, 2H), 4.28 (t, *J* = 6.0 Hz, 2H), 4.04 (s, 3H), 3.24 – 3.15 (m, 4H), 1.94 (p, *J* = 5.9, 5.5 Hz, 2H), 1.34 (s, 9H). **$^{13}$C NMR** (151 MHz, DMSO-d$_6$) δ 155.87, 150.28, 149.80, 147.67, 145.43, 142.73, 137.37, 132.98, 130.60, 126.63, 123.34, 121.53, 120.43, 120.16, 108.43, 105.43, 102.10, 77.65, 71.80, 57.06, 55.30, 54.95, 36.68, 30.28, 28.25, 26.34. **HRMS** Accurate mass (ES+): Found 479.21776 (-0.14 ppm), $C_{27}H_{31}N_2O_6$ ($M^+$) requires 479.21766.

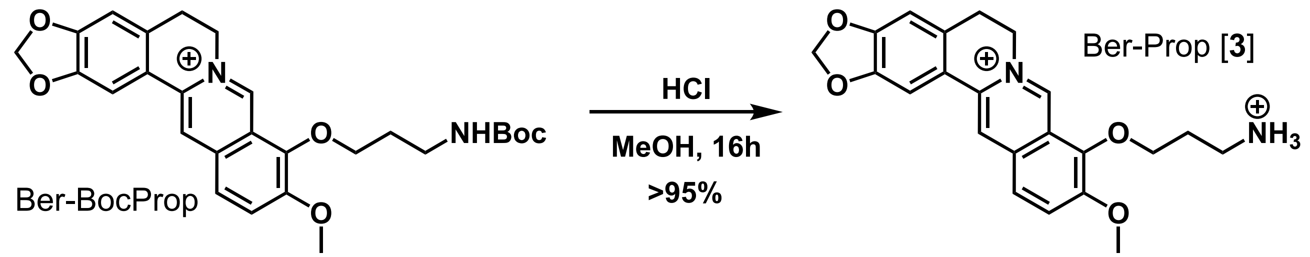


**Ber-Prop [3]: 9-(3-ammoniopropoxy)-10-methoxy-5,6-dihydro-[1,3]dioxolo[4,5-g]isoquinolino[3,2-a]isoquinolin-7-ium dichloride.** Carbamate **Ber-BocProp** (10 mg, 0.019 mmol, 1 eq.) was dissolved in MeOH (4 mL) and HCl (12M, 2.0 mL, 24 mmol) was added. Reaction was stirred at room temperature for 24 hours, then solvent was removed *in vacuo* to afford the pure HCl salt product as a yellow powder (8.5 mg, >95% yield). $^{1}$**H NMR** (600 MHz, DMSO-$d_6$) δ 9.98 (s, 1H), 8.94 (s, 1H), 8.24 – 8.17 (m, 4H), 7.98 (d, $J$ = 9.1 Hz, 1H), 7.79 (s, 1H), 7.08 (s, 1H), 6.16 (s, 2H), 5.01 (t, $J$ = 6.2 Hz, 2H), 4.37 (t, $J$ = 5.8 Hz, 2H), 4.07 (s, 3H), 3.22 – 3.17 (m, 2H), 3.10 (q, $J$ = 6.7, 6.3 Hz, 2H), 2.18 (p, $J$ = 6.3 Hz, 2H). $^{13}$**C NMR** (151 MHz, DMSO) δ 150.66, 150.32, 148.18, 145.96, 143.14, 137.97, 133.44, 131.21, 127.11, 123.97, 121.93, 120.93, 120.67, 108.93, 105.91, 102.58, 72.04, 57.60, 55.60, 36.64, 28.30, 26.80. **HRMS** Accurate mass (ES-): Found 379.16512 (-0.31 ppm), $C_{22}H_{23}N_2O_4$ ($M^{2+}$ -$H^+$) requires 379.16523.

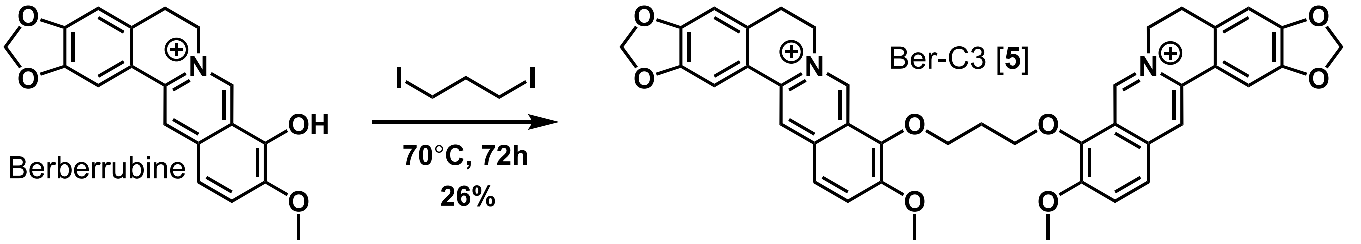


**Ber-C3 [5]: 9,9'-(propane-1,4-diylbis(oxy))bis(10-methoxy-5,6-dihydro-[1,3]dioxolo[4,5-g]isoquinolino[3,2-a]isoquinolin-7-ium) dichloride.** Following general procedure A, diiodide (16.6 mg. 0.056 mmol) yielded the title compound as a yellow solid (11.2 mg, 26% yield). $^{1}$**H NMR** (600 MHz, DMSO-$d_6$) δ 9.85 (s, 2H), 8.97 (s, 2H), 8.22 (d, $J$ = 9.2 Hz, 2H), 8.01 (d, $J$ = 9.1 Hz, 2H), 7.81 (s, 2H), 7.11 (s, 2H), 6.19 (s, 4H), 4.97 – 4.92 (m, 4H), 4.59 (t, $J$ = 6.4 Hz, 4H), 4.04 (s, 6H), 3.24 – 3.19 (m, 4H), 2.57 – 2.52 (m, 2H). $^{13}$**C NMR** (151 MHz, DMSO) δ 150.86, 150.36, 148.21, 145.77, 143.24, 138.02, 133.54, 131.17, 127.24, 123.98, 122.05, 120.90, 120.74, 108.92, 105.93, 102.61, 72.15, 57.60, 55.76, 30.92, 26.80. **HRMS** Accurate mass (ES+): Found 342.12269 (-1.01 ppm), $C_{41}H_{36}N_2O_8$ ($M^{2+}$) requires 342.12303.

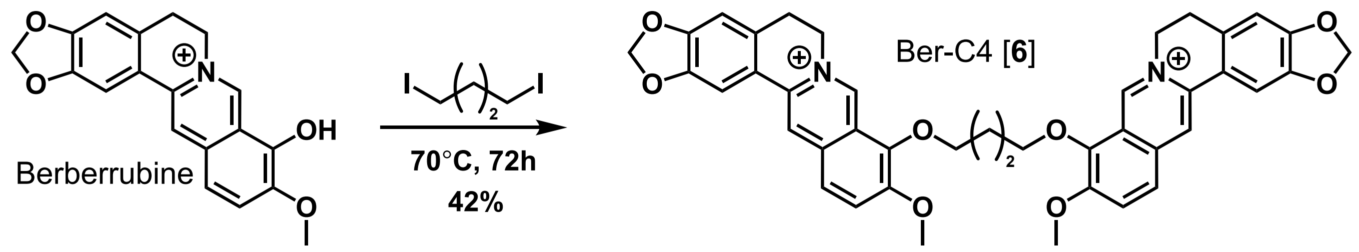


**Ber-C4 [6]: 9,9'-(butane-1,4-diylbis(oxy))bis(10-methoxy-5,6-dihydro-[1,3]dioxolo[4,5-g]isoquinolino[3,2-a]isoquinolin-7-ium) dichloride.** Following general procedure A, diiodide (17.4 mg. 0.056 mmol) yielded the title compound as a yellow solid (18.3 mg, 42% yield). **$^{1}$H NMR** (500 MHz, DMSO-$d_6$) δ 9.82 (s, 2H), 8.96 (s, 2H), 8.21 (d, *J* = 7.9 Hz, 2H), 8.00 (d, *J* = 7.9 Hz, 2H), 7.81 (s, 2H), 7.10 (s, 2H), 6.18 (s, 4H), 4.96 (s, 4H), 4.41 (s, 4H), 4.05 (s, 6H), 3.20 (s, 4H), 2.14 (s, 4H). **$^{13}$C NMR** (151 MHz, DMSO-$d_6$) δ 150.90, 150.32, 148.18, 145.76, 143.17, 137.97, 133.47, 131.16, 127.10, 123.92, 122.11, 120.91, 120.73, 108.92, 105.90, 102.61, 76.98, 74.41, 57.51, 55.78, 26.58. **HRMS** Accurate mass (ES+): Found 349.13082 (-0.11 ppm), $C_{42}H_{38}N_2O_8$ ($M^{2+}$) requires 349.13086.

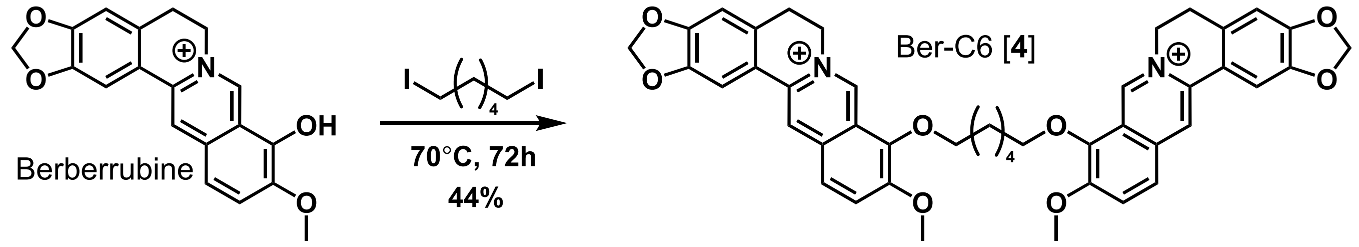


**Ber-C6 [4]: 9,9'-(hexane-1,4-diylbis(oxy))bis(10-methoxy-5,6-dihydro-[1,3]dioxolo[4,5-g]isoquinolino[3,2-a]isoquinolin-7-ium) dichloride.** Following general procedure A, diiodide (38.0 mg. 0.112 mmol) yielded the title compound as a yellow solid (39.3 mg, 44% yield). **$^{1}$H NMR** (600 MHz, DMSO-$d_6$) δ 9.75 (s, 2H), 8.93 (s, 2H), 8.20 (d, *J* = 9.2 Hz, 2H), 7.99 (d, *J* = 9.1 Hz, 2H), 7.78 (s, 2H), 7.09 (s, 2H), 6.17 (s, 4H), 4.94 (t, *J* = 6.1 Hz, 4H), 4.31 (t, *J* = 6.7 Hz, 4H), 4.04 (s, 6H), 3.19 (t, *J* = 6.1 Hz, 4H), 1.94 (p, *J* = 6.2 Hz, 4H), 1.60 (p, *J* = 4.0 Hz, 4H). **$^{13}$C NMR** (151 MHz, DMSO) δ 150.41, 149.84, 147.69, 145.20, 142.83, 137.47, 133.03, 130.64, 126.70, 123.34, 121.66, 120.41, 120.23, 108.40, 105.39, 102.11, 74.23, 57.03, 55.26, 29.48, 26.31, 25.13. **HRMS** Accurate mass (ES+): Found 363.14631 (-0.55 ppm), $C_{44}H_{42}N_2O_8$ ($M^{2+}$) requires 363.14651.

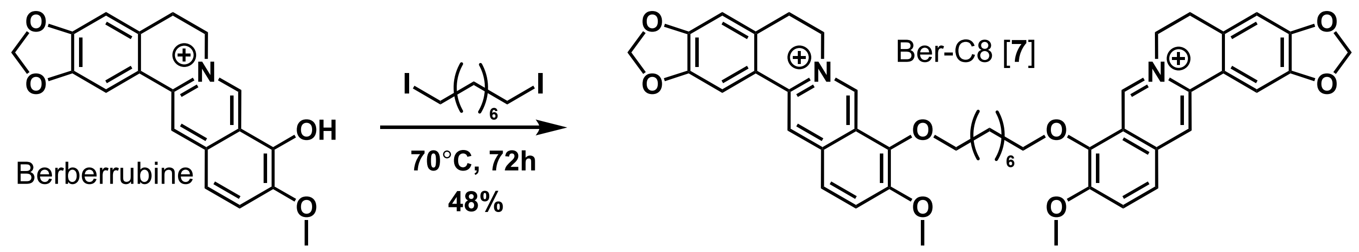


**Ber-C8 [7]: 9,9'-(octane-1,8-diylbis(oxy))bis(10-methoxy-5,6-dihydro-[1,3]dioxolo[4,5-g]isoquinolino[3,2-a]isoquinolin-7-ium) dichloride.** Following general procedure A, diiodide (20.5 mg. 0.056 mmol) yielded the title compound as a yellow solid (22.2 mg, 48% yield). **$^1$H NMR** (600 MHz, DMSO-$d_6$) δ 9.75 (s, 2H), 8.94 (s, 2H), 8.20 (d, $J$ = 9.2 Hz, 2H), 8.00 (d, $J$ = 9.1 Hz, 2H), 7.79 (s, 2H), 7.09 (s, 2H), 6.18 (s, 4H), 4.98 – 4.93 (m, 4H), 4.30 (t, $J$ = 6.6 Hz, 4H), 4.04 (s, 6H), 3.23 – 3.18 (m, 4H), 1.91 (dt, $J$ = 14.3, 6.9 Hz, 4H), 1.55 – 1.49 (m, 4H), 1.44 (s, 4H). **$^{13}$C NMR** (151 MHz, DMSO) δ 150.88, 150.30, 148.16, 145.71, 143.34, 137.94, 133.51, 131.13, 127.19, 123.79, 122.14, 120.91, 120.71, 108.88, 105.89, 102.58, 74.76, 57.54, 55.78, 29.99, 29.33, 26.80, 25.76. **HRMS** Accurate mass (ES+): Found 377.16182 (-0.90 ppm), $C_{46}H_{46}N_2O_8$ ($M^{2+}$) requires 377.16216.

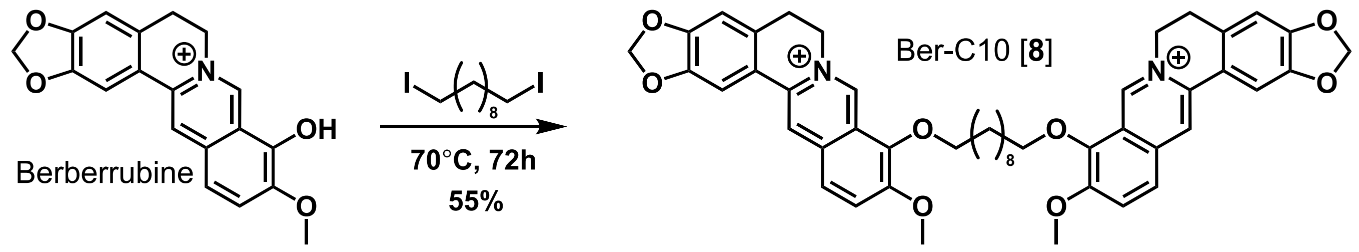


**Ber-C10 [8]: 9,9'-(decane-1,4-diylbis(oxy))bis(10-methoxy-5,6-dihydro-[1,3]dioxolo[4,5-g]isoquinolino[3,2-a]isoquinolin-7-ium) dichloride.** Following general procedure A, diiodide (23.0 mg. 0.058 mmol) yielded the title compound as a yellow solid (27.3 mg, 55% yield). **$^1$H NMR** (600 MHz, DMSO-$d_6$) δ 9.75 (s, 2H), 8.95 (s, 2H), 8.20 (d, $J$ = 9.1 Hz, 2H), 8.00 (d, $J$ = 9.0 Hz, 2H), 7.80 (s, 2H), 7.09 (s, 2H), 6.18 (s, 4H), 4.98 – 4.93 (m, 4H), 4.29 (t, $J$ = 6.7 Hz, 4H), 4.05 (s, 6H), 3.21 (d, $J$ = 11.3 Hz, 4H), 1.89 (dt, $J$ = 14.4, 6.9 Hz, 4H), 1.51 – 1.46 (m, 4H), 1.41 – 1.22 (m, 22H). **$^{13}$C NMR** (151 MHz, DMSO) δ 150.89, 150.30, 148.17, 145.74, 143.35, 137.94, 133.51, 131.15, 127.18, 123.78, 122.14, 120.92, 120.70, 108.89, 105.90, 102.58, 74.74, 57.54, 55.79, 29.99, 29.54, 29.36, 26.81, 25.78. **HRMS** Accurate mass (ES+): Found 391.1773 (-1.31 ppm), $C_{48}H_{90}N_2O_8$ ($M^{2+}$) requires 391.17781.

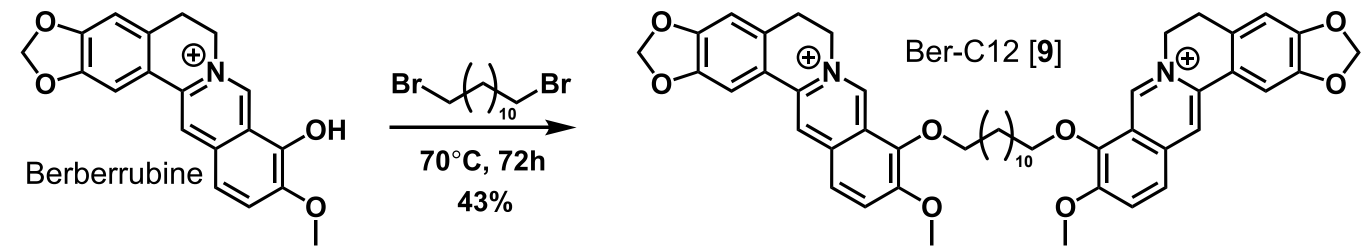


**Ber-C12 [9]: 9,9'-(dodecane-1,4-diylbis(oxy))bis(10-methoxy-5,6-dihydro-[1,3]dioxolo[4,5-g]isoquinolino[3,2-a]isoquinolin-7-ium) dichloride.** Following general procedure B, dibromide (15.3 mg. 0.047 mmol) yielded the title compound as a yellow solid (17.8 mg, 43% yield). $^{1}$**H NMR** (600 MHz, DMSO-$d_6$) δ 9.75 (s, 2H), 8.94 (s, 2H), 8.20 (d, *J* = 9.1 Hz, 2H), 7.99 (d, *J* = 9.0 Hz, 2H), 7.80 (s, 2H), 7.09 (s, 2H), 6.18 (s, 4H), 4.99 – 4.91 (m, 4H), 4.28 (t, *J* = 6.5 Hz, 4H), 4.05 (s, 6H), 3.23 – 3.18 (m, 4H), 1.93 – 1.82 (m, 4H), 1.52 – 1.43 (m, 4H), 1.40 – 1.21 (m, 22H). $^{13}$**C NMR** (151 MHz, DMSO) δ 150.89, 150.31, 148.18, 145.75, 143.35, 137.94, 133.51, 131.15, 127.19, 123.78, 122.15, 120.93, 120.72, 108.90, 105.90, 102.58, 74.74, 57.53, 55.79, 29.99, 29.57, 29.50, 29.37, 26.82, 25.77. **HRMS** Accurate mass (ES+): Found 405.19332 (-0.33 ppm), $C_{50}H_{54}N_2O_8$ ($M^{2+}$) requires 405.19346.

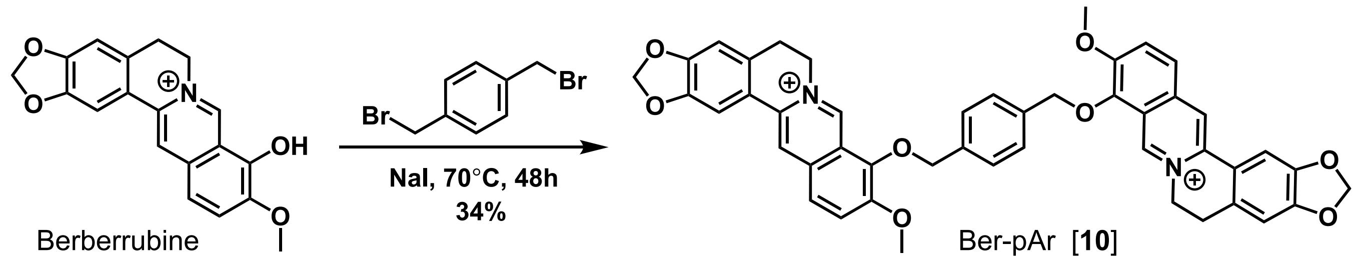


**Ber-pAr [10]: 9,9'-((1,4-phenylenebis(methylene))bis(oxy))bis(10-methoxy-5,6-dihydro-[1,3]dioxolo[4,5-g]isoquinolino[3,2-a]isoquinolin-7-ium) dichloride.** Following general procedure B, dibromide (29.2 mg. 0.111 mmol) yielded the title compound as a yellow solid (34.0 mg, 34% yield). $^{1}$**H NMR** (600 MHz, DMSO-$d_6$) δ 9.70 (s, 2H), 8.91 (s, 2H), 8.21 (d, *J* = 9.2 Hz, 2H), 8.00 (d, *J* = 9.0 Hz, 2H), 7.75 (s, 2H), 7.60 (s, 4H), 7.05 (s, 2H), 6.16 (s, 4H), 5.34 (s, 4H), 4.91 (t, *J* = 6.2 Hz, 4H), 4.07 (s, 6H), 3.22 – 3.14 (t, *J* = 6.2 Hz, 4H). $^{13}$**C NMR** (151 MHz, DMSO) δ 150.67, 149.82, 147.66, 145.16, 141.79, 137.38, 136.58, 132.87, 130.59, 128.85, 126.52, 123.79, 121.82, 120.33, 120.17, 108.35, 105.35, 102.11, 75.02, 57.03, 55.25, 26.30. **HRMS** Accurate mass (ES+): Found 373.13073 (-0.35 ppm), $C_{46}H_{38}N_2O_8$ ($M^{2+}$) requires 373.13086.

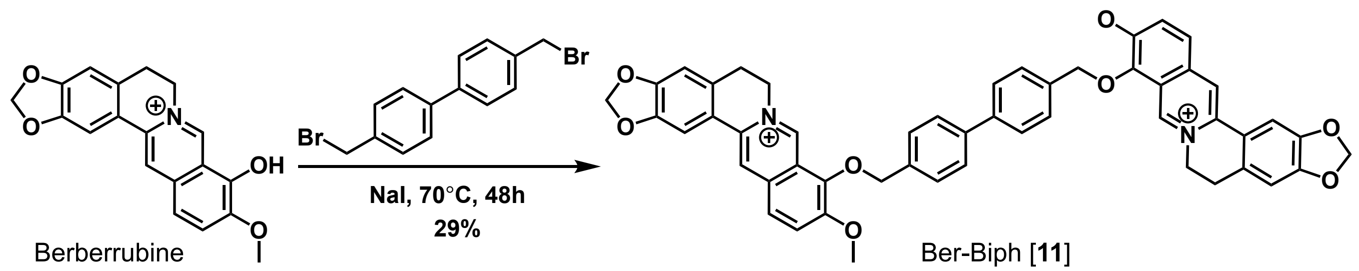


**9,9'-Ber-Biph [11]: (([1,1'-biphenyl]-4,4'-diylbis(methylene))bis(oxy))bis(10-methoxy-5,6-dihydro-[1,3]dioxolo[4,5-g]isoquinolino[3,2-a]isoquinolin-7-ium) dichloride.** Following general procedure B, dibromide (38.2 mg, 0.112 mmol) yielded the title compound as a yellow solid (28.5 mg, 29% yield). **$^{1}$H NMR** (600 MHz, DMSO-$d_6$) δ 9.80 (s, 2H), 8.95 (s, 2H), 8.24 (d, J = 9.2 Hz, 2H), 8.02 (d, J = 9.1 Hz, 2H), 7.80 (s, 2H), 7.74 (d, J = 8.3 Hz, 4H), 7.70 (d, J = 8.4 Hz, 4H), 7.10 (s, 2H), 6.18 (s, 4H), 5.41 (s, 4H), 4.94 (t, J = 6.4 Hz, 4H), 4.11 (s, 6H), 3.20 (t, J = 6.4, 4H). **$^{13}$C NMR** (151 MHz, DMSO-$d_6$) δ 150.69, 149.88, 147.72, 145.35, 142.06, 139.62, 137.48, 135.91, 132.97, 130.68, 129.65, 129.38, 126.63, 123.79, 121.80, 120.42, 120.28, 108.44, 105.44, 102.12, 75.05, 57.09, 55.34, 26.35. **HRMS** Accurate mass (ES+): Found 411.14652 (+0.02 ppm), $C_{52}H_{42}N_2O_8$ ($M^{2+}$) requires 411.14651.

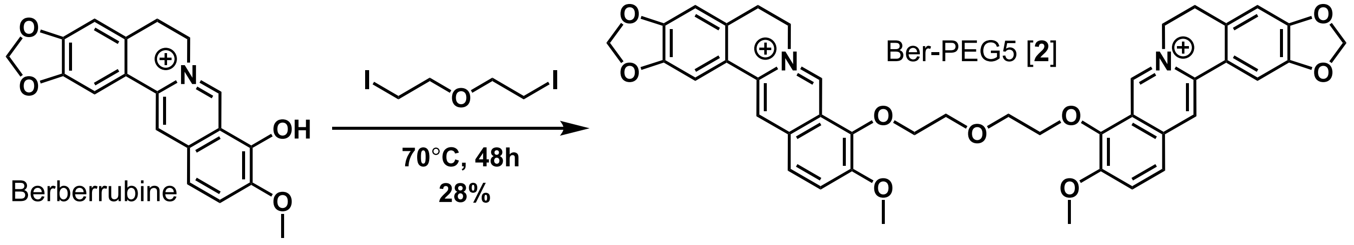


**Ber-PEG5 [12]: 9,9'-((oxybis(ethane-2,1-diyl))bis(oxy))bis(10-methoxy-5,6-dihydro-[1,3]dioxolo[4,5-g]isoquinolino[3,2-a]isoquinolin-7-ium) dichloride.** Following general procedure A, diiodide (15.0 mg. 0.046 mmol) yielded the title compound as a yellow solid (12.3 mg, 28% yield). **$^{1}$H NMR** (600 MHz, DMSO-$d_6$) δ 9.54 (s, 2H), 8.75 (s, 2H), 8.09 (d, *J* = 9.2 Hz, 2H), 7.93 – 7.85 (m, 2H), 7.39 (s, 2H), 7.04 (s, 2H), 6.13 (s, 4H), 4.88 (t, *J* = 5.7 Hz, 4H), 4.30 – 4.23 (m, 4H), 3.92 (s, 6H), 3.88 – 3.84 (m, 4H), 3.18 – 3.11 (m, 4H). **$^{13}$C NMR** (151 MHz, DMSO) δ 150.14, 149.70, 147.50, 145.08, 142.58, 137.04, 132.77, 130.19, 126.10, 123.56, 121.82, 120.11, 119.95, 108.22, 105.04, 102.14, 72.83, 69.48, 56.79, 55.06, 26.16. **HRMS** Accurate mass (ES+): Found 357.12815 (-0.47 ppm), $C_{42}H_{38}N_2O_9$ ($M^{2+}$) requires 357.12832.

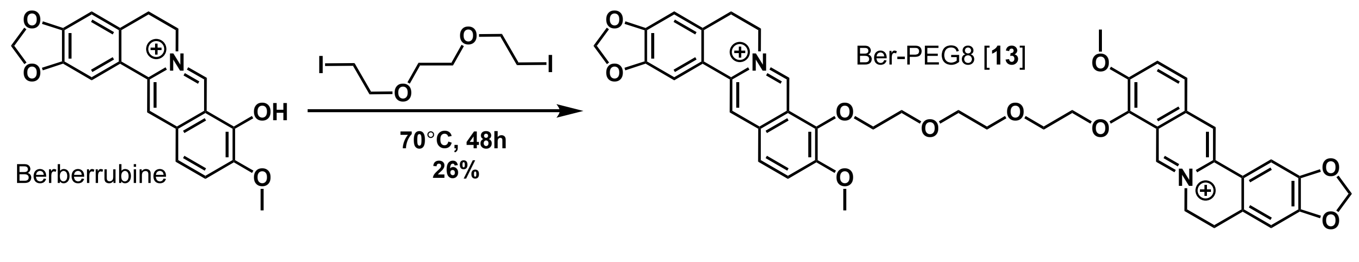


**Ber-PEG8 [13]: 9,9'-(((ethane-1,2-diylbis(oxy))bis(ethane-2,1-diyl))bis(oxy))bis(10-methoxy-5,6-dihydro-[1,3]dioxolo[4,5-g]isoquinolino[3,2-a]isoquinolin-7-ium) dichloride.** Following general procedure A, diiodide (17.0 mg. 0.047 mmol) yielded the title compound as a yellow solid (12.1 mg, 26% yield). **$^{1}$H NMR** (600 MHz, DMSO-$d_6$) δ 9.69 (s, 2H), 8.86 (s, 2H), 8.11 (d, $J$ = 9.2 Hz, 2H), 7.92 (d, $J$ = 9.1 Hz, 2H), 7.68 (s, 2H), 7.05 (s, 2H), 6.15 (s, 4H), 4.92 (t, $J$ = 6.1 Hz, 4H), 4.35 – 4.31 (m, 4H), 3.95 (s, 6H), 3.83 – 3.79 (m, 4H), 3.67 (s, 4H), 3.22 – 3.17 (m, 4H). **$^{13}$C NMR** (151 MHz, DMSO) δ 150.77, 150.24, 148.08, 145.70, 142.80, 137.79, 133.28, 130.96, 126.79, 123.98, 122.10, 120.79, 120.55, 108.78, 105.73, 102.60, 73.66, 70.13, 69.90, 57.34, 55.89, 26.78. **HRMS** Accurate mass (ES+): Found 379.1414 (-0.07 ppm), $C_{44}H_{42}N_2O_{10}$ ($M^{2+}$) requires 379.14142.


### Appendix

The following pages show $^{1}$H NMR (*top*) and $^{13}$C NMR (*bottom*) spectra for all synthesized intermediates and berberine analogs. All spectra were taken in DMSO-$d_6$ with the exception of berberrubine ($CDCl_3$) and Ber-Carb [**2**] ($D_2O$).

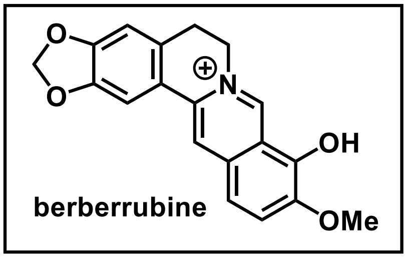


berberrubine

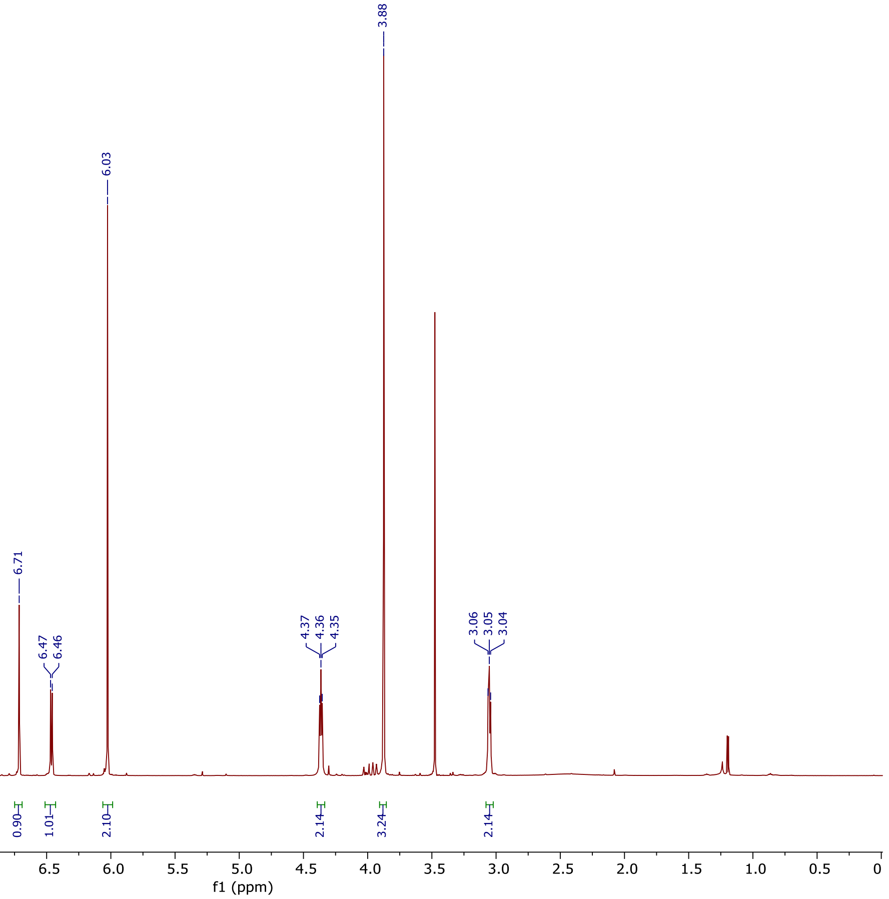


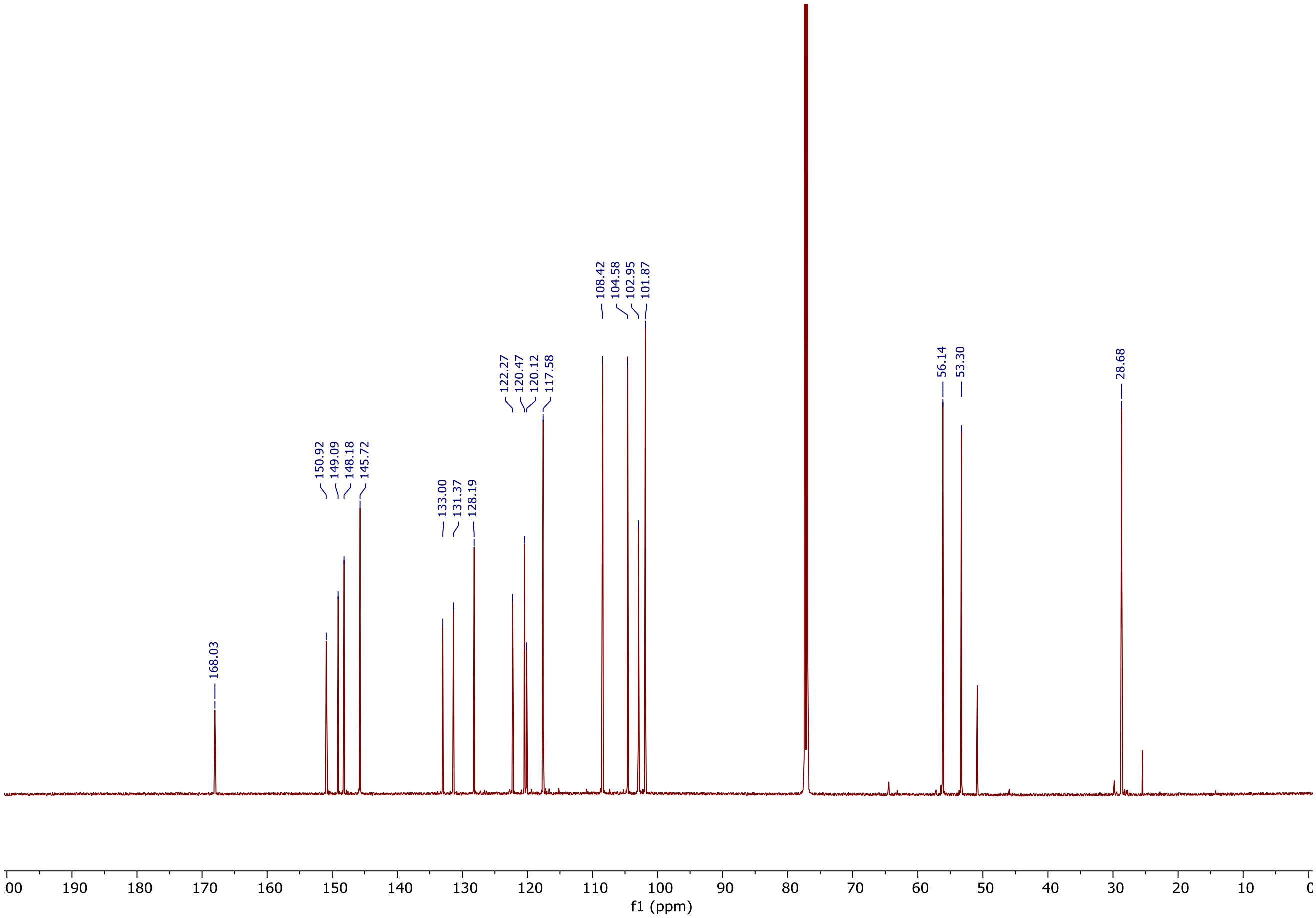

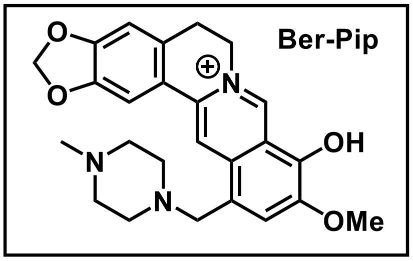
Ber-Pip
OH
OMe

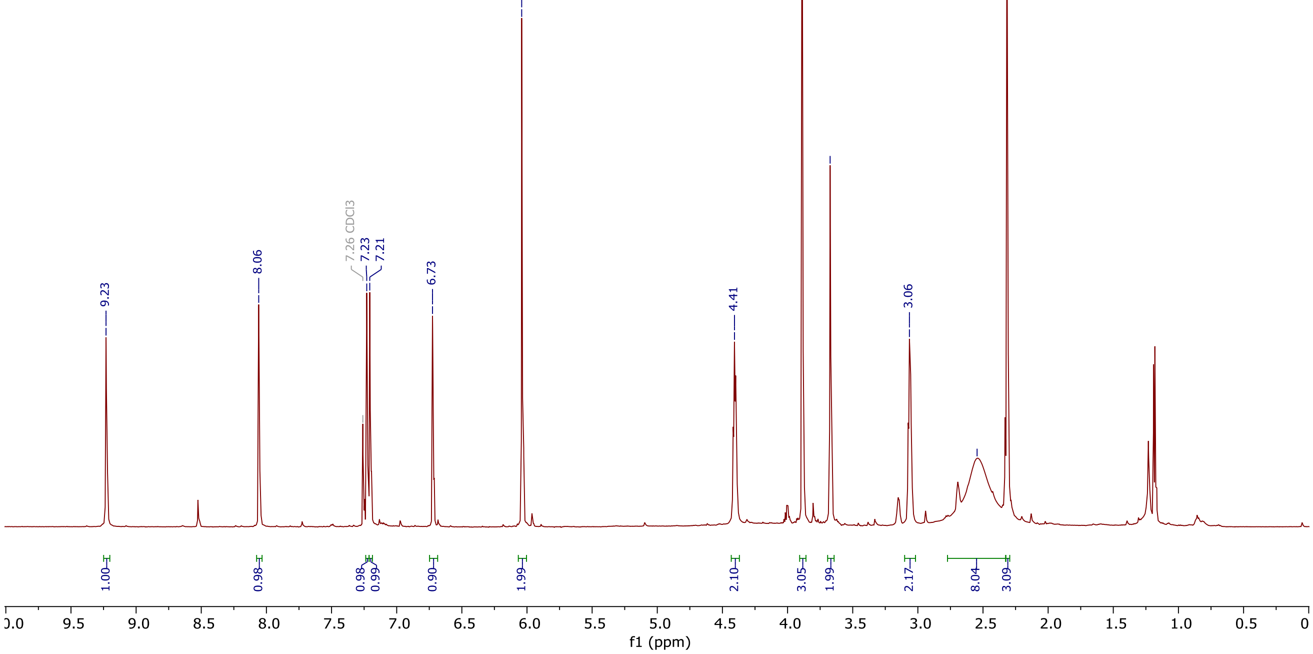
9.23
8.06
7.26 CDCl3
7.23
7.21
6.73
6.04
4.41
3.89
3.67
3.06
2.55
2.32
1.00
0.98
0.98
0.99
0.90
1.99
2.10
3.05
1.99
2.17
8.04
3.09
f1 (ppm)

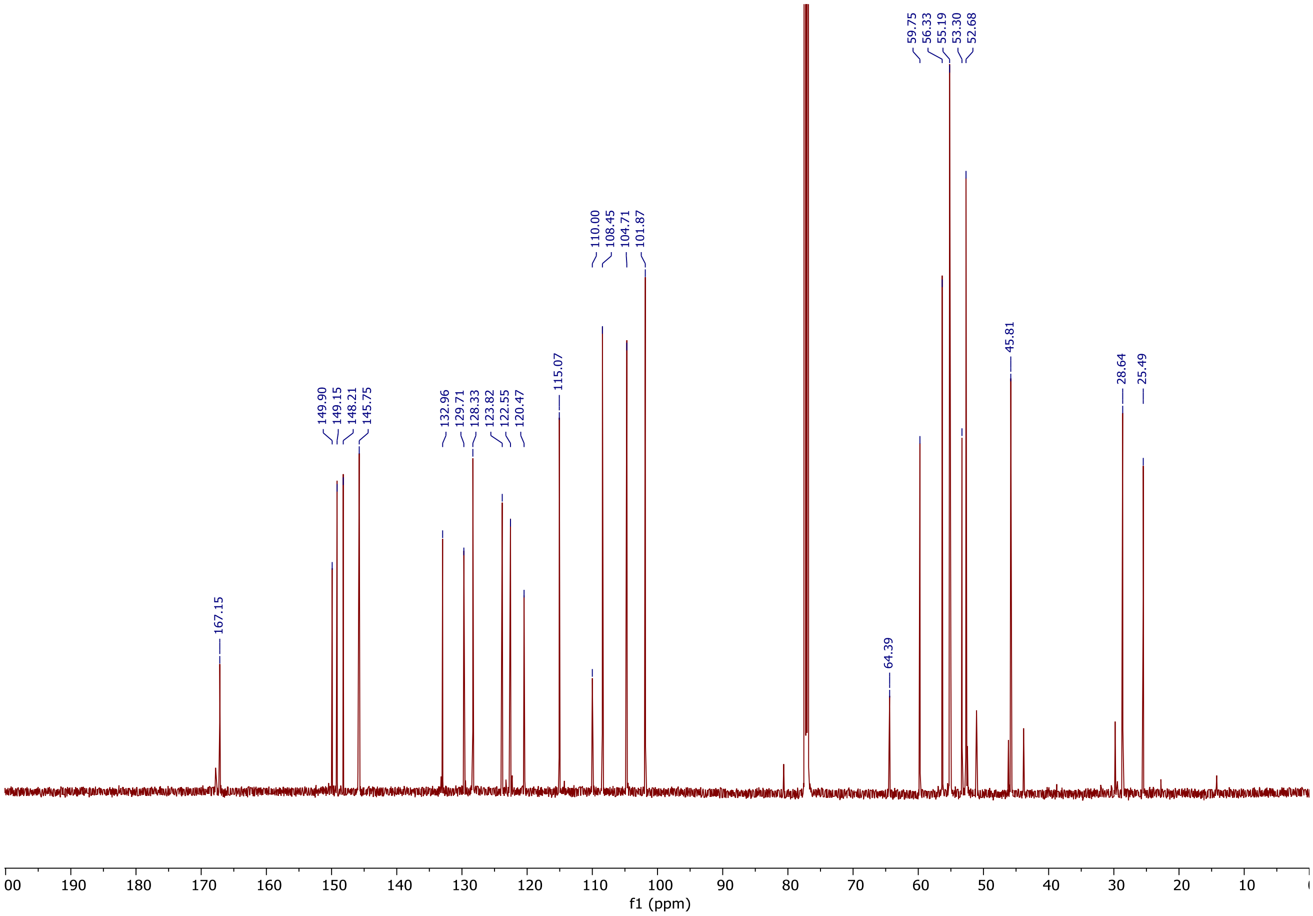
167.15
149.90
149.15
148.21
145.75
132.96
129.71
128.33
123.82
122.55
120.47
115.07
110.00
108.45
104.71
101.87
64.39
59.75
56.33
55.19
53.30
52.68
45.81
28.64
25.49
f1 (ppm)

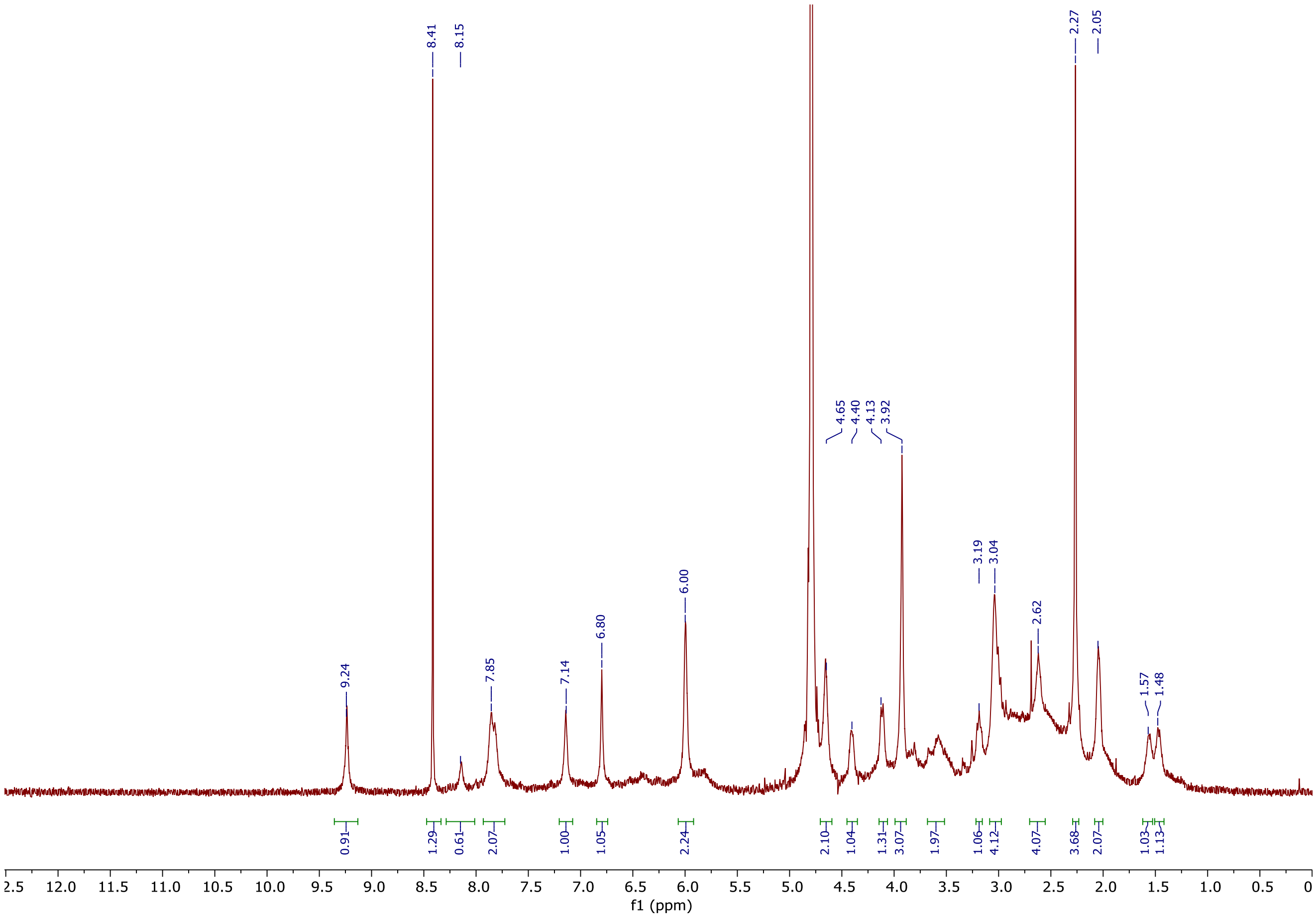
9.24
8.41
8.15
7.85
7.14
6.80
6.00
4.65
4.40
4.13
3.92
3.19
3.04
2.62
2.27
2.05
1.57
1.48
0.91
1.29
0.61
2.07
1.00
1.05
2.24
2.10
1.04
1.31
3.07
1.97
1.06
4.12
4.07
3.68
2.07
1.03
1.13
f1 (ppm)

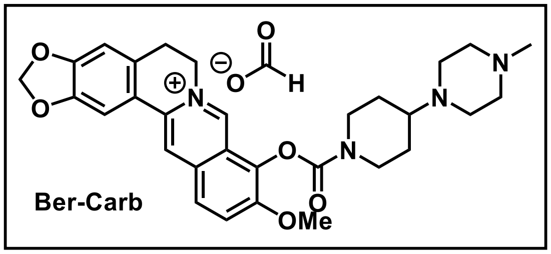
Ber-Carb
OMe
H

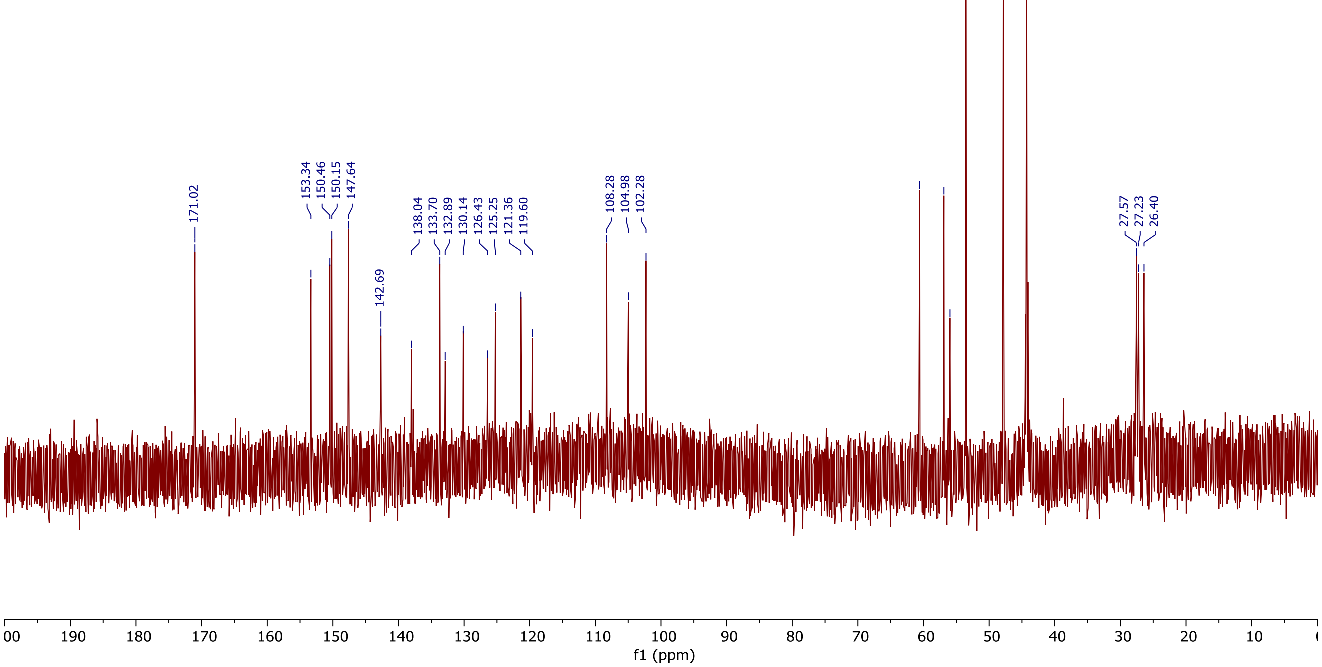
171.02
153.34
150.46
150.15
147.64
142.69
138.04
133.70
132.89
130.14
126.43
125.25
121.36
119.60
108.28
104.98
102.28
60.58
56.89
55.97
53.54
47.84
44.28
27.57
27.23
26.40
f1 (ppm)

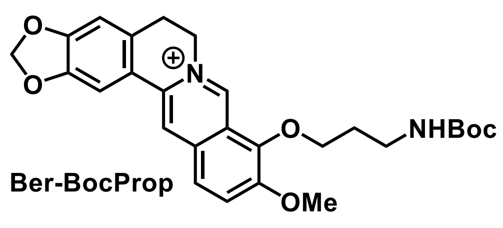
O
O
N
O
NHBoc
OMe

Ber-BocProp

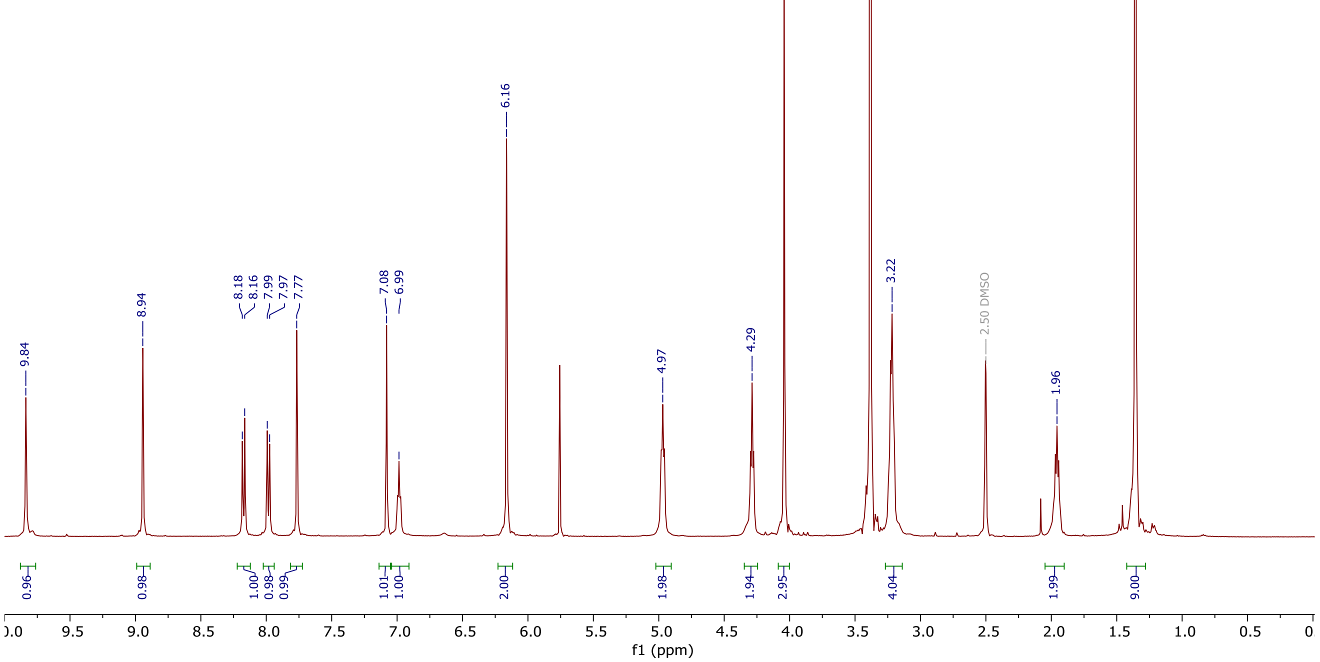
9.84
8.94
8.18
8.16
7.99
7.97
7.77
7.08
6.99
6.16
4.97
4.29
4.04
3.22
2.50 DMSO
1.96
0.96
0.98
1.00
0.98
0.99
1.01
1.00
2.00
1.98
1.94
2.95
4.04
1.99
9.00
f1 (ppm)

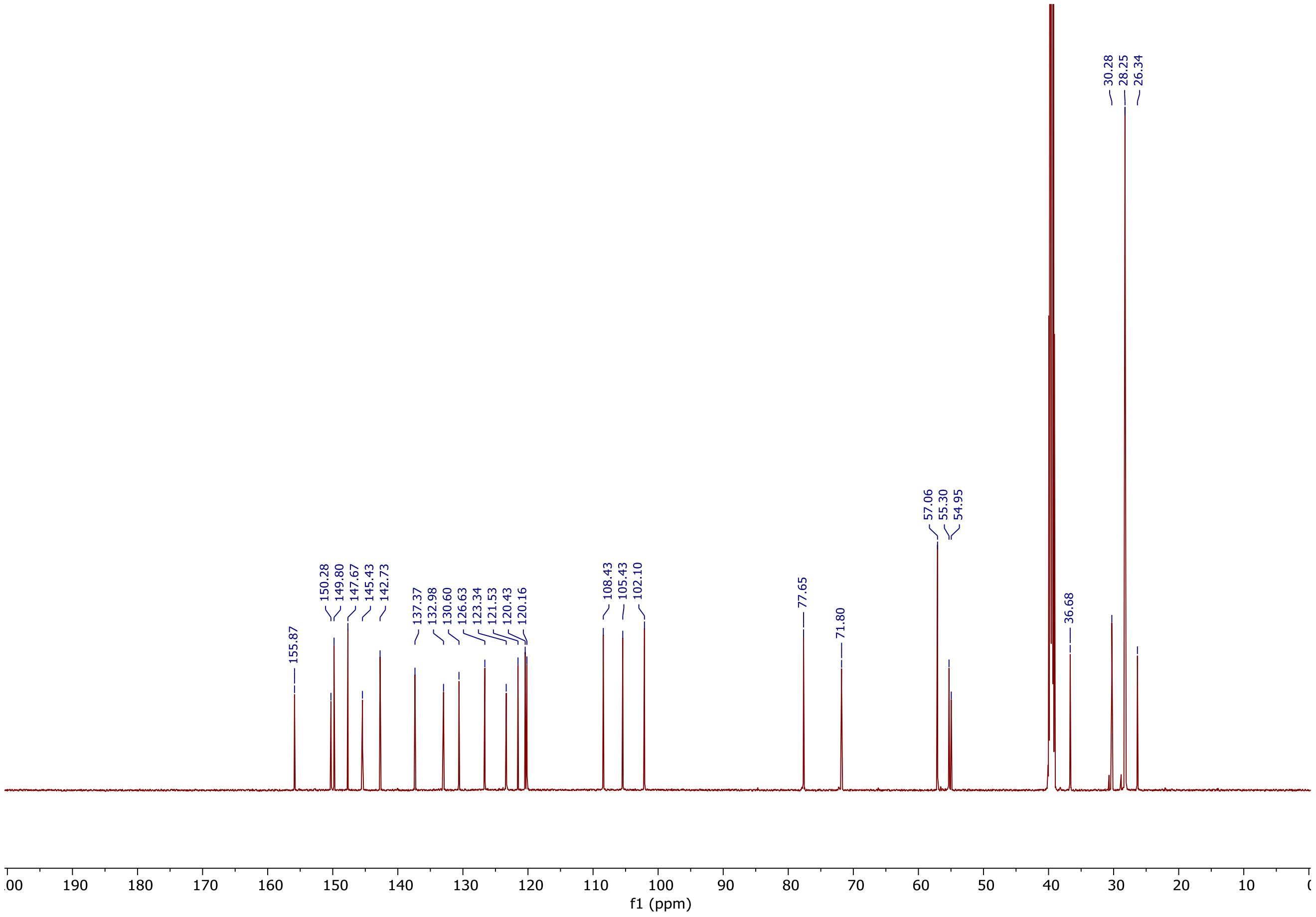
155.87
150.28
149.80
147.67
145.43
142.73
137.37
132.98
130.60
126.63
123.34
121.53
120.43
120.16
108.43
105.43
102.10
77.65
71.80
57.06
55.30
54.95
36.68
30.28
28.25
26.34
f1 (ppm)

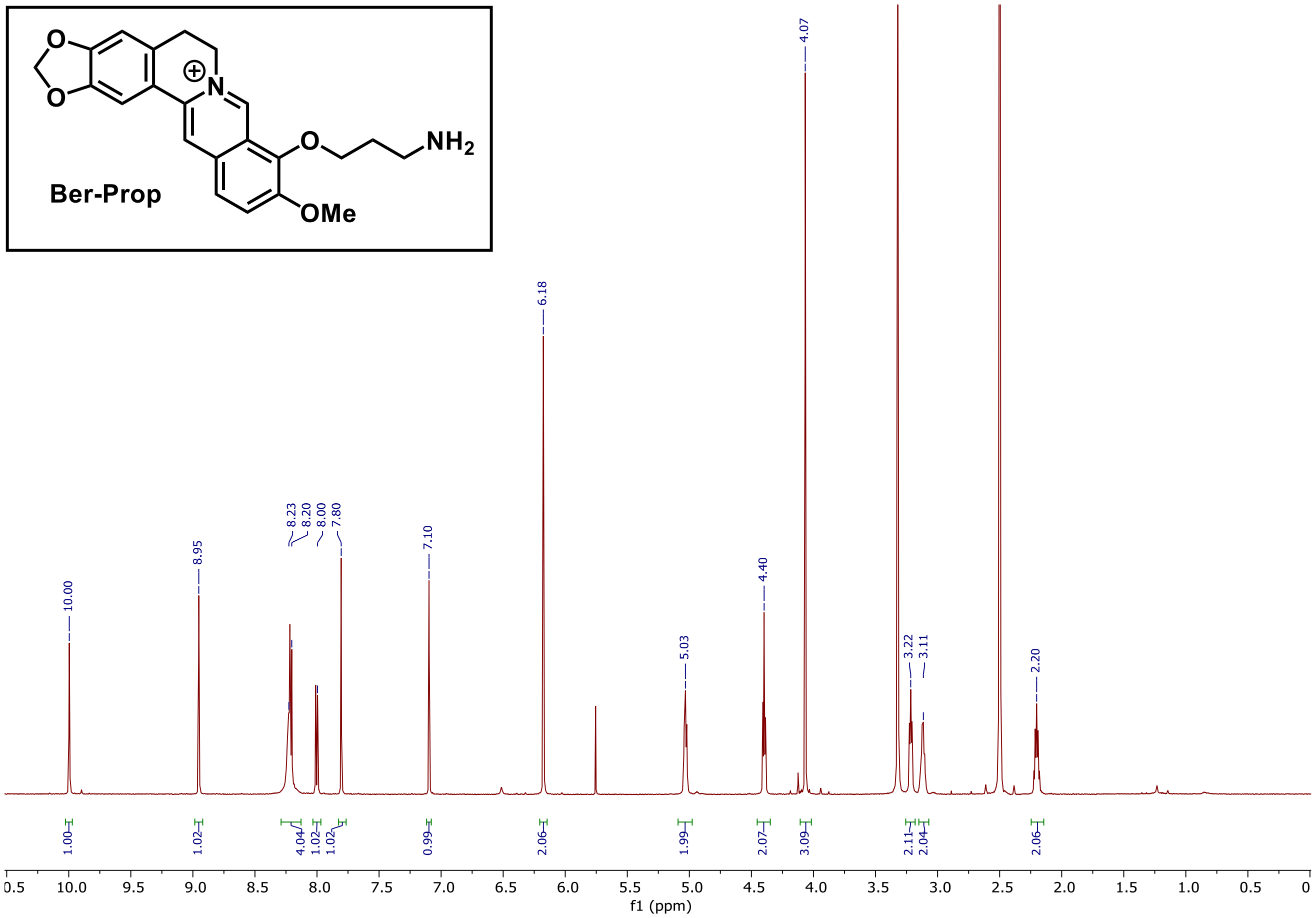
Ber-Prop
NH2
OMe
10.00
8.95
8.23
8.20
8.00
7.80
7.10
6.18
5.03
4.40
4.07
3.22
3.11
2.20
1.00
1.02
4.04
1.02
1.02
0.99
2.06
1.99
2.07
3.09
2.11
2.04
2.06
f1 (ppm)

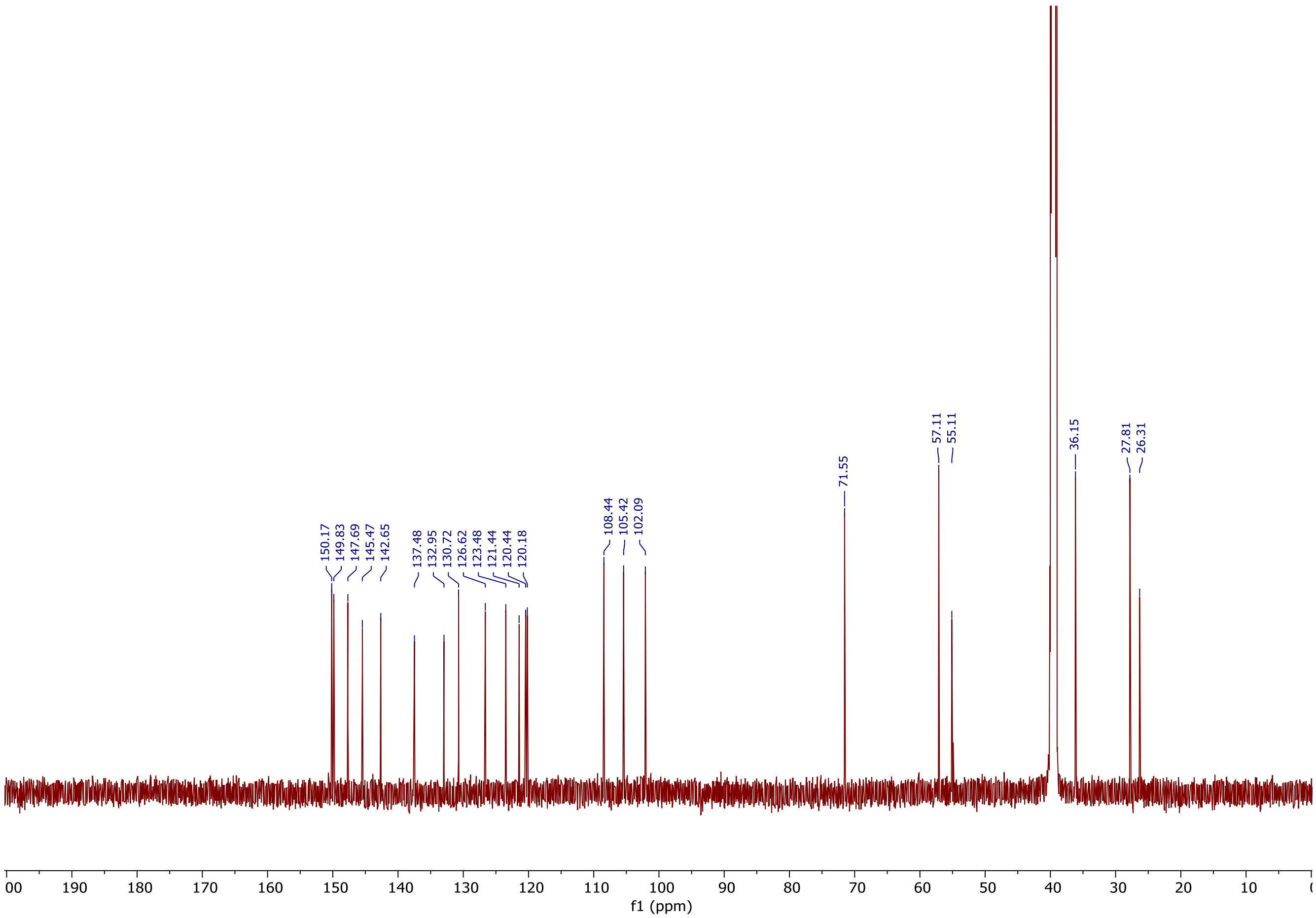
150.17
149.83
147.69
145.47
142.65
137.48
132.95
130.72
126.62
123.48
121.44
120.44
120.18
108.44
105.42
102.09
71.55
57.11
55.11
36.15
27.81
26.31
f1 (ppm)

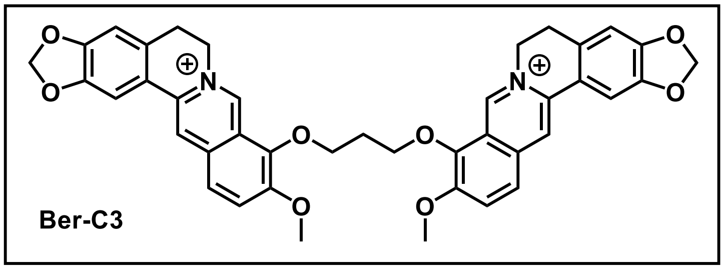
Ber-C3

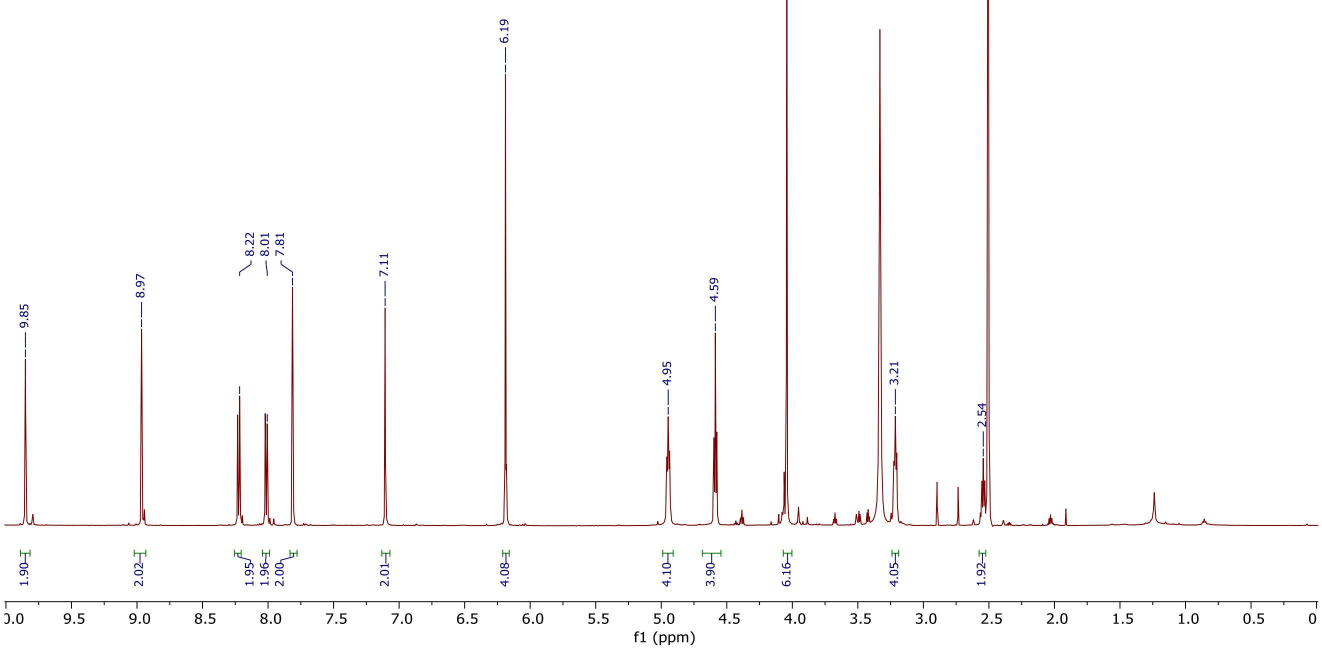
9.85
8.97
8.22
8.01
7.81
7.11
6.19
4.95
4.59
4.04
3.21
2.54
1.90
2.02
1.95
1.96
2.00
2.01
4.08
4.10
3.90
6.16
4.05
1.92
f1 (ppm)

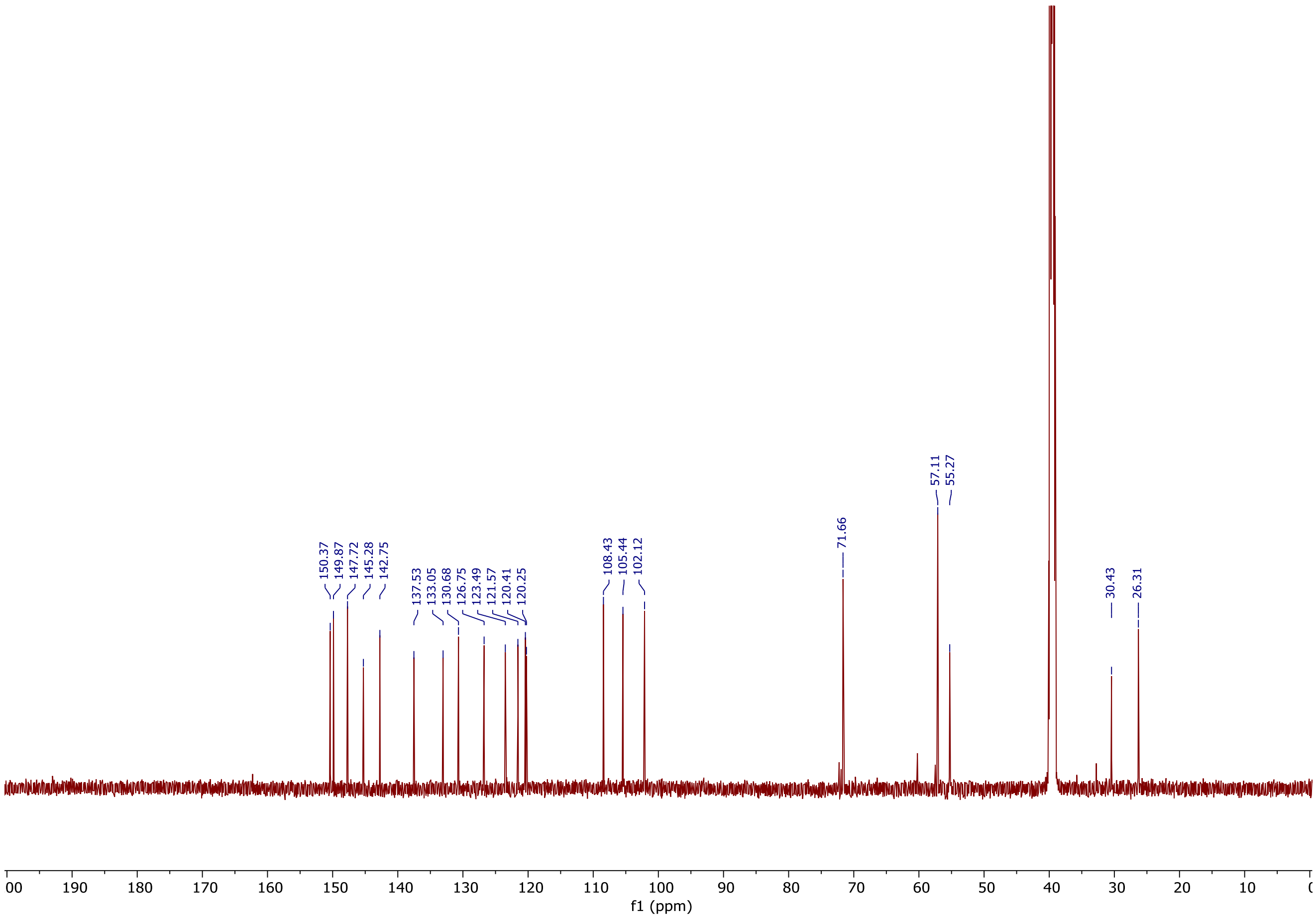
150.37
149.87
147.72
145.28
142.75
137.53
133.05
130.68
126.75
123.49
121.57
120.41
120.25
108.43
105.44
102.12
71.66
57.11
55.27
30.43
26.31
f1 (ppm)

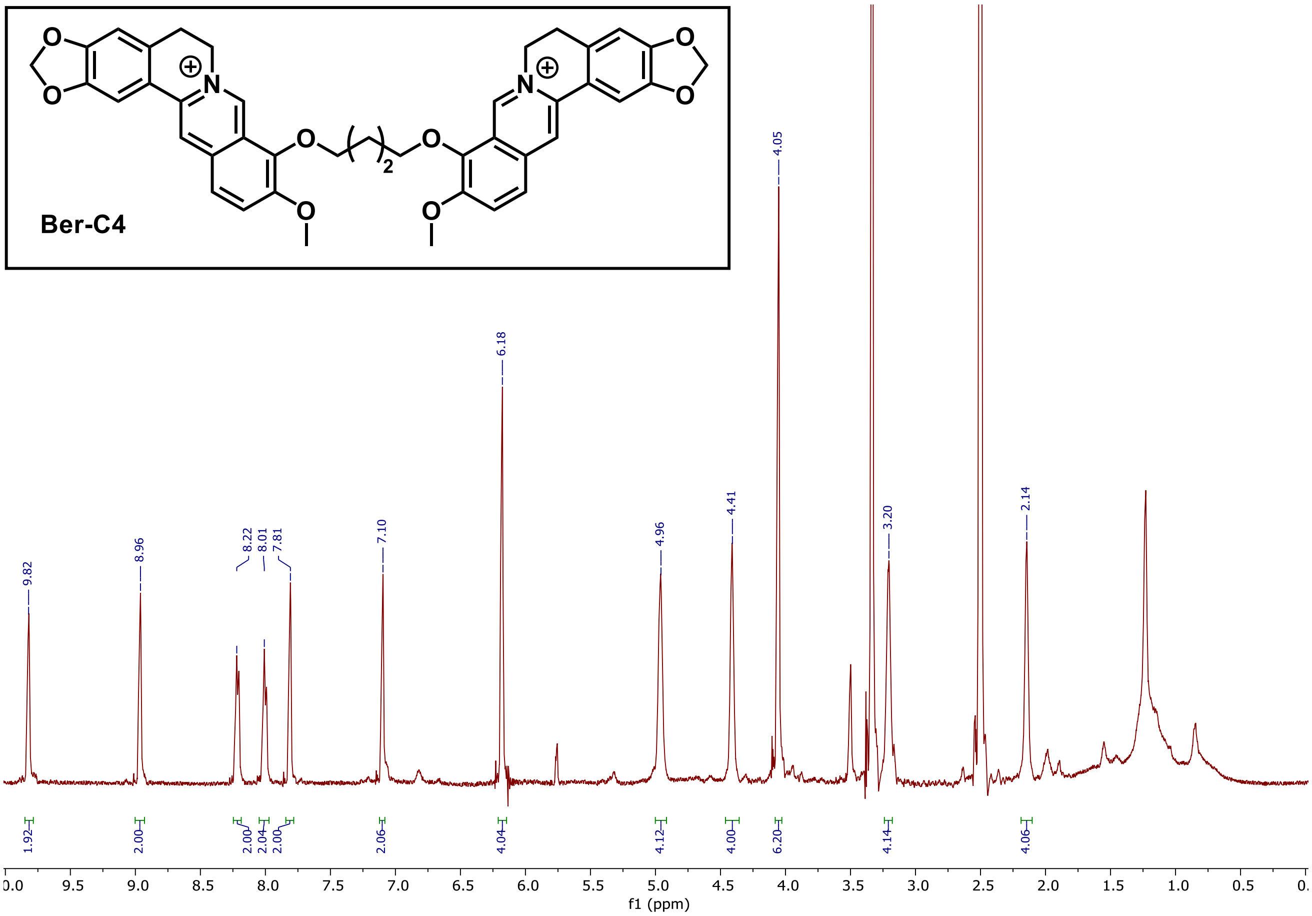
Ber-C4
9.82
8.96
8.22
8.01
7.81
7.10
6.18
4.96
4.41
4.05
3.20
2.14
1.92
2.00
2.00
2.04
2.00
2.06
4.04
4.12
4.00
6.20
4.14
4.06
f1 (ppm)

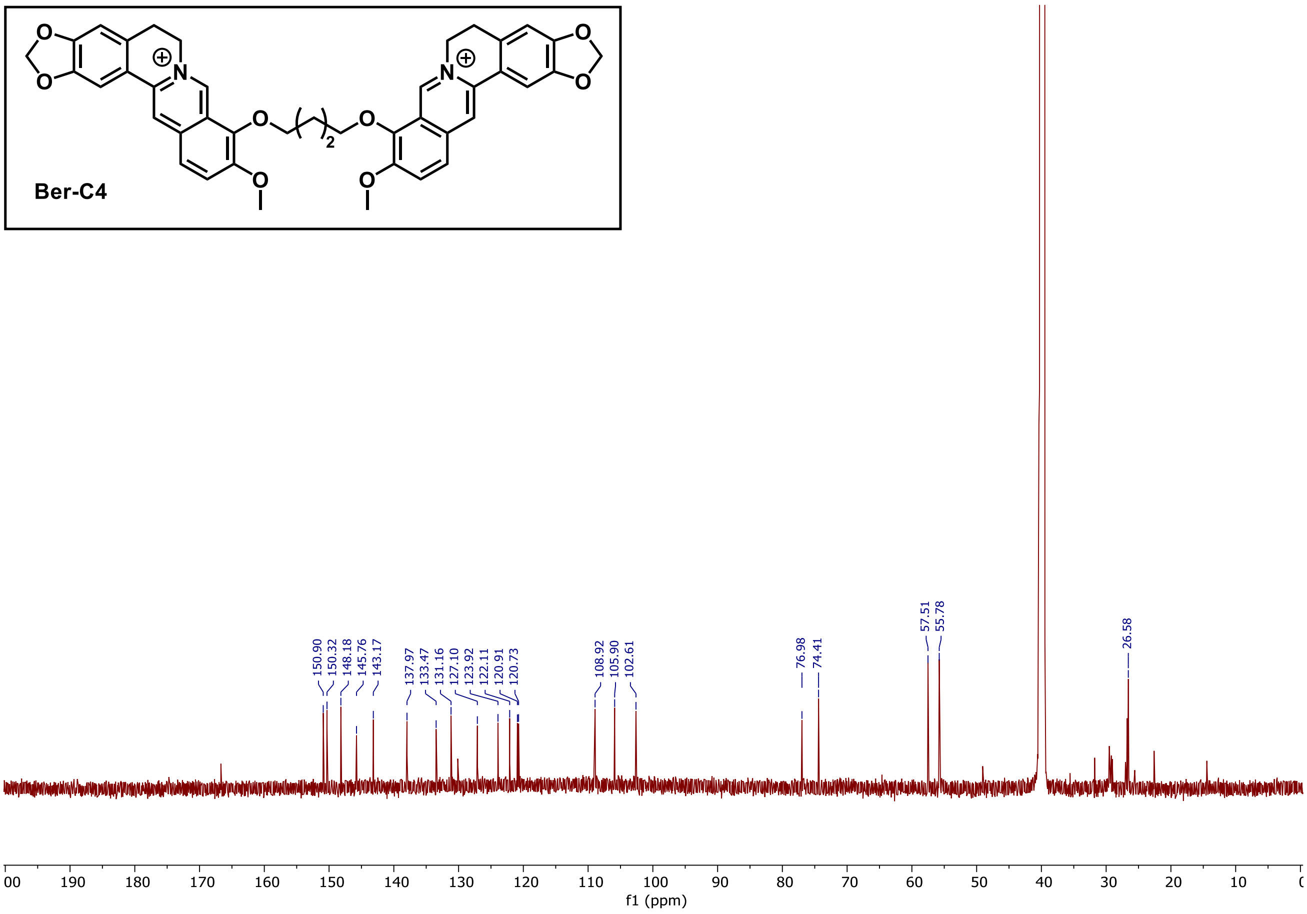
Ber-C4
150.90
150.32
148.18
145.76
143.17
137.97
133.47
131.16
127.10
123.92
122.11
120.91
120.73
108.92
105.90
102.61
76.98
74.41
57.51
55.78
26.58
f1 (ppm)

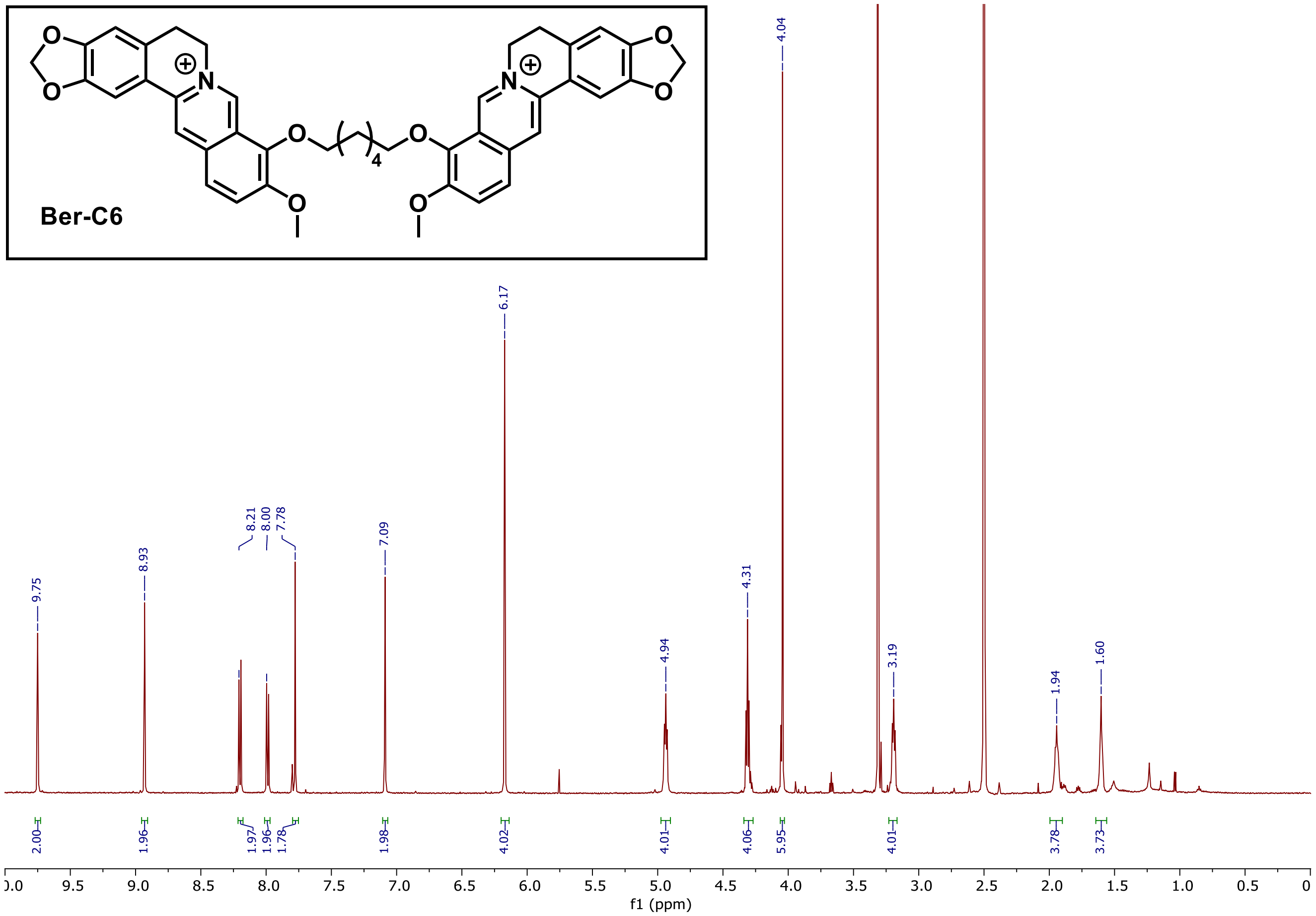
Ber-C6
9.75
8.93
8.21
8.00
7.78
7.09
6.17
4.94
4.31
4.04
3.19
1.94
1.60
2.00
1.96
1.97
1.96
1.78
1.98
4.02
4.01
4.06
5.95
4.01
3.78
3.73
f1 (ppm)

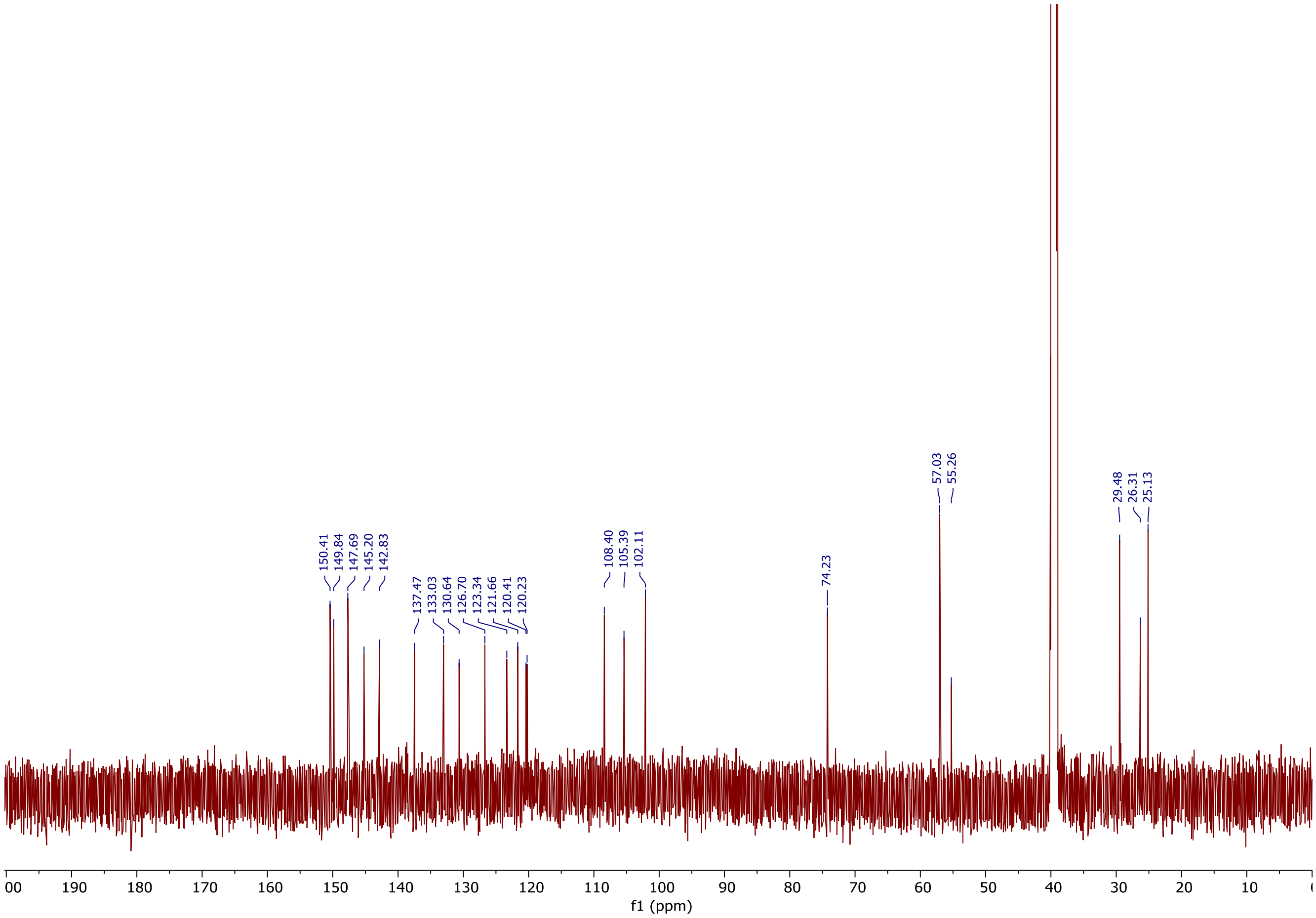
150.41
149.84
147.69
145.20
142.83
137.47
133.03
130.64
126.70
123.34
121.66
120.41
120.23
108.40
105.39
102.11
74.23
57.03
55.26
29.48
26.31
25.13
f1 (ppm)

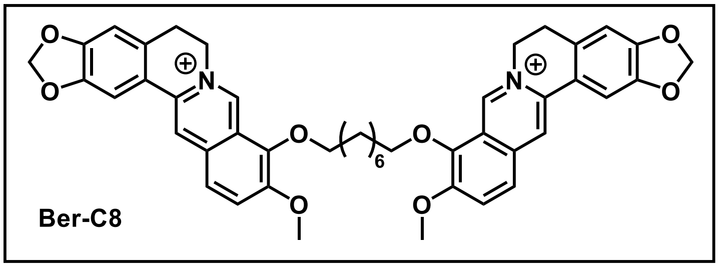

Ber-C8

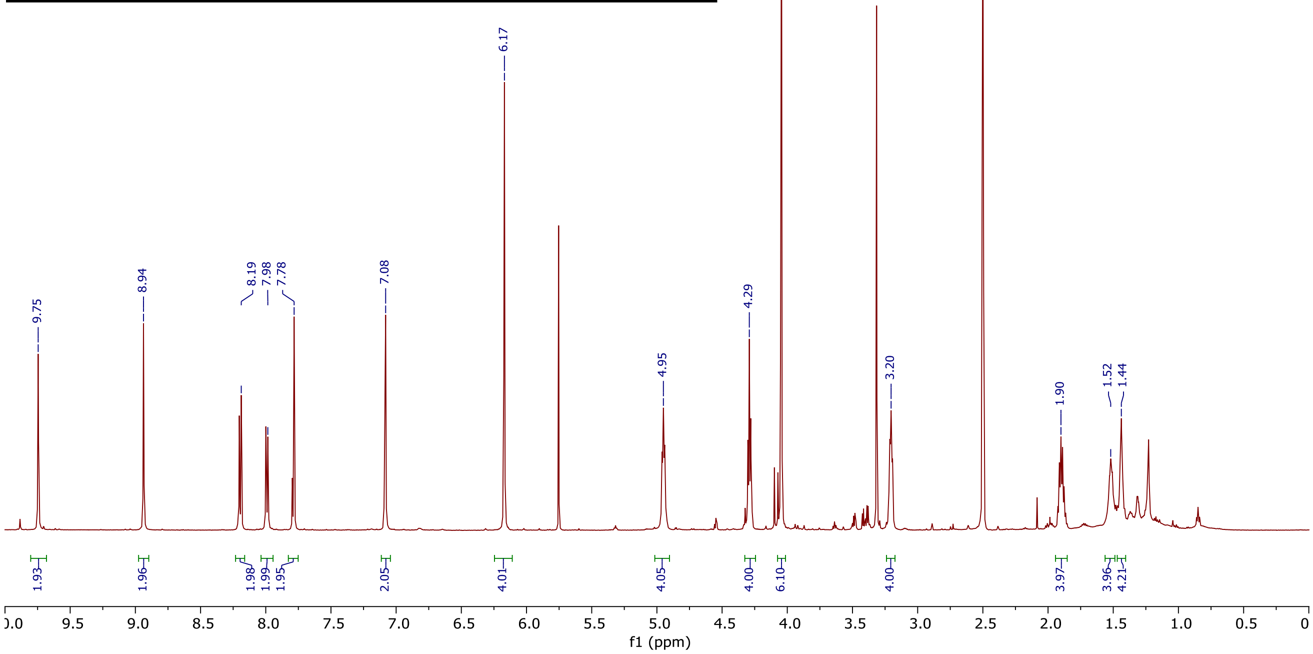
9.75
8.94
8.19
7.98
7.78
7.08
6.17
4.95
4.29
4.04
3.20
2.50 DMSO
1.90
1.52
1.44
1.93
1.96
1.98
1.99
1.95
2.05
4.01
4.05
4.00
6.10
4.00
3.97
3.96
4.21
f1 (ppm)

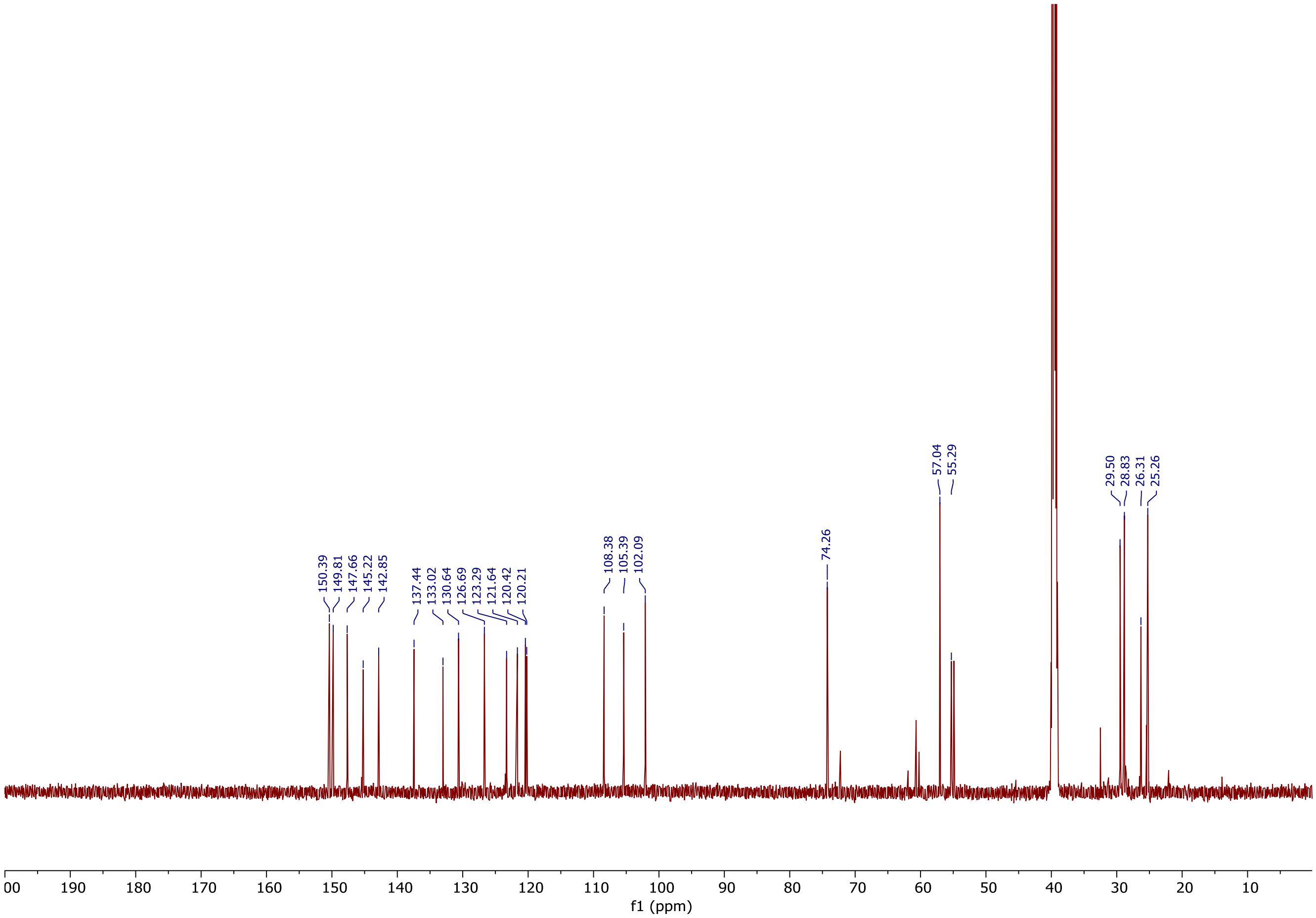
150.39
149.81
147.66
145.22
142.85
137.44
133.02
130.64
126.69
123.29
121.64
120.42
120.21
108.38
105.39
102.09
74.26
57.04
55.29
29.50
28.83
26.31
25.26
f1 (ppm)

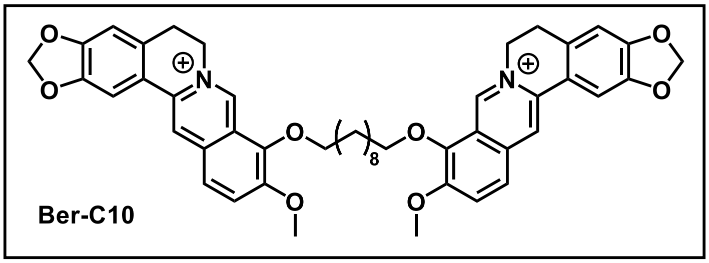
Ber-C10
8

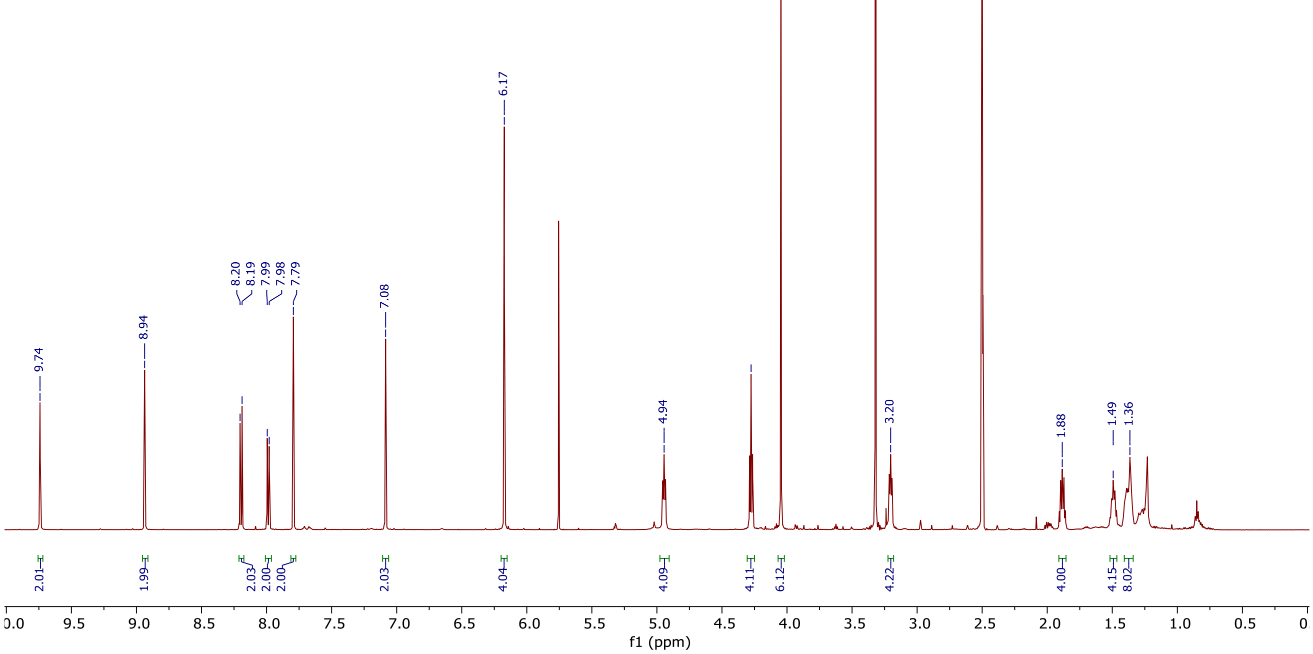
9.74
8.94
8.20
8.19
7.99
7.98
7.79
7.08
6.17
4.94
4.28
4.05
3.20
1.88
1.49
1.36
2.01
1.99
2.03
2.00
2.00
2.03
4.04
4.09
4.11
6.12
4.22
4.00
4.15
8.02
f1 (ppm)

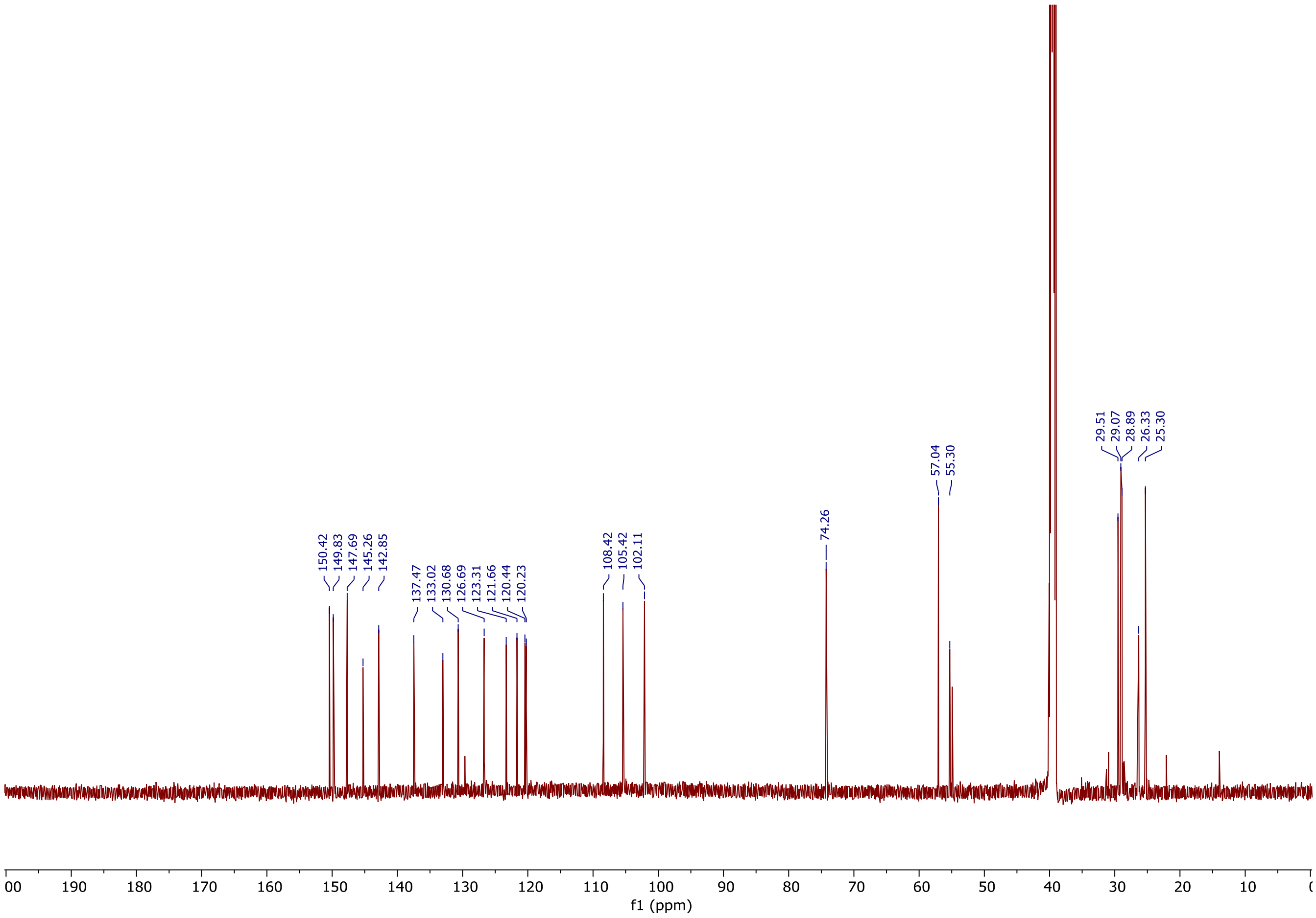
150.42
149.83
147.69
145.26
142.85
137.47
133.02
130.68
126.69
123.31
121.66
120.44
120.23
108.42
105.42
102.11
74.26
57.04
55.30
29.51
29.07
28.89
26.33
25.30
f1 (ppm)

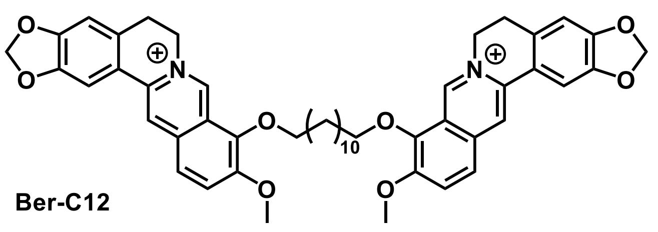

O
O
N
O
O
10
O
O
O
O
N
O


Ber-C12

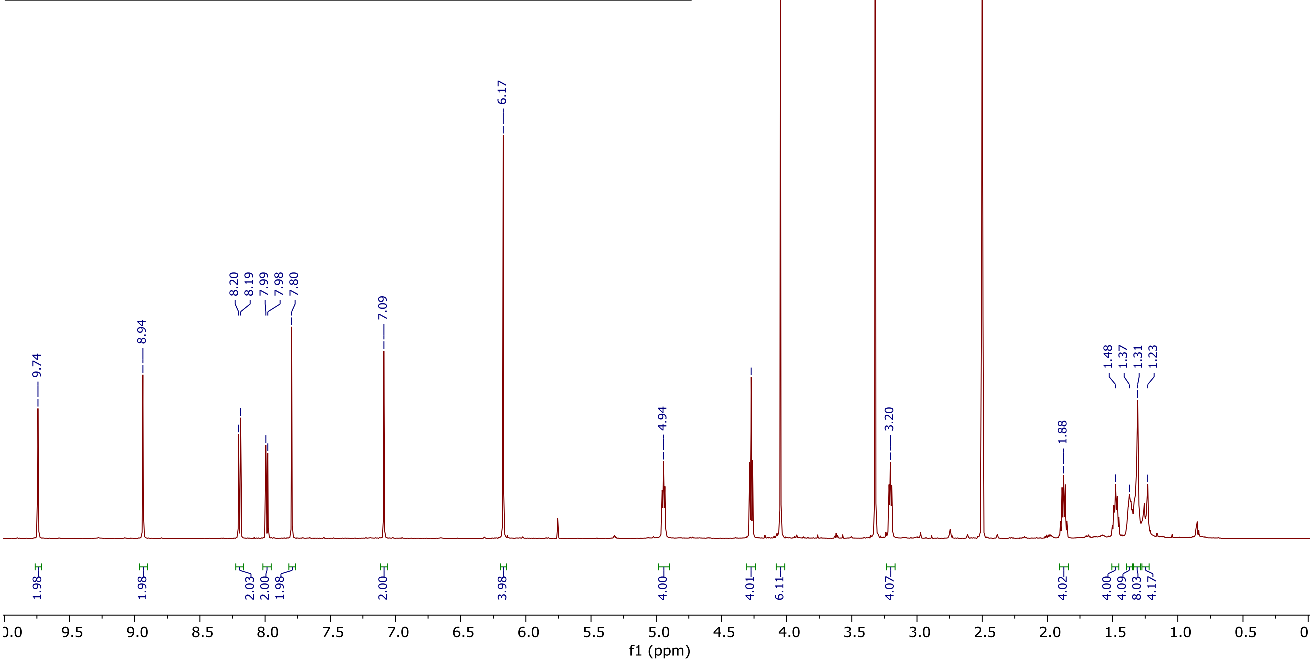

9.74
8.94
8.20
8.19
7.99
7.98
7.80
7.09
6.17
4.94
4.27
4.05
3.20
2.50 DMSO
1.88
1.48
1.37
1.31
1.23
1.98
1.98
2.03
2.00
1.98
2.00
3.98
4.00
4.01
6.11
4.07
4.02
4.00
4.09
8.03
4.17
f1 (ppm)


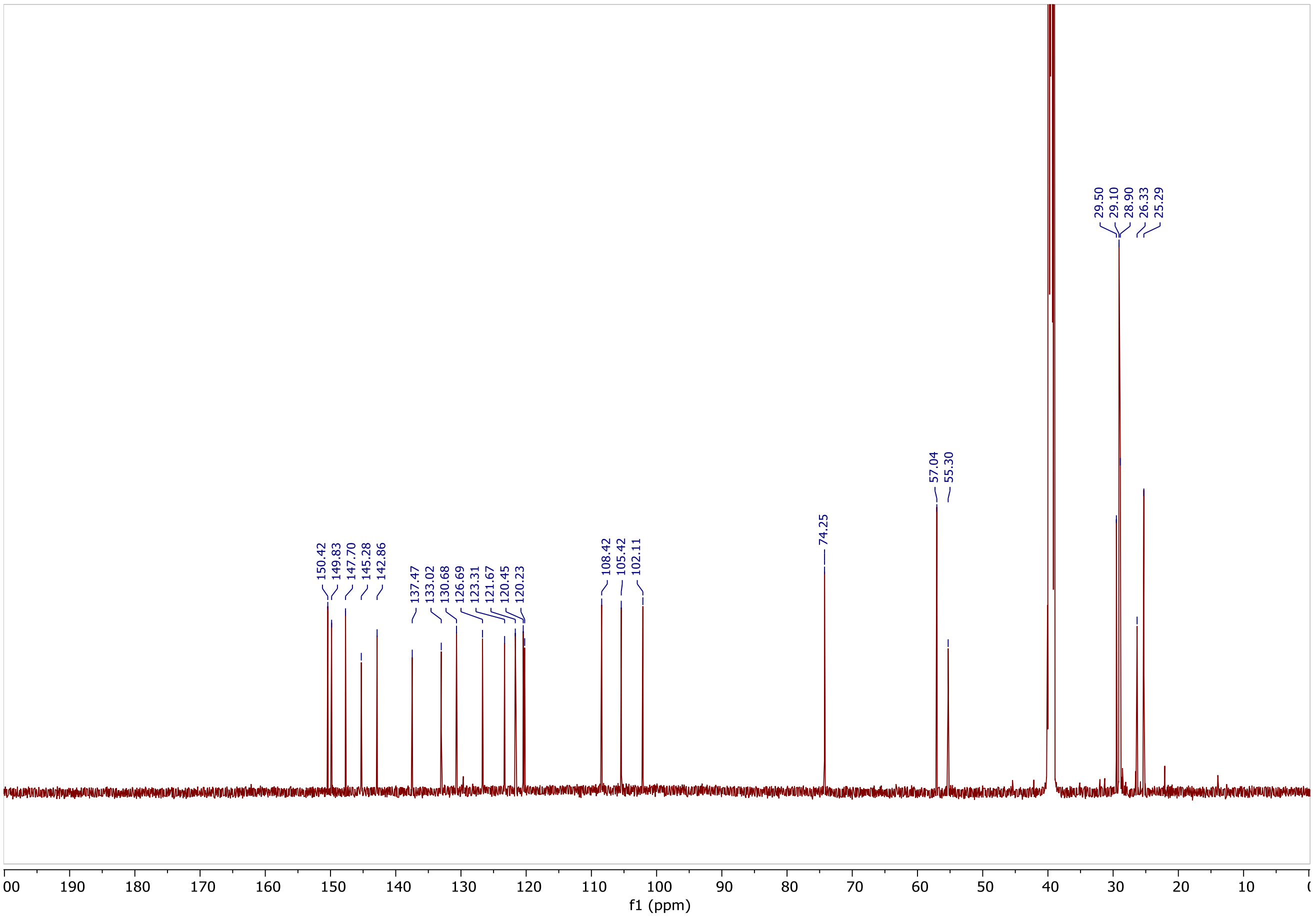

150.42
149.83
147.70
145.28
142.86
137.47
133.02
130.68
126.69
123.31
121.67
120.45
120.23
108.42
105.42
102.11
74.25
57.04
55.30
29.50
29.10
28.90
26.33
25.29
f1 (ppm)

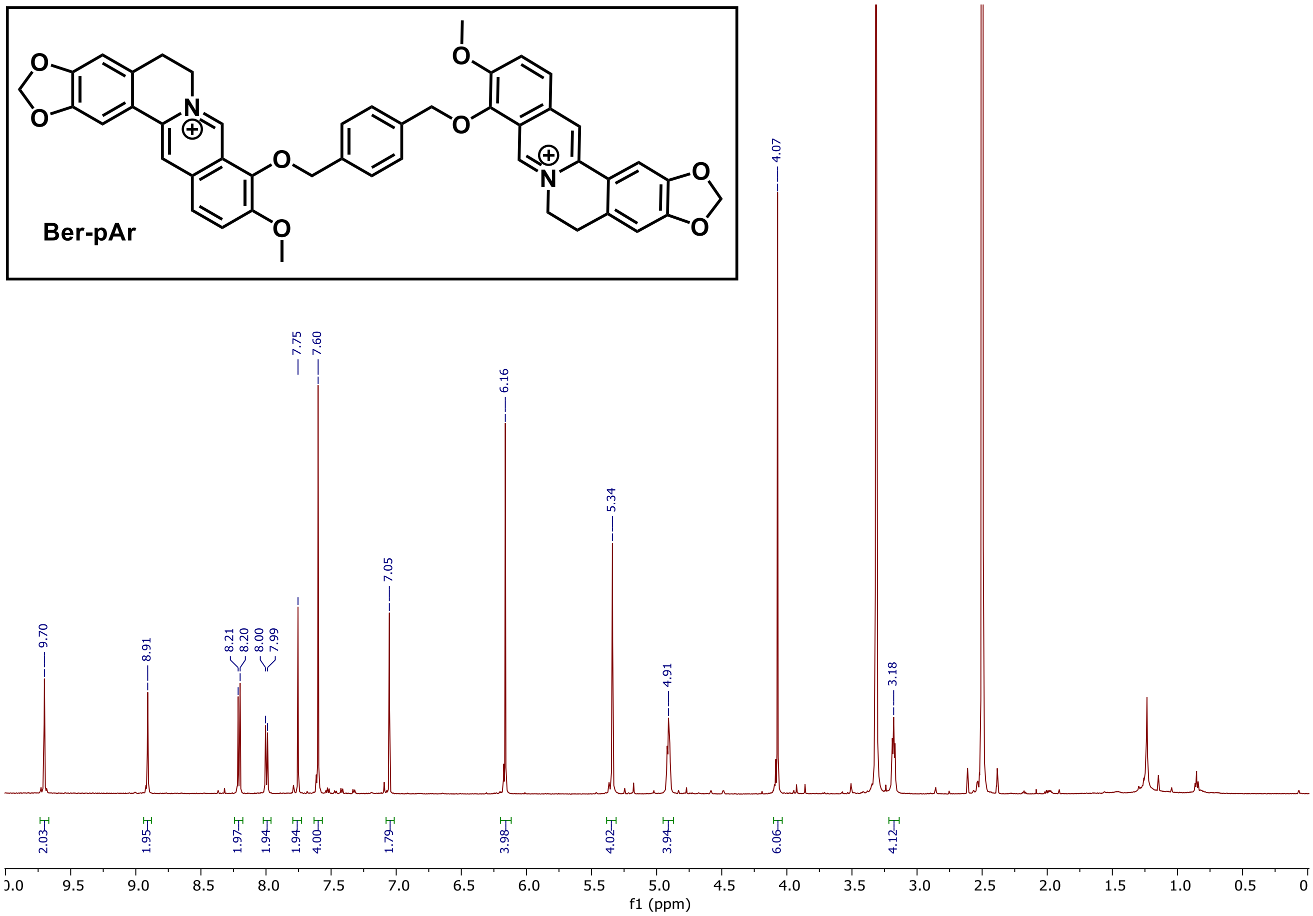
Ber-pAr
9.70
8.91
8.21
8.20
8.00
7.99
7.75
7.60
7.05
6.16
5.34
4.91
4.07
3.18
2.03
1.95
1.97
1.94
1.94
4.00
1.79
3.98
4.02
3.94
6.06
4.12
f1 (ppm)

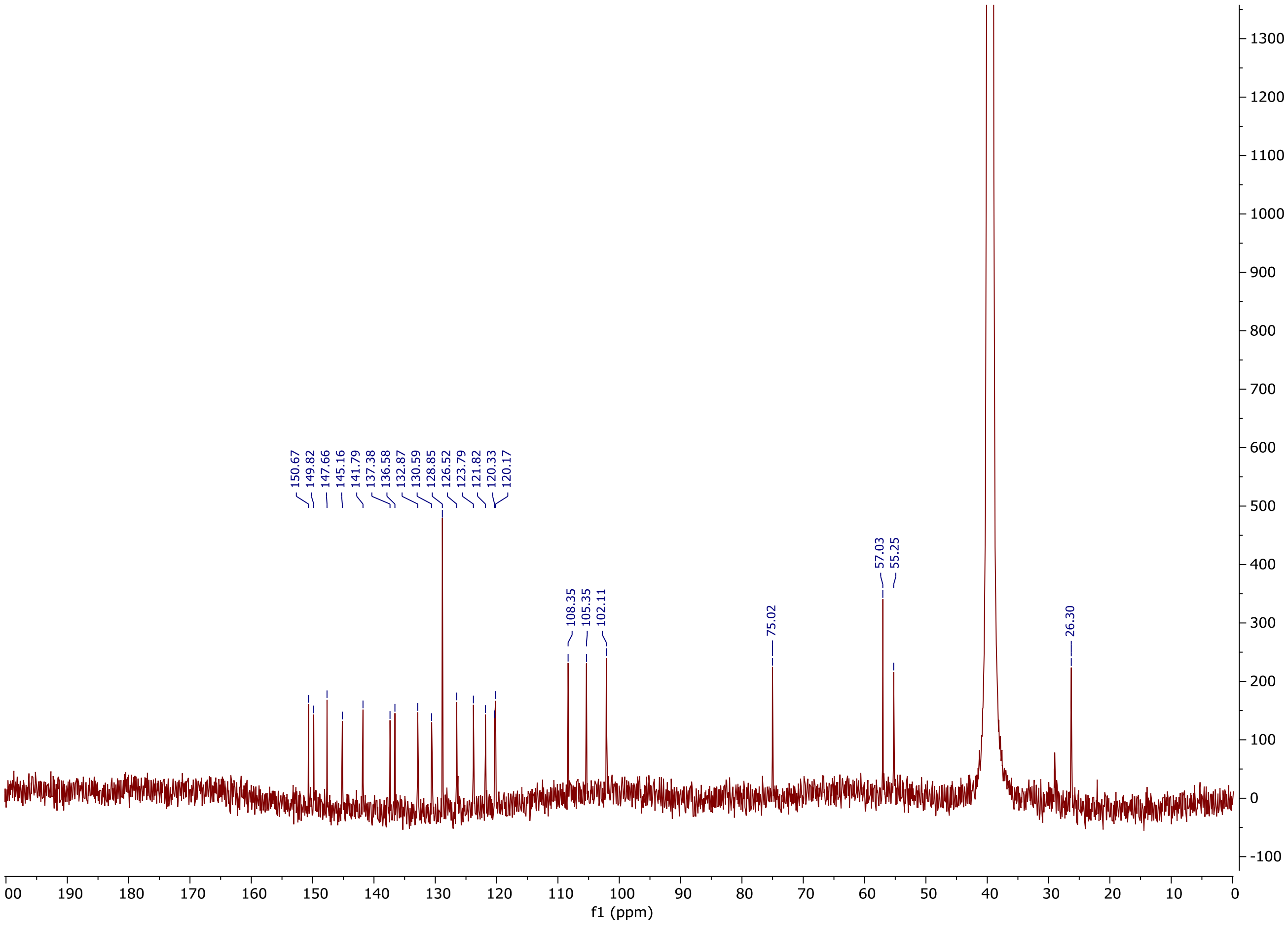
150.67
149.82
147.66
145.16
141.79
137.38
136.58
132.87
130.59
128.85
126.52
123.79
121.82
120.33
120.17
108.35
105.35
102.11
75.02
57.03
55.25
26.30
f1 (ppm)

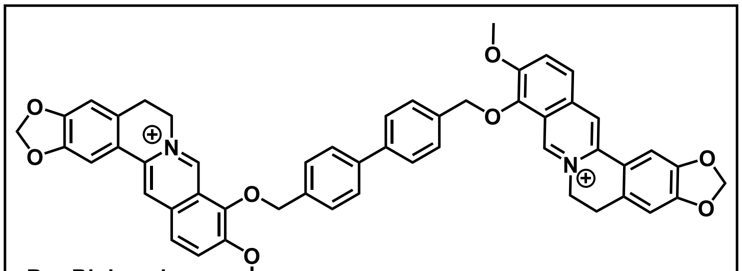

**Ber-Biphenyl**

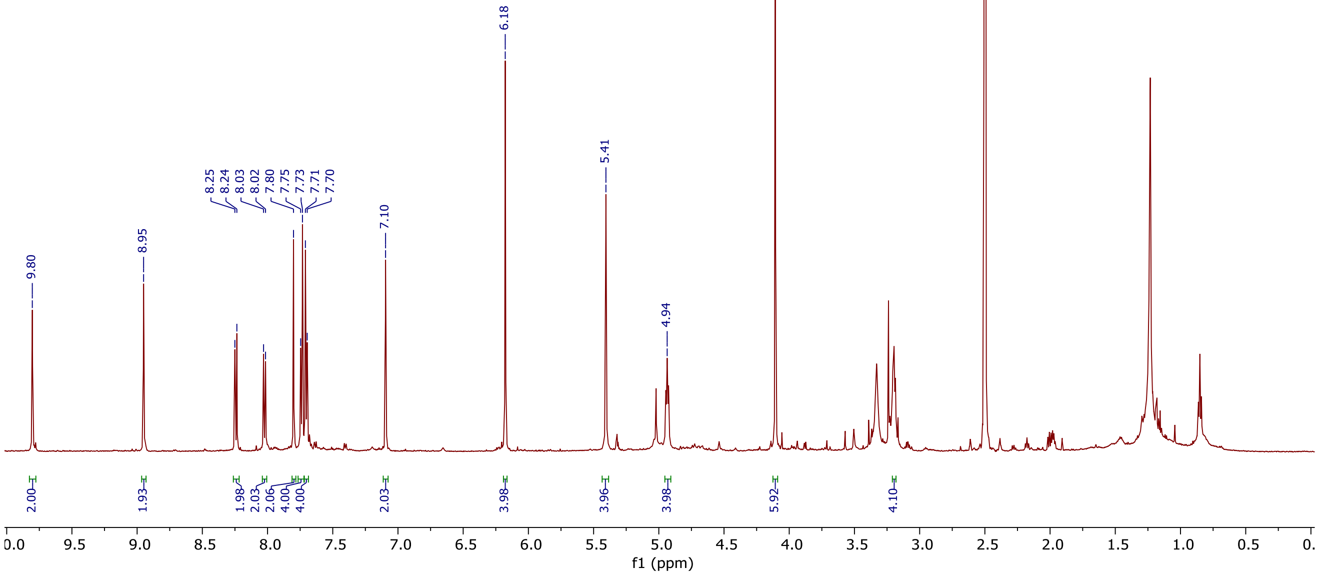


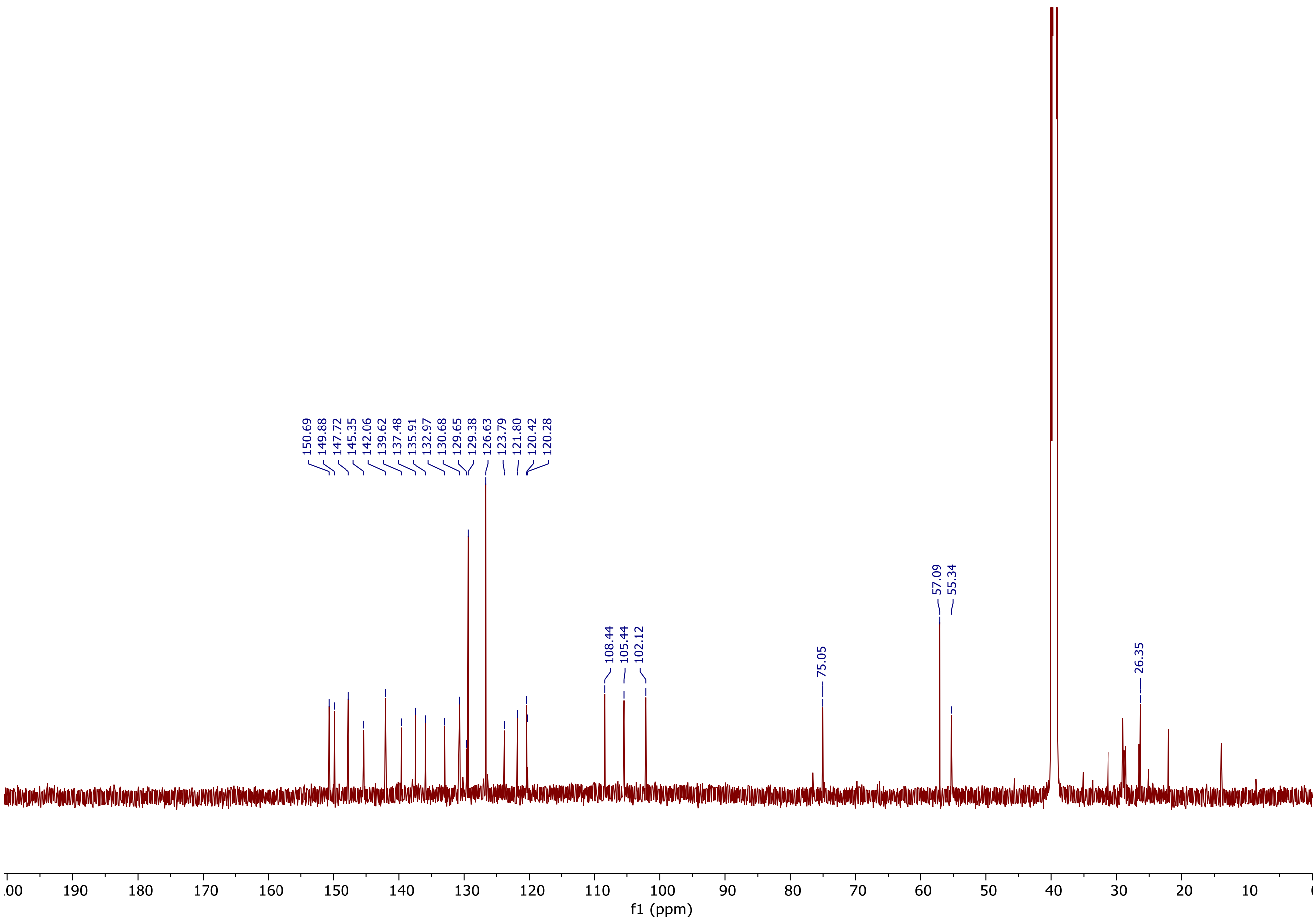

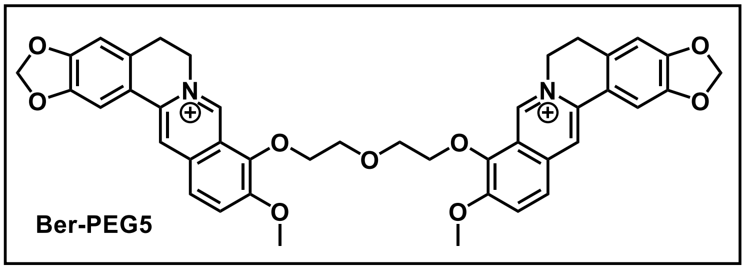
O
O
N
O
O
O
O
O
N
O
O

Ber-PEG5

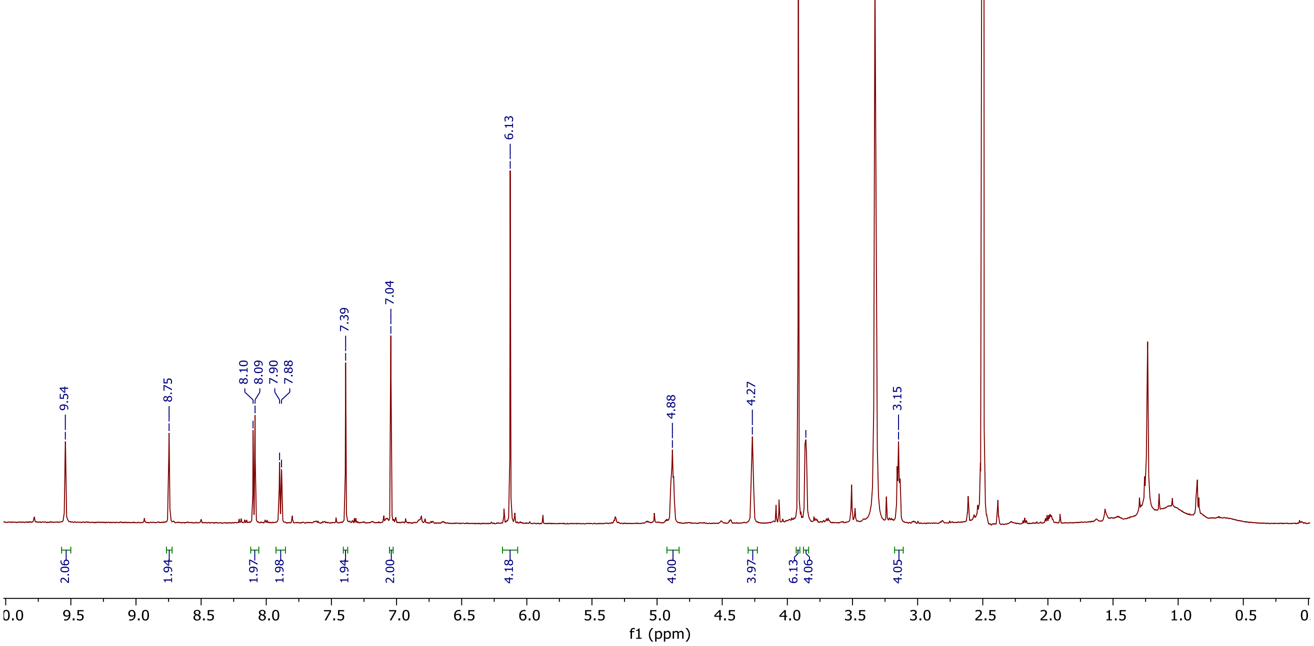
9.54
8.75
8.10
8.09
7.90
7.88
7.39
7.04
6.13
4.88
4.27
3.92
3.86
3.15
2.06
1.94
1.97
1.98
1.94
2.00
4.18
4.00
3.97
6.13
4.06
4.05
f1 (ppm)

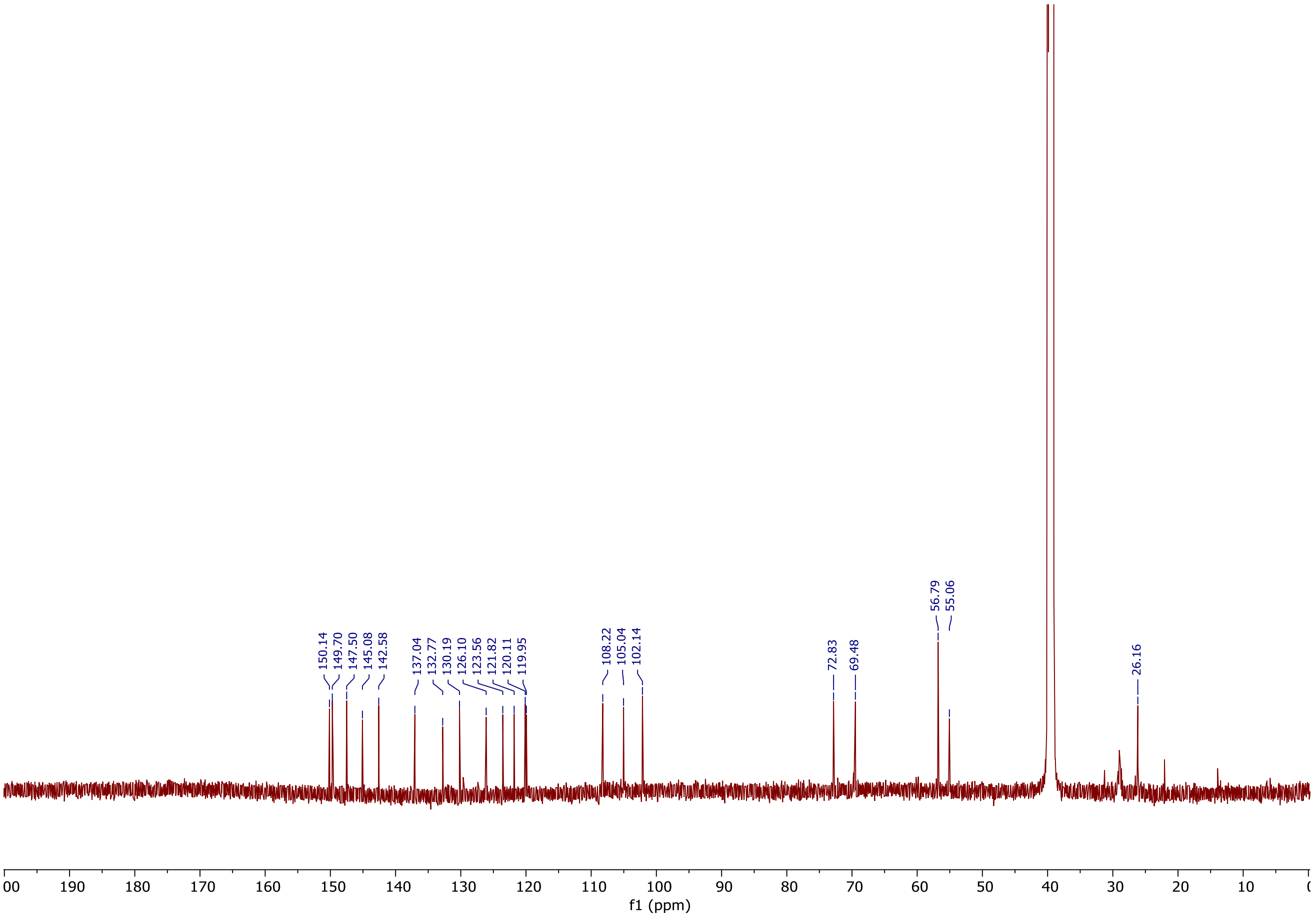
150.14
149.70
147.50
145.08
142.58
137.04
132.77
130.19
126.10
123.56
121.82
120.11
119.95
108.22
105.04
102.14
72.83
69.48
56.79
55.06
26.16
f1 (ppm)

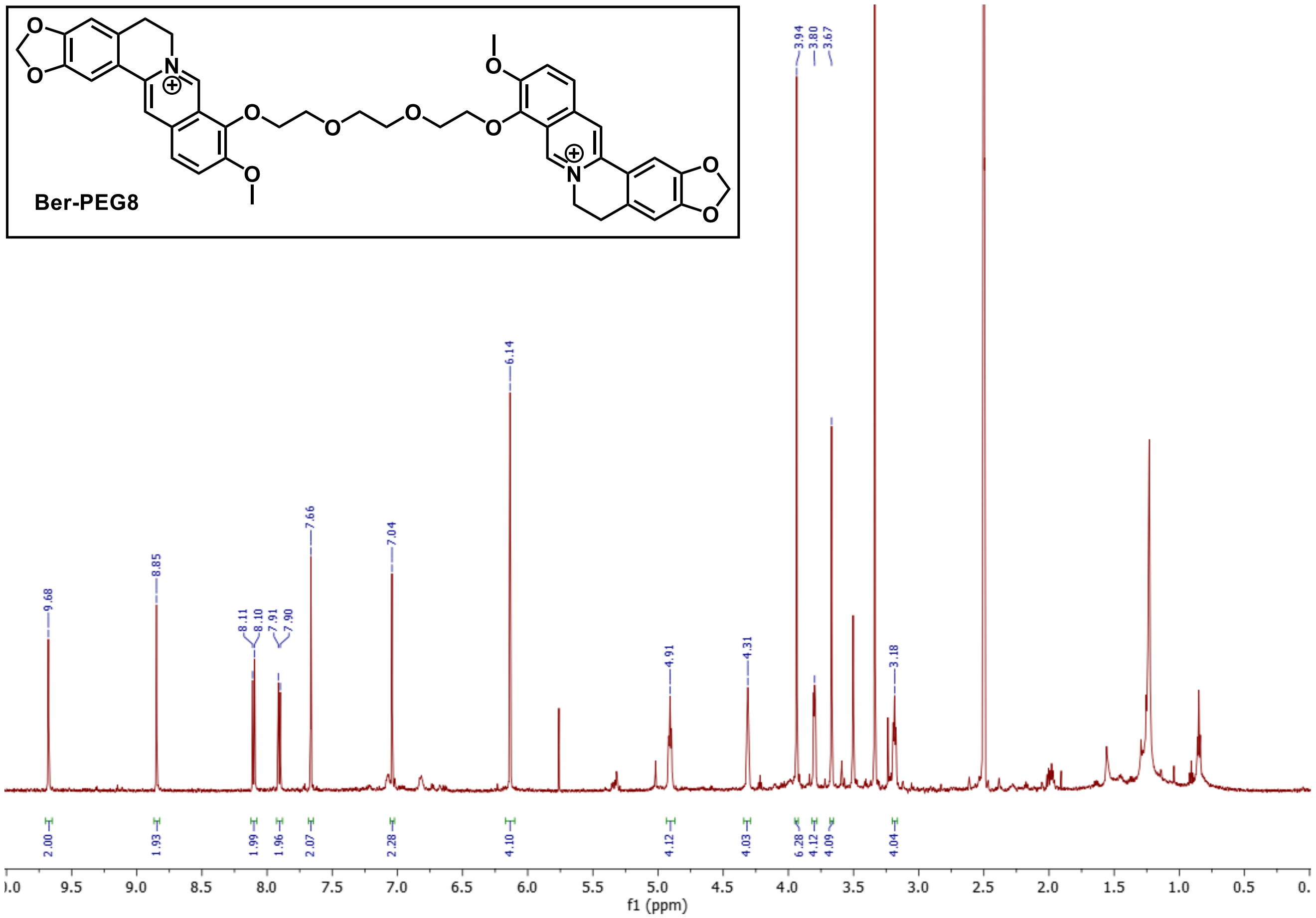
Ber-PEG8
9.68
8.85
8.11
8.10
7.91
7.90
7.66
7.04
6.14
4.91
4.31
3.94
3.80
3.67
3.18
2.00
1.93
1.99
1.96
2.07
2.28
4.10
4.12
4.03
6.28
4.12
4.09
4.04
f1 (ppm)

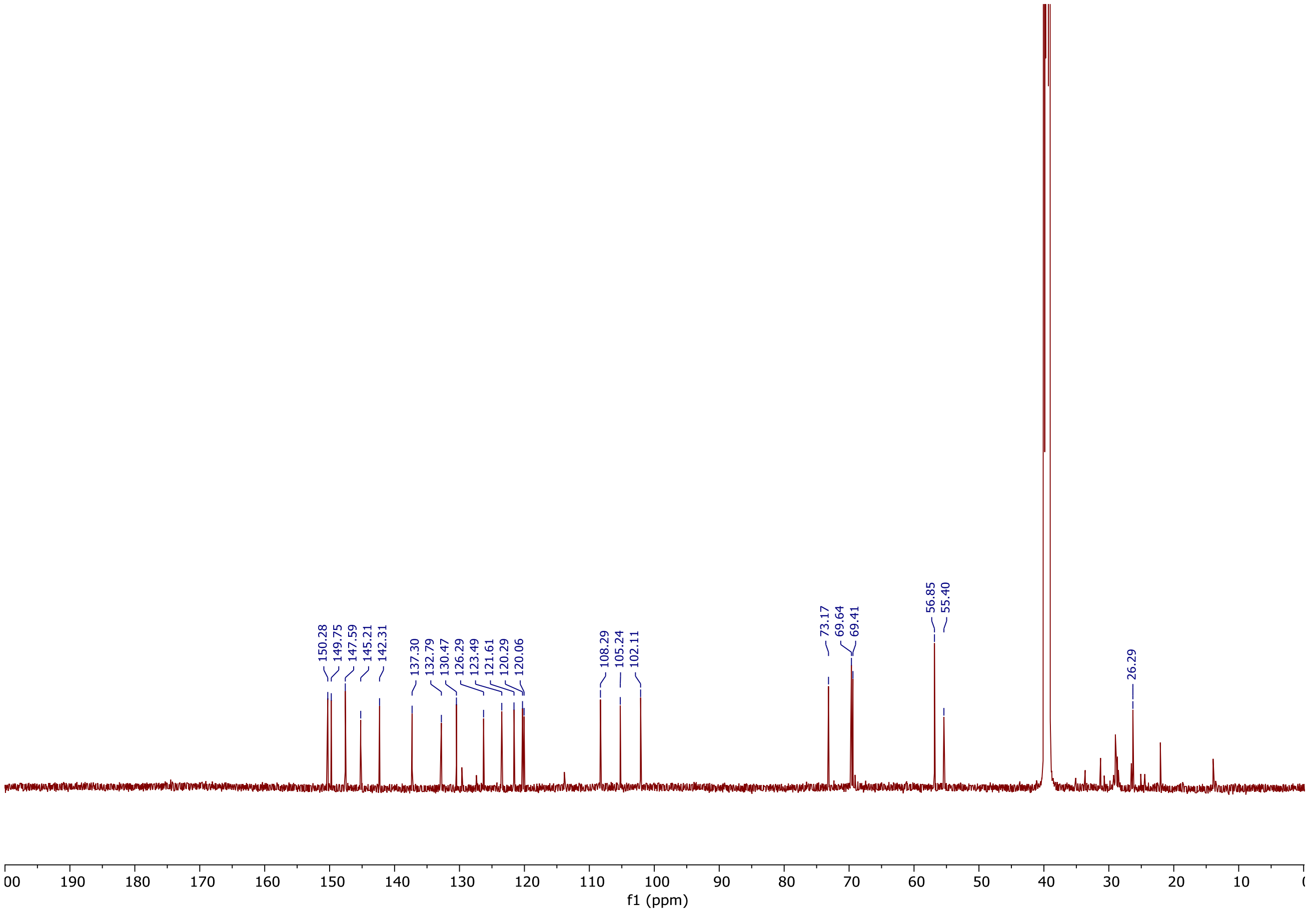
150.28
149.75
147.59
145.21
142.31
137.30
132.79
130.47
126.29
123.49
121.61
120.29
120.06
108.29
105.24
102.11
73.17
69.64
69.41
56.85
55.40
26.29
f1 (ppm)
